# Ancestors of Arylmalonate Decarboxylase show increased Activity, Stability and Stereoselectivity

**DOI:** 10.64898/2026.01.14.699310

**Authors:** Elske van der Pol, Jan Gerstenberger, Xenia Georgiadou, Gyula Hoffka, Lisa-Marie Krammer, Klaus Schliep, Colin Schür, Shina Caroline Lynn Kamerlin, Selin Kara, Robert Kourist

## Abstract

Bacterial aryl malonate decarboxylase (AMDase) generates a wide spectrum of α-chiral carboxylic acids in outstanding optical purity, including several non-steroidal anti-inflammatory drugs and chiral building blocks. However, extant AMDase from *Bordetella bronchiseptica* (*Bb*AMDase) and related enzymes of the same family have three main limitations: (i) low operational and thermal stability, (ii) limited substrate spectrum regarding the size of the smaller substituent on the α-C-atom, and (iii) low stereoselectivity towards α-alkenyl-α-alkyl malonic acids. To address these limitations, we expanded the structural diversity of the AMDase family by ancestral sequence reconstruction (ASR). The phylogenetic analysis of the decarboxylase revealed conserved structural motifs and key amino acids in the hydrophobic active-site cavity, a catalytic motif crucial for activity and selectivity of the enzyme. The analysis highlighted the natural distribution of amino acid exchanges previously identified via enzyme engineering. AMDase ancestors showed higher stability, activity, and, in one case, also stereoselectivity than *Bb*AMDase. While a 10 °C higher unfolding temperature of AMDase ancestors is a frequent ASR result, ancestor N131 exhibited an exceptional 294-fold improvement in half-life time and a 1.8-fold increased specific activity (25.1 U·mg⁻¹) toward 2-methyl-2-phenyl malonate in a preparative scale application. Computational analysis attributes this improved stability to a substantial increase in surface Glu side chains, which form a protective hydration shell that enhances solubility and mitigates aggregation. Ancestor N31 formed 2-methyl-but-3-enoic acid from its corresponding malonic acid in excellent optical purity (99.7% *eeR*). Finally, the stereoselectivity of the ancestors was completely inverted by switching the catalytic cysteine residues (G74C/C188G).

## Introduction

Chiral carboxylic acids are important fine chemicals with various applications, including their use in active pharmaceutical ingredients, chiral building blocks, and ligands in homogeneous catalysis.^1,2,3^ Notably, the (*S*)-enantiomers of naproxen, ibuprofen, and ketoprofen are widely employed as non-steroidal anti-inflammatory drugs (NSAIDs). Enantiopure α-alkenylcarboxylic acids find application as key intermediates in the synthesis of macrocyclic inhibitors such as Multistriatin, Zincophorin, and the Factor XIa (FXIa) inhibitor.^2,4^ The latter is currently under investigation as a new anticoagulant. In preclinical thrombosis models, it has shown outstanding efficacy while exerting minimal impact on hemostasis.^1^

Arylmalonate decarboxylase from *Bordetella bronchiseptica (Bb*AMDase) catalyzes cofactor-free C-C bond cleavage of arylmethyl and alkenylmethyl malonic acids, followed by stereoselective protonation by a catalytic Cys residue, yielding enantiomerically pure α-chiral carboxylic acids.^5^ The mechanism of AMDase requires an α-substituent with a π-electron system such as aryl, heteroaryl, naphthyl or alkenyl, which is believed to be necessary for the stabilization of the nascent negative charge upon cleavage of the carboxylic group.^5^ Regarding the size of the larger substituent, the enzyme has a wide substrate spectrum and accepts a large number of aryl, heterocyclic, and alkenyl malonates.^6–8^ To date, all characterized arylmalonate decarboxylases (AMDases) have predominantly catalyzed the formation of the (*R*)-enantiomers of α-aryl propionic acids, such as compound **2b** (**Figure 1A**).^6^ This has been exploited for the synthesis of aryl, heterocyclic, and alkenyl malonates with outstanding stereoselectivity.^8^ Inversion of the enantiopreference of *Bb*AMDase by locating the catalytic cysteine on the opposite side of the active center gave access to a wide range of (*S*)-enantiomers of α-arylpropionates.^9,10^ This made the synthesis of the biologically active enantiomers of drugs such as (*S*)-naproxen and (*S*)-flurbiprofen by enzymatic decarboxylation possible.^11^ The reduced activity caused by the switch of the position of the catalytic Cys residue could be recovered by saturation mutagenesis of residues in a hydrophobic active-site pocket, leading to variant *Bb*AMDase ICPLLG (V43I/G74C/A125P/V156L/M159L/C188G) with a 9,500-fold higher activity in the decarboxylation of **2a** compared to the initial (*S*)-selective *Bb*AMDase G74C/C188S variant.^10^

**Figure 1.**
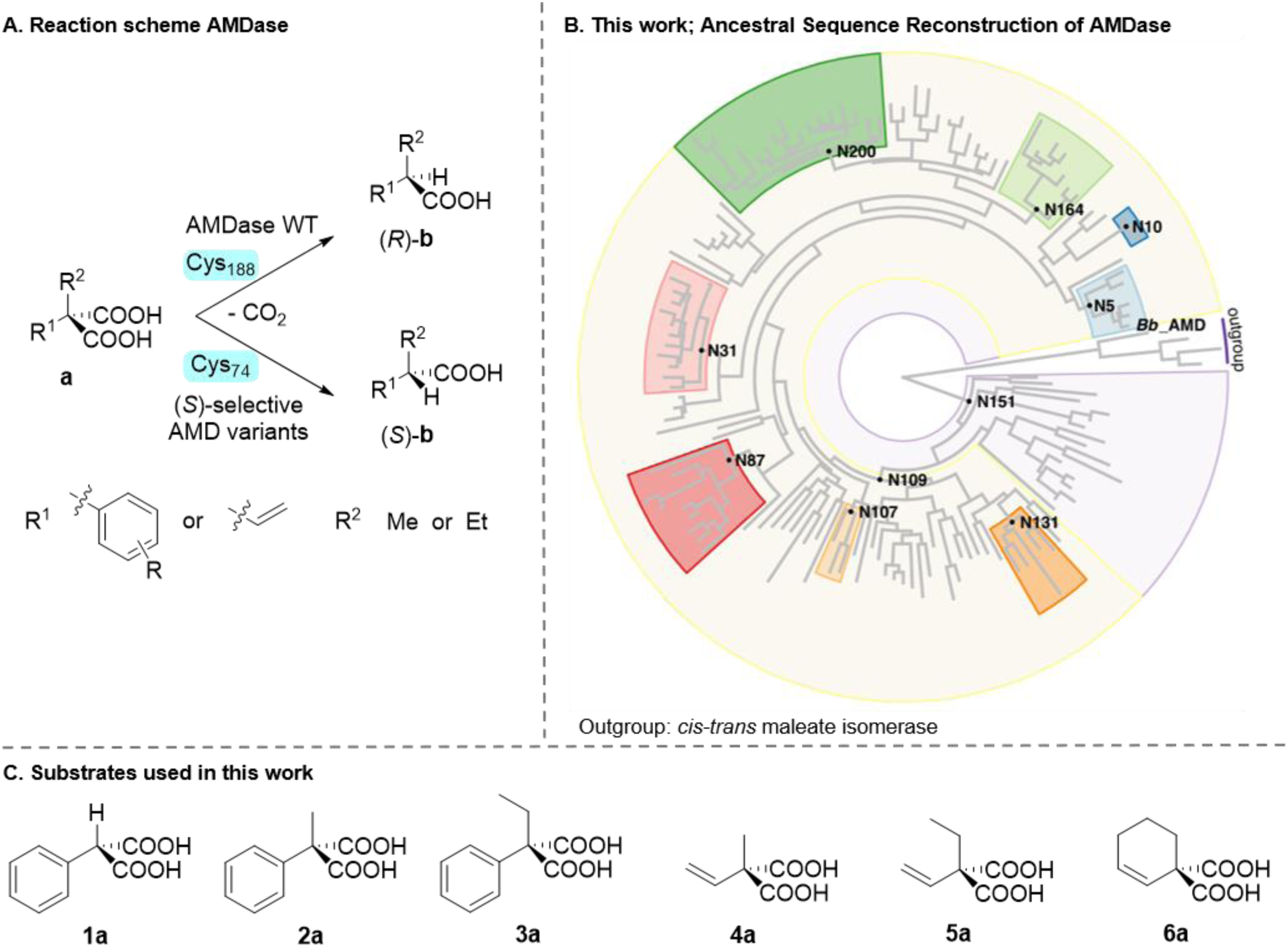
**A.** Reaction scheme of AMDase-catalyzed synthesis of α-aryl and α-alkenyl acids. Wild-type *Bb*AMDase is only able to convert α-methyl malonates, whereas AMDase ICPLLG shows additional capacity to also convert α-ethyl malonates in a highly stereoselective fashion.^12^ **B.** Maximum likelihood tree pruned to 140 and sequences visualized with phytools.^13,14^ The tree is rooted using MI as outgroup. The boxes contain the clades/descendants of the selected nodes used in this study. **C.** Malonic acids used in this work.

Despite the high value of AMDase catalysis for stereoselective synthesis, its broader application is hindered by three key challenges. The first is the limited stability of AMDase, encompassing both process stability and thermostability. *Bb*AMDase, for instance, exhibits low operational stability, with a half-life (*t*_1/2_) of only 1.2 hours under process conditions at 30 °C.^15^ Moreover, the thermostability of AMDase is also limited within the analyzed temperature range up to 50 °C. Miyamoto and Ohta reported that incubating AMDase at 40 °C for 10 minutes resulted in approximately 10% loss of activity, whereas incubation at 50 °C for the same duration led to a 50% reduction in activity.^5^ These stability limitations necessitate high catalyst loadings and constrain the operational process window, particularly regarding the reaction temperature.

In a recent industrial application of *Bb*AMDase for the synthesis of α-heterocyclic carboxylic acids, the free (i.e., non-immobilized) enzyme was employed, making stability a limiting factor.^8^ While immobilization has been shown to increase the stability of AMDase to some extent,^15^ further improvements are anticipated by combining protein engineering with immobilization strategies. Such stabilization efforts are expected to significantly increase the total turnover numbers (TTNs, mol product synthesized per mol of enzyme until its complete deactivation).^16^ From a sustainability viewpoint, improvements in enzyme stability can directly increase product yield per unit of bacterial biomass, thereby potentially reducing the energy consumption associated with biocatalyst production.^17^

The second challenge is the limited substrate scope regarding the size of the smaller substituent on the α-C-atom. While *Bb*AMDase does not accept α-ethyl substituted malonates, (*S*)-selective *Bb*AMDase ICPLLG can fully convert these substrates.^12^ The reason for this lies in the larger hydrophobic pocket that can accommodate α-ethyl substituents.

The third challenge is the lower stereoselectivity of AMDase toward smaller malonic acids with two sterically similar substituents. For example, 2-methyl-2-vinylmalonic acid (**4a**) is converted by the wild-type enzyme to the corresponding product (*R*-**4b**) with only 96% enantiomeric excess (*ee*).^12^ In practical synthesis, the purity of an enantiomerically enriched product can be further increased by methods such as selective crystallization. However, this adds extra downstream processing steps that use more resources and generate additional waste, ultimately compromising the sustainability of the process.

Ancestral sequence reconstruction (ASR) allows sequence diversification of enzyme panels used for biocatalysis. The method enables the identification of selective variants,^18^ and to explore the function or selectivity of ancestral enzymes.^19,20,21^ The gained knowledge is helpful for the guidance of protein engineering campaigns. For example, Narayan and coworkers employed ASR to decipher the sequence-selectivity relationship of flavin-dependent monooxygenases.^20^ Knowledge attained from the evolutionary path can be utilized to develop more selective biocatalysts. Moreover, ancestral proteins often show higher thermal stability than the extant enzymes, which has been exploited for protein stabilization.^22,23^

A key advantage of ASR is its capacity to accommodate variation in active-site residues. This is important because single-point mutations in the active site of an enzyme often compromise activity. Unlike traditional protein engineering by isolated point mutations, ASR provides a suitable sequence context for these amino-acid exchanges and is less likely to exert an adverse effect on enzyme function.^18^ In contrast to traditional site-directed mutagenesis, ASR allows the introduction of stabilizing or selectivity-enhancing mutations with a lower risk of compromising catalytic efficiency.

In our study, we hypothesized that ASR might provide AMDase ancestors with improved properties, specifically enhanced operational and thermal stability, higher stereoselectivity toward malonic acids bearing two similar substituents, such as compound **4a**, and possibly an expanded substrate scope. To assess this, we generated a series of ancestral AMDase variants (**Figure 1B**) and systematically characterized their stability, catalytic activity, substrate scope, and stereoselectivity against a representative panel of malonic acids (**Figure 1C**).

## Results and discussion

### Ancestral sequence reconstruction (ASR) of AMDases

To identify a suitable outgroup for the ASR, we analyzed the similarity of AMDases with cofactor-free amino acid racemases (AAR) and *cis*-*trans* maleate isomerase (MI) that also belong to the Glu/Asp racemase superfamily.^24^ Enzymes from this fold are characterized by one or two catalytic Cys-residues. Structurally, MI is most closely related to AMDase (**Figure S1**). MI catalyzes the *cis*-*trans* isomerization of the C2-C3 double bond in maleate to form fumarate by general acid/base catalysis, with two catalytic cysteine residues being responsible for the (de)protonation. AAR and MI possess two structurally conserved Cys residues at opposite sides of the substrate, whereas AMDase has only one.^25^ This Cys-residue selectively protonates the α-carbon after decarboxylation. The addition of a second Cys residue on the opposite side of the reactive center in *Bb*AMDase G74C introduced racemizing activity.^26^ Both MI and AMDase feature a conserved dioxyanion hole that stabilizes the carboxylate via a hydrogen bonding network.^25^ MI possesses an additional dioxyanion hole (as it necessitates stabilization of the second carboxylate), whereas AMDase contains a hydrophobic pocket required for decarboxylation. Unfavorable interactions between the carboxylate and the hydrophobic side chain residues of this pocket initiate cleavage of the carboxylate group. The racemase and isomerase mechanisms involve co-catalytic residues that modulate the pK_a_-values of the catalytic cysteines.^24^ Such co-catalytic residues are not found in the vicinity of the catalytic Cys-residue in AMDase. Based on this analysis, we reasoned that MI would be suitable as an outgroup to root the phylogenetic AMDase tree.

For the identification of representative decarboxylase sequences for the generation of the phylogenetic tree, we used conserved sequence patterns for AMDase suggested by Streit and coworkers. Focusing mainly on the active site, they identified six conserved sequence motifs, including catalytic residues. This approach led to the discovery of 58 unique AMDase sequences, categorized into eight distinct clusters.^27^ From each AMDase cluster, one representative sequence was used for a homology search.^27^ After a multiple-sequence alignment was done using T-COFFEE, in-house *R* scripts were used to obtain a dataset with a minimum of four amino acid differences. We aligned the remaining 222 AMDase sequences with four maleate *cis*-*trans* isomerase (MI) sequences as an outgroup using Mafft.^28^ These 226 sequences were used to construct a phylogenetic tree via *R phangorn* (**Figure S5**).^29^

As a transition, the WAG model with rate variation and invariant sites was identified as the best fitting amino acid model using the modelTest function in *phangorn*, which resembles *ProtTest*.^30,31^ We estimated the phylogenetic tree using maximum likelihood (ML) with *phangorn*. Afterward, we rooted the ML tree using the four MI outgroup sequences. As some major clades still contained a large number of sequences with high similarity, it was decided to prune the tree further, only to contain sequences with a maximum 90% amino acid sequence similarity (**Figure 1B**).

### Analysis of conserved sequence patterns and crucial amino acid residues

The obtained set of AMDase amino acid sequences was aligned, and a Hidden Markov Model (HMM) was created based on 136 AMDase sequences. Three conserved motifs in the periphery with potential structural importance were identified in addition to the six conserved sequences containing catalytically active residues described by Streit *et al*. (**Figure 2A and 2B**).^27^ Four MI sequences were chosen as the outgroup. MI possesses two structurally well-conserved catalytic cysteine residues (C76 and C194), each located on opposite sides of the substrate-binding pocket.^25^ In the following, the amino acid numbering of *Bb*AMDase will be used. The decarboxylase possesses only one Cys in the active site (C188). In position 74 (which is the position corresponding to the second Cys residue of GluR), AMDase has a Gly residue, which is highly conserved in the HMM. As substitution of G74 by Cys results in AMDase variants with inverted stereoselectivity or racemizing activity,^32^ both positions 74 and 188 were aligned for phylogenetic analysis.

**Figure 2.**
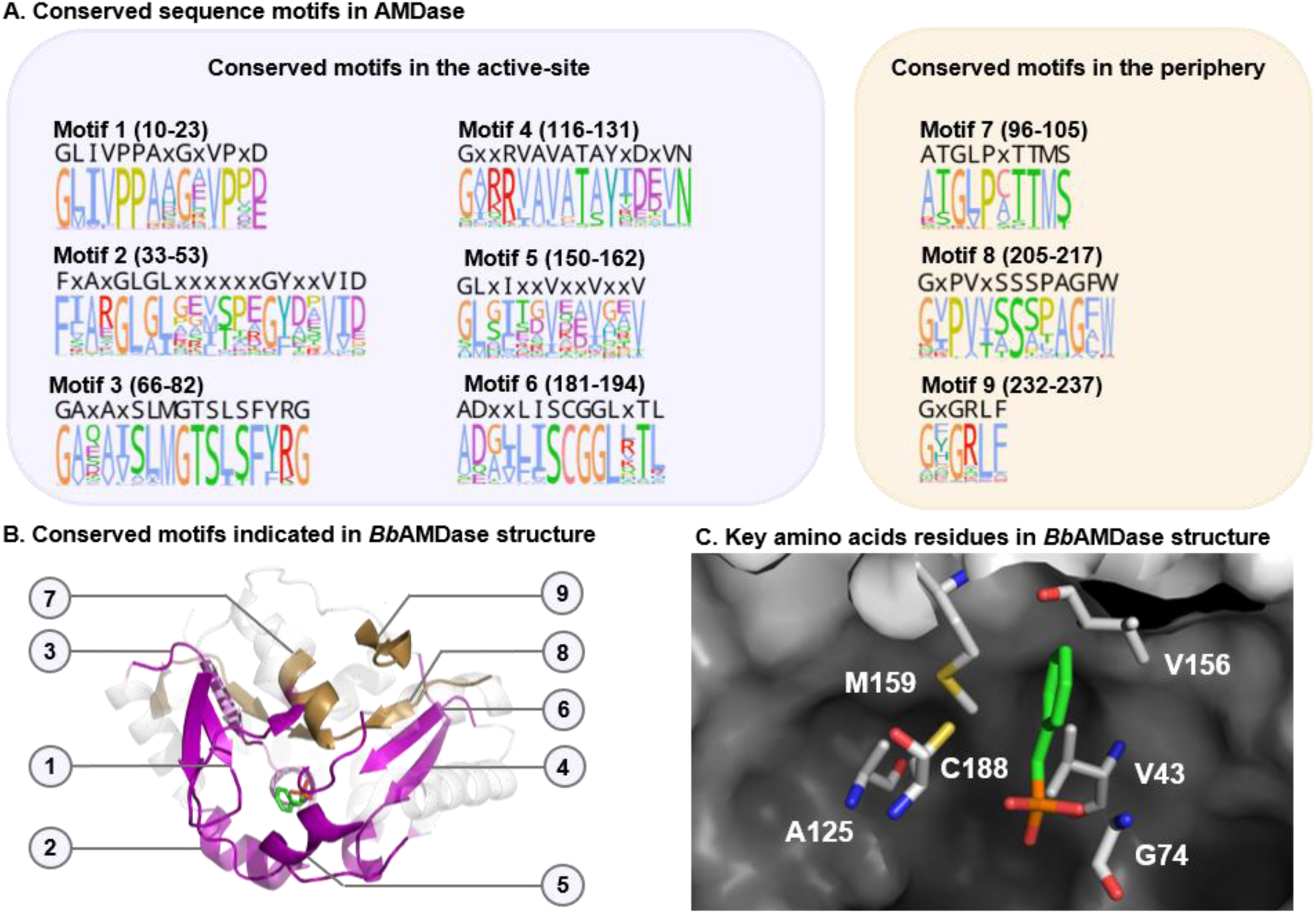
**A.** HMM of conserved sequence motifs found in AMDase. **B.** Structure of *Bordetella bronchiseptica* AMDase with conserved motifs highlighted (PDB entry: 3IP8). Motifs 1-6 are located in the reactive center of the enzyme, while motifs 7-9 are conserved motifs in the periphery. **C.** Key amino acid residues of the active site. These residues were mutated to generate (*S*)-selective AMDase ancestors.

Visual inspection of the alignment revealed some differences between MI and AMDase regarding catalytically essential residues. MI possesses an additional residue before C188. For instance, MI from *Nocardia farcinica* has a Cys-residue in this position; the other three MI have an Ala. This position was not present in any of the AMDase sequences and was not included in the HMM because it would generate a gap in all decarboxylase ancestors. The neighboring amino acids of the catalytic Cys of AMDase are highly conserved.

Saturation mutagenesis of active-site residues of *Bb*AMDase revealed several positions in the hydrophobic pocket with a strong impact on activity.^6,9,33^ Therefore, amino acids leading to higher activity were contrasted to the natural abundance of amino acids in these positions. V43 is a key residue for activity as it is positioned in close proximity to the α-carbon of the malonic acid substrate. Substitution by isoleucine increased the activity of the (*S*)-selective variants.^10^ In position 43, Val is the most abundant residue throughout the AMDase and MI sequences. Y48 participates in the dioxyanion hole. The substitution Y48F was identified as a key mutation in a saturation mutagenesis campaign using the NNK codon.^9^ Interestingly, Phe is indeed the second-most-conserved residue at this position, after Tyr. A125 is situated in a turn between two alpha-helices. MI has a Pro in this position. The rationally introduced mutation A125P increased the decarboxylating activity of (*S*)-selective AMDase variants.^10^ Yet none of the analyzed AMDase sequences contained a Pro at this position. M159 is part of the hydrophobic pocket. Its exchange by valine, leucine, or isoleucine strongly increased the activity of both (*R*)-selective and (*S*)-selective AMDase variants.^6,9^ Interestingly, Val and Ile are the most conserved residues in this position and, therefore, present in most ancestors. MI has a Val in this position (**Table 1**).

**Table 1.** Conservation of active-site residues identified in protein engineering studies as activity determinants of the decarboxylase.

| **Position**<br>**(in *Bb***<br>**AMD)** | ***Bb* AMD** | **HMM**<br>**AMDase –**<br>**most**<br>**abundant** | **HMM AMD**<br>**– Second**<br>**most**<br>**abundant** | **Outgroup** | **Root**<br>**AMD**<br>**and MI**<br>**(N1)** | **Other**<br>**comments** |
| --- | --- | --- | --- | --- | --- | --- |
| 15 | Proline | Proline | Proline | Serine | P/S | - |
| 43 | Valine | Valine | Methionine | Valine | M/V | - |
| 48 | Tyrosine | Tyrosine | Phenylalanine | Leucine | Y/L | - |
| 73 | Methionine | Methionine | Only 1 F and<br>1 T (out of<br>136 AMDase) | Alanine | M/A | Motif AMD:<br>LMGTSLSF<br>Motif MI:<br>YACLVAxM<br>(position: 72-79) |
| 74 | Glycine | Glycine | Glycine | Cysteine | G/C | Probability G<br>70.9% and C<br>23.1%. 6%<br>posterior<br>probability of<br>observing any<br>other state (aa) |
| 77 | Leucine | Leucine | Isoleucine | Alanine | L/A | - |
| 125 | Alanine | Alanine | Serine | Proline | A/P | - |
| 156 | Valine | Valine | Isoleucine | Asparagine | Valine | - |
| 159 | Methionine | Valine | Isoleucine | Valine | V/I | - |
| 188 | Cysteine | Cysteine | Cysteine | Cysteine | Cysteine | - |
| Between 98 and 99 |  |  |  | Insertion xxGxx |  | Conserved motif not in the active site |

### AMDase ancestor selection and reconstruction

To cover the influence of the complete tree, we selected ten nodes for further investigation. Eight ancestors were chosen to cover different clades of the tree. Two ancestors show a higher hierarchy. Node 109 covers all these clades, and node N151 represents the common ancestor of AMD, which is also closer and likely influenced by the outgroup. Nodes N5, N10, N31, N87, N107, N109, N131, N151, N164, and N200 were selected for biochemical characterization (indicated as a black dot in **Figure 1B**). Based on the phylogeny and model, we performed ASR. We computed the marginal reconstruction for the ten selected nodes using marginal reconstruction in *phangorn*.^34^ The state with the highest marginal likelihood was used as the ancestral state. Whereas in phylogenetic analysis, gaps are usually treated as ambiguous states, we define gaps as their own state. Gaps were treated as insertions or deletions. This differs from the approach of Grasp, which defines two transition processes, one for the amino acid replacement and one for the gaps.^35^

### Biochemical characterization of AMDase ancestors

#### Gene expression, solubility analyses, and initial activity test

The genes of ten selected AMDase ancestors were heterologously expressed in *E. coli* BL21 (DE3) to compare their biochemical properties with *Bb*AMDase. All ten ancestors exhibited expression levels similar to *Bb*AMDase (30–45% soluble expression yield in batch-to-batch deviations, **Figure S10**). Qualitative analysis of produced proteins by SDS-PAGE displays prominent bands within the soluble fraction of the cell lysate. Notably, N151 displayed a low amount of inclusion bodies, and N200 an equal distribution of soluble and insoluble protein (**Figure S8**). Attempts to enhance soluble protein production by reducing induction temperature or IPTG concentration did not lead to further improvement.^36^

Eight of ten ancestors showed activity in a pH indicator assay with **1a** as substrate (**Figure S7**).^6,37^ From these eight, N87 and N151 showed significantly lower activity towards **1a** compared to the wild-type *Bb*AMDase, whereas the remaining ancestors displayed similar activity as *Bb*AMDase.

#### Analysis of the thermostability of AMDase ancestors

We examined the thermal stability of the AMDase ancestors. The unfolding temperatures (*T*_m_) were assessed through Differential Scanning Fluorimetry (DSF, Thermofluor).^38^ *Bb*AMDase displays a *T*_m_ value of 55 °C (**Figure 3A**). Significant variation in thermal stability was observed among the ancestral enzymes. Four ancestors exhibited enhanced thermal stability, with N164 having the highest *T*_m_ value (65 °C), remarkably 10 °C higher *T*_m_ than *Bb*AMDase. Both N5 (*T*_m_ of 62 °C) and N131 (*T*_m_ of 62 °C) showed notably higher unfolding temperatures, while ancestor N200, based on extant enzymes from *Ramlibacter* species, showed similar thermal stability to *Bb*AMDase. The remaining selected ancestors proved less thermally stable than *Bb*AMDase, with N10 and N151 displaying *T*_m_ values of 47 °C and 26 °C, respectively (**Figure 3A**).

**Figure 3.**
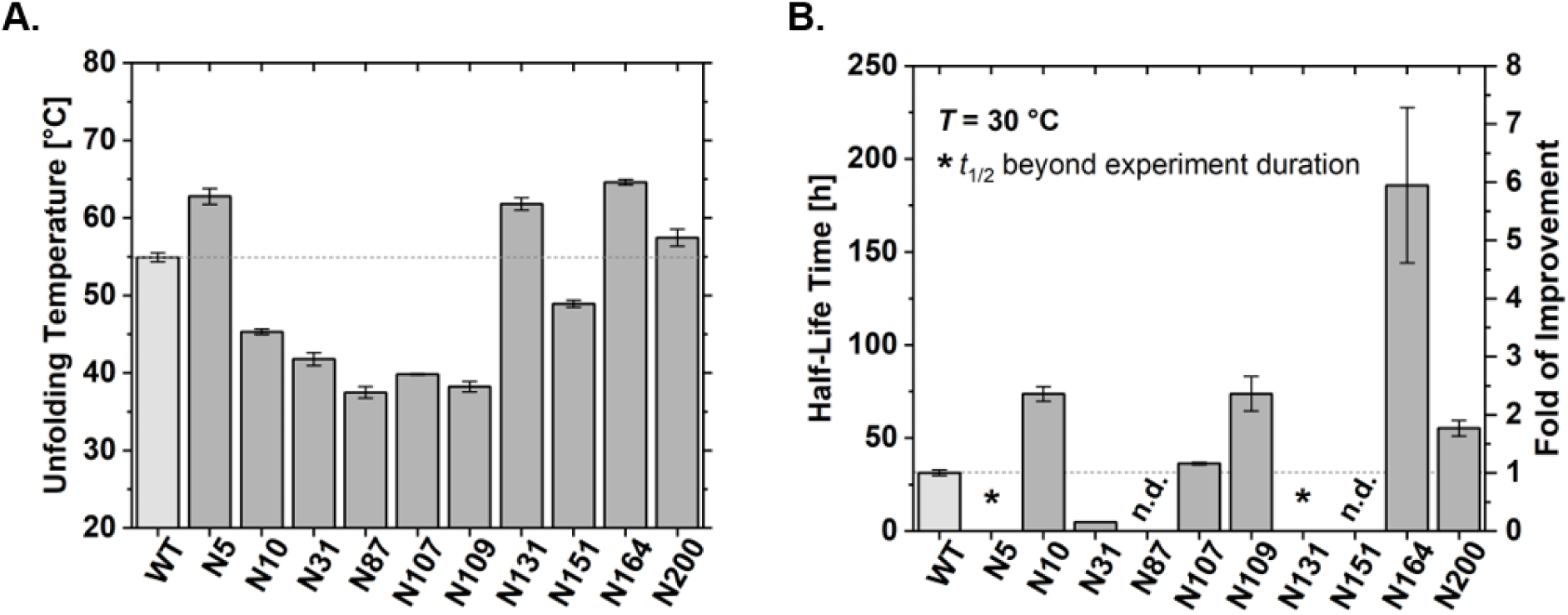
**A.** Unfolding temperature of AMDase ancestors compared to the *Bb*AMDase determined by Differential Scanning Fluorimetry (DSF, Thermofluor) measured in technical triplicate with purified enzymes in concentrations of 5, 10, and 15 μM in KPi (48 μL, 50 mM, pH 8) with 2 μL SYPRO Orange (200x diluted with ddH_2_O). **B.** Half-life time of purified AMDase ancestors at 30 °C in Tris-HCl buffer (50 mM, pH 8) determined with the initial activity of the respective sample with the conversion of 20 mM **1a**. The half-life times of AMD N87 and N151 were not determined due to activity rates below the detection limit. *Decrease in activity over 485 hours: AMDase N5: 32%, N131: 23%. No exponential fitting could be done (**Figure S33, S34**).

To investigate the contribution of increased thermostability to catalytic activity over time, we measured the half-life (*t*_1/2_) of each enzyme by incubating the purified proteins at the reaction temperature (30 °C). Interestingly, ancestors N10, N107, and N109 deviated from the expected trend, exhibiting longer half-lives than *Bb*AMDase despite having lower unfolding temperatures (**Figure 3B**). This suggests that the inactivation of the extant enzyme is also associated with factors other than unfolding.

The ancestors N5, N131, N164, and N200 showed a strikingly prolonged enzymatic activity, the extent of which is surprising given their moderately higher unfolding temperatures. Remarkably, the stability of N5 and N131 extended beyond the experiment’s duration. Even after 485 hours at incubation temperature, N5 activity retained 68%, and N131 still retained 76% (**Figure 3B**).

We decided to increase the incubation temperature to 45 °C to further assess the thermal stability of the reconstructed ancestral enzymes (**Figure 4**). *Bb*AMDase served as the benchmark, with a half-life of 1.3 hours under these conditions. Ancestor N5 showed a 72-fold improvement, with a half-life of 92.6 hours. The most stable variant, N131, exhibited a 294-fold increase in stability (*t*_1/2_ = 381 hours) and retained 37% of its activity even after 458 hours of incubation. This exceptional stability translated into vastly superior catalytic endurance, as evidenced by the observed total turnover numbers (TTN) (**Table S7**).^16^ Compared to *Bb*AMDase’s TTN of 474,780, N5 and N131 achieved values of 1.3 × 10^8^ and 3.0 × 10^8^, representing a ∼275- and 633-fold improvement. These results highlight the potential of ASR as a powerful strategy for engineering enzymes with enhanced thermal stability and higher TTN. However, not all reconstructed ancestors display improved properties, and the observed half-life times of the investigated ancestors had not been anticipated on the basis of the unfolding temperatures; therefore, a reliable prediction of the best ancestral variants is challenging. Therefore, selecting ancestral nodes without experimental data from extant enzymes continues to pose a significant limitation.

**Figure 4.**
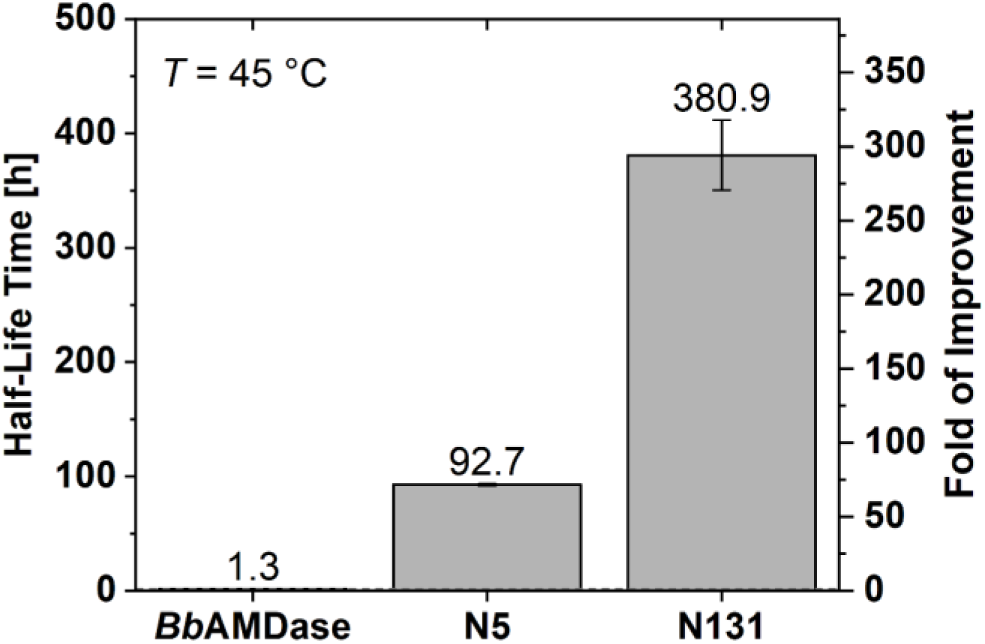
Half-life time of purified AMDase ancestors N5 and N131 compared to *Bb*AMDase at 45 °C in Tris-HCl buffer (50 mM, pH 8), determined with the initial activity of the respective sample, with the conversion of 20 mM **1a** (**Figure S35**).

#### Molecular Dynamics Simulations and Analysis

Following from successes in recent work,^39,40^ to better understand the origins of the increased thermal stability and half-life of the ancestral enzymes (N131 in particular), we performed sequence composition analysis using the ProtParam tool on the ExPasy webserver,^41^ predicted protein thermostability from sequence based on propensity scores calculated using the scoring-card method based on the SCMTPP webserver,^42^ predicted scaled solubility from sequence using Protein-Sol,^43^ as well as 10 x 0.5 μs MD simulations of wild-type *Bb*AMDases and N131, to monitor differences in sequence and structural properties across systems.

Based on this analysis (**Table S11**), we observe that, as in our prior work on borneol dehydrogenase (BDH),^40^ SCMTPP propensity scores predict enhanced thermostability for N131 compared to wild-type *Bb*AMDase (increase from 404.7 to 421.8). We also observe that, compared to wild-type, N131 shows a substantive (6%) increase in Glu residues, manifesting as 14 additional negatively charged residues compared to wild-type *Bb*AMDase. Examination of the predicted structure of N131 (**Figure 5 and S49**) indicates that these Glu side chains are primarily located on the surface of the protein, and expected to impact its solubility. Indeed, we observe a small drop in instability index from 44.4 in wild-type *Bb*AMDase to 42.2 in N131 (**Table S11**), bringing N131 slightly closer to the stability cutoff of <40.^44^ N131 also shows a reduced aliphatic index compared to wild-type (proteins with high thermal stability typically have higher aliphatic indices),^45^ but also a lower GRAVY (Grand Average of Hydropathicity) score,^46^ suggesting the protein has become more hydrophilic. This is further supported by scaled solubility predictions using Protein-Sol,^43^ which suggest that N131 is more soluble than wild-type *Bb*AMDase (**Table S11**).

**Figure 5.**
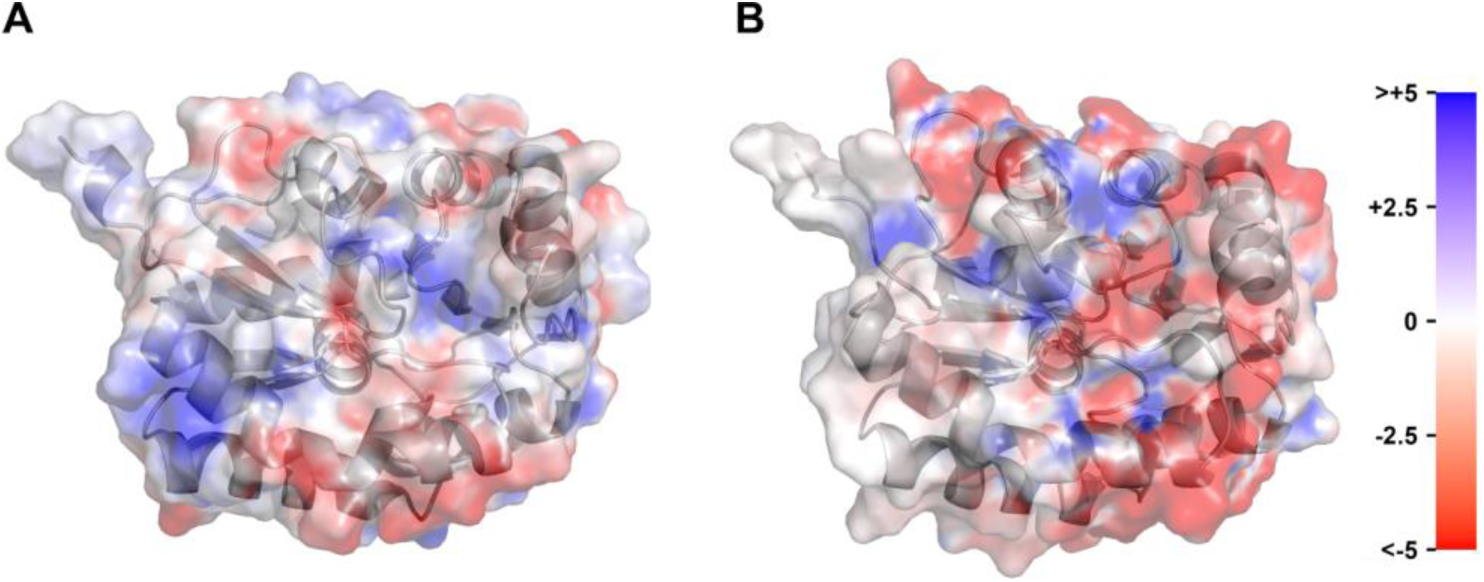
Comparison of the surface charge distribution of (**A**) wild-type AMDase and the (**B**) N131 ancestor, based on 10 x 0.5 μs simulations of each system performed at 300 K. Analysis was performed as described in the **Materials and Methods**. This figure illustrates the greater negative surface charge of N131 compared to the wild-type, caused by the increased propensity of surface Glu residues (see **Table S12**).

For structural analysis of thermostability, we predicted the structures of N5, N31, and N131 by Chai-1.^47^ AMDase has the Glu/Asp-Racemase fold that is characterized by two structurally similar α/β domains containing a central β-sheet surrounded by α-helices. The active site is located in the cleft between the two domains. This two-domain arrangement positions the catalytic Cys188 and the residues forming the oxyanion hole and hydrophobic substrate-binding pocket around the bound malonate substrate. Visual inspection of the structures (**Figure S50**) showed that all structural features of *Bb*AMDase are found in the ancestral proteins, and no major differences were identified in the structural comparison.

To validate our prediction of differential solubility, we performed 10 x 0.5 μs MD simulations of the wild-type enzyme and the N131 ancestor, to capture any differences in the solvation of these two systems. **Figure S49** shows a visual comparison between the water shells of the two enzymes, as well as the position of the surface Glu residues in N131 (that are not present in wild-type *Bb*AMDase). Simulations were performed at both 300K and 360K (which over the melting temperature of both enzymes), to see how stable the surface hydration of each system is at elevated temperatures. Comparison of the total solvent accessible surface area (SASA, **Figure S51**) for both variants shows that N131 has higher baseline SASA than the wild-type at both temperatures, with both systems showing increased SASA at higher temperature, consistent with thermal expansion/unfolding.

Examining the breakdown of the total SASA into its hydrophobic and hydrophilic components (**Figure S51**) similarly shows that increasing temperature increases both hydrophobic and hydrophilic SASA (and thus total SASA) in both variants, as both proteins expose more surface area to the solvent. Further, while N131 shows a modest increase in hydrophobic SASA compared to the wild-type, there is a substantive increase in hydrophilic SASA, resulting in a drop in the hydrophobic SASA ratio. This reflects a stable hydrophobic core while the surface remains shielded by water (likely driven by the additional Glu side chains), protecting N131 from thermal aggregation.

Further, heating both wild-type and N131 to 360 K results in a drop in intramolecular hydrogen bonds and hydrophobic interactions (**Figure S52**), in agreement with the thermal expansion hinted at by the changes in SASA. However, at both temperatures, N131 maintains both more intramolecular hydrogen bonds and hydrophobic interactions than the wild-type, indicating more dense internal packing, leading to stability. Further, N131 shows a greater increase in salt bridge interactions with temperature compared to wild-type, where instead salt-bridge interactions are lost with temperature. This suggests the formation of heat-induced salt bridges in N131 that contribute to maintaining the stability of the protein. This can be observed visually in the surface charge maps shown in **Figure 5**.

Finally, **Figure S53** shows the radial distribution function of the water oxygens around the carboxylate oxygens of wild-type and N131 Glu residues (**Figure S53**). Based on this data, N131 shows a higher intensity first hydration peak compared to wild-type at both temperatures, indicating the presence of an ordered shell of water molecules interacting with the surface Glu, that protect N131 from thermal unfolding. Further, at high temperature, the first solvation peak for both variants drops (although it remains higher for N131 than wild-type), freeing the glutamic acids on the surface to become involved in salt bridges (see also surface charge distribution in **Figure 5**).

Taken together, this suggests that the presence of the additional surface Glus in N131 has created a protective tight water shell around the protein, improving solubility and stability, and reducing propensity towards aggregation via dynamic surface charge networks, accounting for the impressive increase in half-life of this variant.

### Enzymatic Activity of Native and Engineered Ancestors

In activity tests, AMDase ancestor N151 exhibited very low catalytic activity, which may be attributed to the influence of the outgroup during ASR. Generally, higher-order ancestors such as N151 and N109 tend to have greater sequence prediction uncertainty, which possibly contributed to their significantly reduced enzymatic function. Ancestor N87 represents a clade comprising nine extant enzymes, predominantly from *Terriglobia* species, that are genetically diverse. This broad diversity may have led to a soluble but catalytically impaired ancestral enzyme. Overall, these results indicate that accurately predicting the catalytic function of early ancestral enzymes within a specific clade remains challenging, particularly in the absence of functional data from their extant counterparts.

In contrast, ancestor N5 is phylogenetically closely related to *Bb*AMDase and displayed slightly reduced activity toward substrate **1a** (**Figure 6A**). Interestingly, the more distantly related ancestor N10, based on sequences from *Alcaligenes*, exhibited higher activity. Ancestors N31, N107, N131, N164, and N200 showed catalytic activities comparable to *Bb*AMDase.

**Figure 6.**
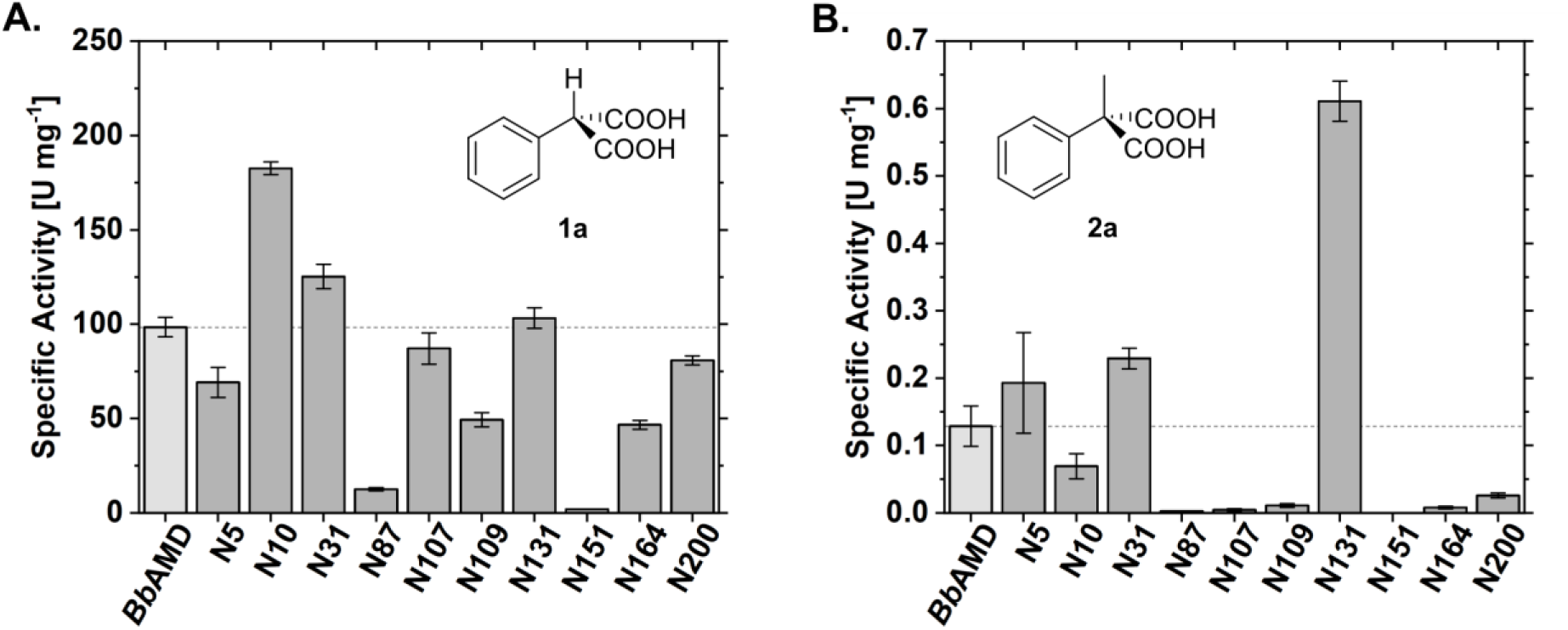
**A.** Specific activity (U·mg⁻¹ total protein) of cell-free extracts of the ancestors in comparison to *Bb*AMDase determined by HPLC. Reaction conditions: 20 mM **1a**, Tris-HCl (pH 8, 50 mM), at 30 °C and 600 rpm; max. conversion of 10%, 4.25 min total assay time. **B.** Specific activity (U·mg⁻¹) of the *Bb*AMDase compared to the ancestors. Reaction conditions: 20 mM **2a**, Tris-HCl (pH 8, 50 mM), at 30 °C and 600 rpm; max. conversion of 10%, 4.25 min total assay time. This study was performed on CFE.

Subsequently, we evaluated the activity of the prochiral **2a**. Similar to *Bb*AMDase,^6^ all ancestral variants showed lower activity toward **2a** than toward **1a** (**Figure 6B**). This reduced activity is likely due to a combination of steric and electronic effects introduced by the additional methyl group. Ancestors N107, N109, N164, and N200 show specific activity towards **2a** below 0.03 U·mg⁻¹, notably lower than *Bb*AMDase (0.13 U·mg⁻¹) (**Figure 6B**). Besides low activity towards **1a**, AMDase N87 and N151 also displayed minimal conversion of **2a**, with initial rates below the detection limit. Hence, it was decided to exclude them from further activity measurements. Compared with *Bb*AMDase, N5, N10, and N131 showed similar or enhanced activity. Moreover, we were delighted to observe a five-fold increase in the activity of AMD N131 towards **2a**. Interestingly, N131 converts **2a** faster than *Bb*AMDase, with a *k*_cat_ of 34 s^−1^ and a K_M_-value of 0.3 mM for the former compared to a k_cat_ of 29 s^−1^ and a K_M_-value of 25.5 mM for the extant enzyme (**Table 2**),^5^ making this early yet distant AMDase ancestor a promising candidate for further examination.

**Table 2.** Kinetic parameters of AMDase ancestors N31 and N131 with 2-methyl-2-phenyl malonate (2a) and 2-methyl-2-vinyl malonate (4a).

| AMDase ancestor | Substrate | $K_{\text{M}}$ (mM) | $k_{\text{cat}}$ (s<sup>-1</sup>) |
| --- | --- | --- | --- |
| N31 | **2a** | 6.0 | 10.0 |
|  | **4a** | 9.9 | 0.1 |
| N131 | **2a** | 0.3 | 33.9 |
|  | **4a** | 1.0 | 0.8 |

In conclusion, all ancestral enzymes were predominantly localized in the soluble fraction; however, their catalytic activities varied considerably. The common AMDase ancestor, N151, exhibited little to no catalytic function. In contrast, the remaining nine ancestral variants demonstrated the ability to convert **1a** at levels comparable to the reference enzyme *Bb*AMDase. Notably, only four of these ancestors displayed appreciable activity toward the prochiral substrate **2a**.

We then explored whether one of the (*R*)*-*selective ancestors would be able to convert the sterically demanding α-ethyl substrates **3a** and **5a**. *Bb*AMDase has been reported not to accept α-ethyl substrates.^5,48^ Using cell-free extract (4–5 mg·mL^−1^ total protein), we observed partial conversion of **3a** for *Bb*AMDase after a reaction time of 16 h (30 °C). Apart from the common AMDase ancestor N151, all ancestors showed acceptance of **3a** to a certain extent (**Table S1**). Notably, full conversion was only achieved by N131.

Based on the catalytic Cys residue at position 188, we inferred that all ancestors should have (*R*)-selectivity, which was confirmed by chiral GC analysis (see **Figure 8**). Relocating C188 to the opposite side of the active site should generate (*S*)-selective decarboxylases.^32^ We selected the ancestors N5, N131, and N164 for further point mutations in the active site to achieve (*S*)-selectivity, as previously done with the *Bb*AMDase ICPLLG mutant (positions highlighted in **Figure 2C**),^10^ while also retaining the increased stability of the designed ancestors. Miyauchi et al. showed that additional mutations of the hydrophobic pocket of *Bb*AMDase were necessary in order to compensate for the detrimental effect of these two substitutions on the overall activity of the enzyme.^9^ Therefore, we incorporated several amino acid exchanges from *Bb*AMDase ICPLLG into ancestors N5, N131, and N164, yielding eight ancestral proteins (**Table 3**).

**Table 3.**
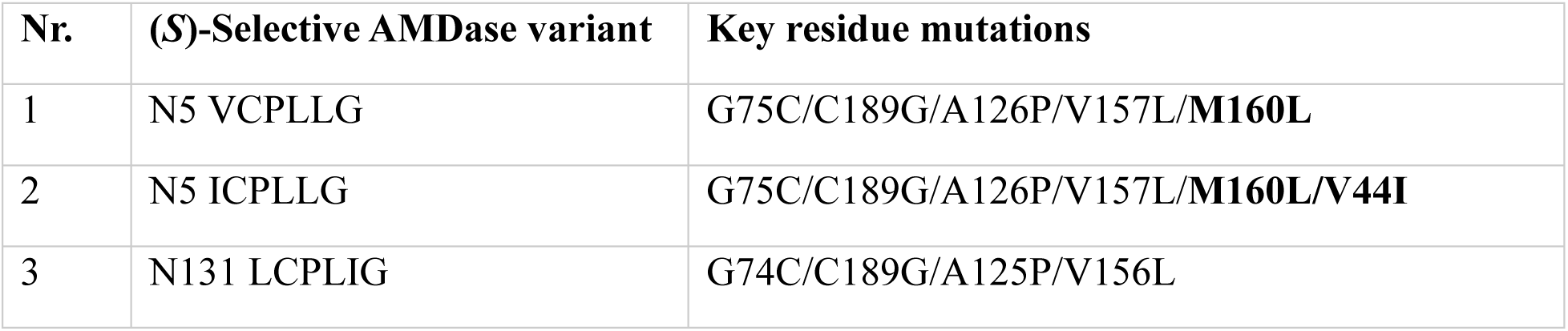
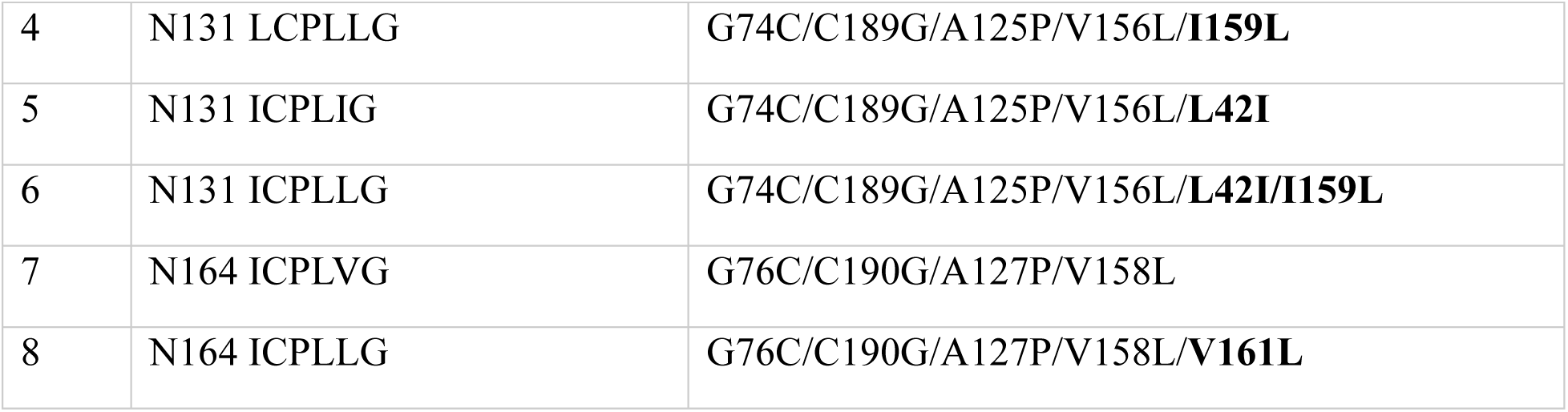
Amino acid substitutions of (*S*)-selective AMDase variants created from ancestral sequences N5, N131, and N164. All (*S*)-selective ancestors have the following mutations: G74C/C188G/A125P/V156L, numbering according to the *Bb*AMDase sequence. Amino acid substitutions L42I and M/I/V159L are shown in bold.

To assess the enzymatic activity of the (*S*)-selective ancestors towards **1a**, one variant from each ancestor was chosen (N5 VCPLLG, N131 LCPLLG, and N164 ICPLVG). We were pleased to find that the N5 variant showed a 2-fold increase in activity compared to *Bb*ICPLLG, 14.8 U·mg⁻¹ and 7.7 U·mg⁻¹, respectively (**Figure 7**). Consequently, we also measured the specific activity of the other N5 variant (N5 ICPLLG), whose activity (21 U·mg⁻¹) showed a 2.7-fold increase compared to *Bb*ICPLLG. The difference between the N5 variants is the V44I mutation (position 43 in *Bb*AMDase), which was reported to enhance the specific activity.^10^ On the other hand, the corresponding (*S*)*-*selective N131 LCPLLG and N164 ICPLVG were barely active (**Figure 7**). This is in contrast with their originating (*R*)-selective ancestral variants being the two most active of the characterized set (**Figure 6**).

**Figure 7.**
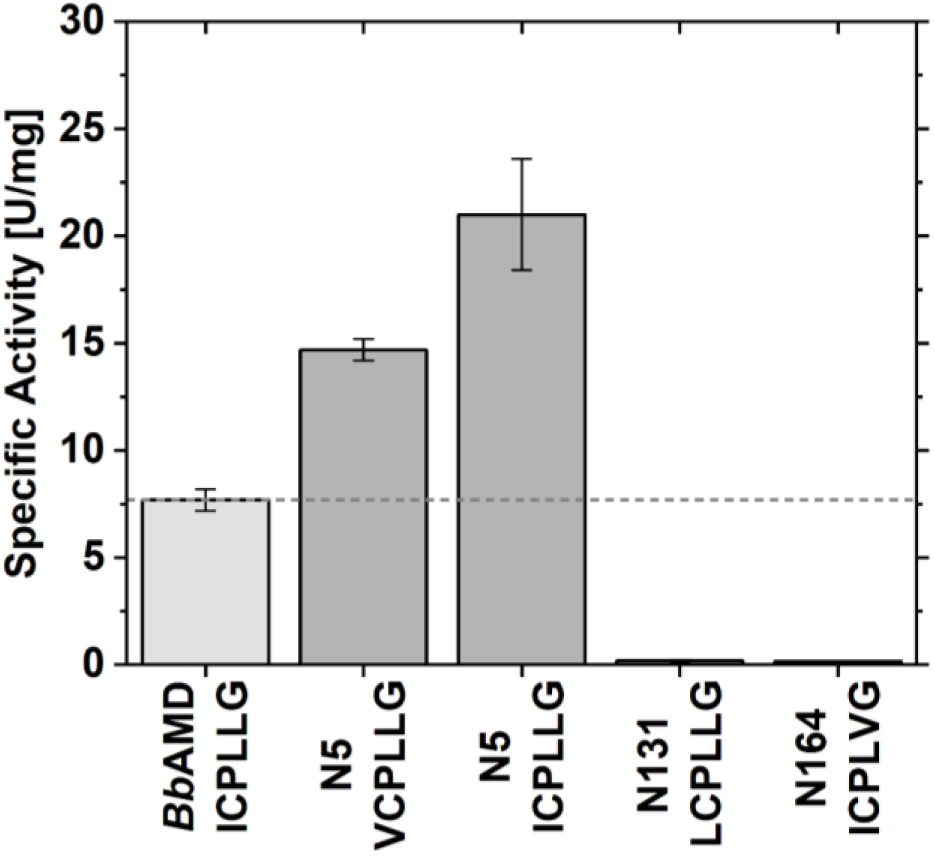
Specific activity (U·mg⁻¹) of the *Bb*AMDase ICPLLG, compared to the (*S*)-selective ancestral variants N5 VCPLLG, N5 ICPLLG, N131 LCPLLG, and N164 ICPLVG. Reaction conditions: 20 mM **1a**, Tris-HCl (pH 8, 50 mM), at 30 °C and 600 rpm; max. conversion of 10%, 4.25 min total assay time. This study was performed on CFE.

Overall, we can report (*S*)-selective ancestral AMDase variants with improved catalytic activity compared to *Bb*AMDase ICPLLG. Although AMDase N131 exhibited activity similar to that of *Bb*AMDase, the change in stereoselectivity appeared to significantly reduce its activity relative to its (*S*)-selective *Bb*AMDase variant. Conversely, AMDase N5 showed lower activity compared to *Bb*AMDase. Nevertheless, engineered (*S*)-selective N5 AMDases demonstrate significantly higher activity than *Bb*AMDase ICPLLG. This emphasizes the difference in enzymatic activity between the (*R*)-selective ancestors and their engineered (*S*)-selective variants, while also illustrating the complexity involved in predicting activity in engineered ancestors. Additionally, the structural differences between *Bb*AMDase and the ancestral enzymes make it challenging to explain the observed results.

### Analysis of the stereoselectivity of AMDase ancestors and their variants

With all ancestors at hand, we evaluated the acceptance and stereoselectivity toward prochiral substrates (**2a**–**5a**), (**Figure 8**). We used α-methyl aromatic malonate **2a** as a model for the synthesis of chiral α-arylpropionic acids. As hypothesized, the (*R*)-selective ancestors gave a stereochemical outcome favoring (*R*)*-***2b.** Vinylic malonate **4a** was investigated because *Bb*AMDase and its (*S*)*-*selective mutants show unsatisfactory stereoselectivity towards this substrate.^12^ Finally, we investigated whether the ancestors could convert α-ethyl malonates (substrates **3a** and **5a**), considering that *Bb*AMDase does not accept them as substrates.^12^ The 2-aryl-carboxylic acids were derivatized to yield their corresponding methyl esters, which were subsequently analyzed by chiral GC. Hence, to determine the stereoselectivity of aromatic compounds, full conversion is not required, as the corresponding dimethyl malonates are stable and cannot undergo spontaneous decarboxylation. In the case of the 2-vinyl-malonic acids, however, the instability of malonic acids at elevated temperatures makes an analytical yield of >99% necessary in order to avoid spontaneous decarboxylation of residual substrate during chiral GC measurements. Therefore, an incomplete conversion results in no measurable stereoselective preference.

**Figure 8.**
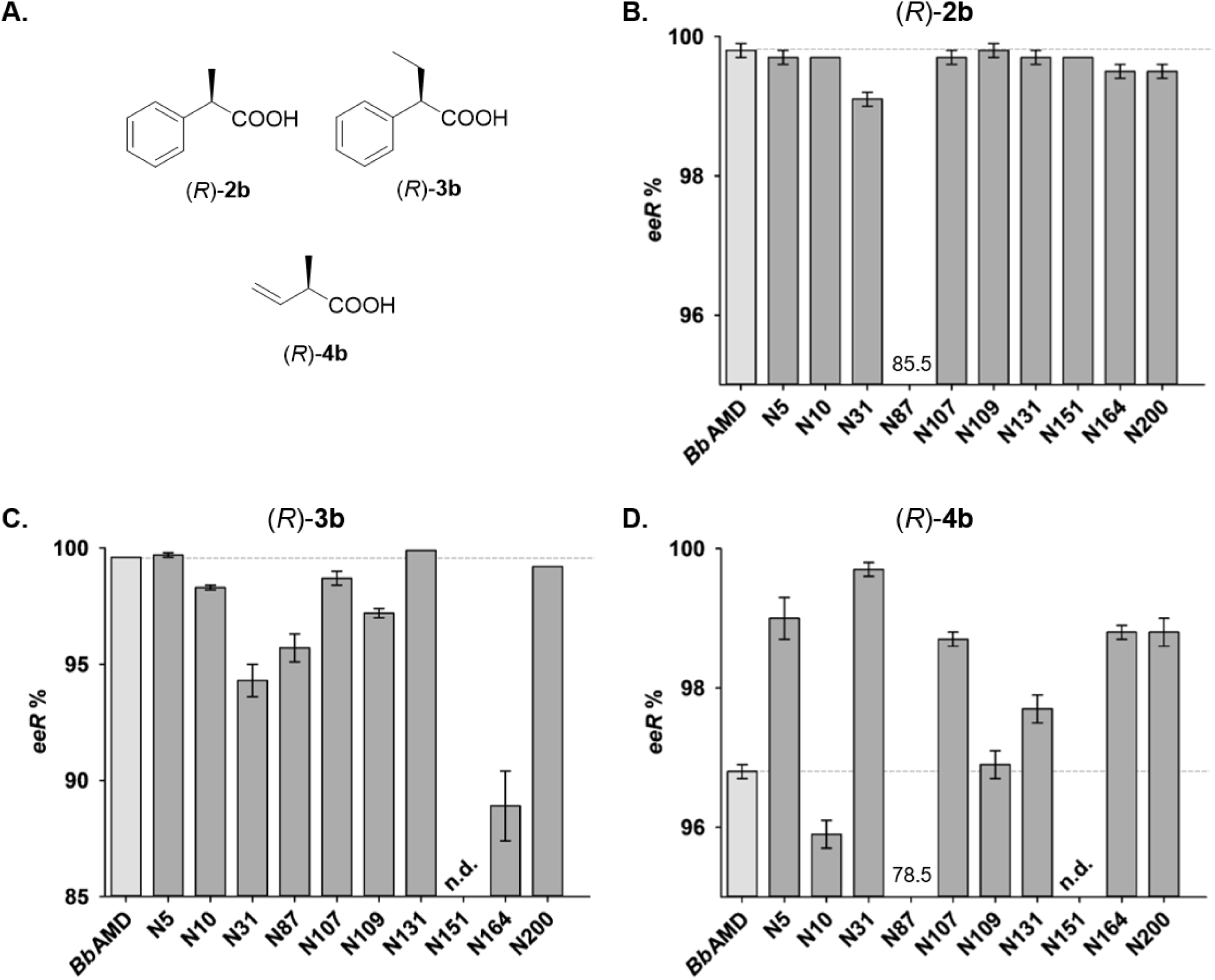
Chiral products obtained by engineered (*R*)-selective AMDase variants. **B. – D.** Stereoselectivity of engineered (*R*)-selective AMDase variants after the decarboxylation of substrates **2a**-**4a**. Reactions were performed at 30 °C using a 5 mM substrate concentration for **2a** and **3a** and 2.5 mM for **4a**. Chiral aryl carboxylic acids (**2b** and **3b**) were derivatized to their corresponding methyl esters prior to chiral GC analysis. All reactions were performed in triplicate, and the stereoselectivity was determined by chiral GC.

*Bb*AMDase and nine ancestors achieved excellent stereoselectivity towards (*R*)*-***2b**, ranging from 99.1–99.8% *ee*. Only ancestor N87, which proved to be unstable and exhibited minimal reactivity, showed only 85.5% *eeR* based on the detectable amount of product. *Bb*AMDase exhibits moderate (*R*)-selectivity for 2-vinyl-2-methyl malonate (**4a**) with an enantiomeric excess (*ee*) of 96.8%.^12^ Compared to the extant decarboxylase, seven of the ten selected ancestors showed a higher selectivity towards **4a**. Remarkably, N31 was found to produce (*R*)*-***4b** in an exquisite enantiopurity of 99.7% *ee* (**Figure S3**). AMDase N31 was identified as an ancestor of a clade of extant AMDases from *Variovorax* species. Also, the excellent selectivity of N5 leading to 99.0% *ee* of (*R*)-**4b** is notable, as N5 is the closest ancestor of the relatively unselective *Bb*AMDase from the selected nodes of the constructed phylogenetic tree. Optical purity of formed product **3b** was >98% *eeR* in all cases except for N31 (94.3% *eeR*) and N164 (88.9 % *eeR*). Furthermore, we assessed how our ancestors accepted **5a**, which is accepted well by the (*S*)-selective AMDase variant *Bb*AMDase ICPLLG, but not *Bb*AMDase.^12^ Ancestors produced only traces of **5b** as indicated by TLC analysis (<5%).

Next, we examined the stereoselectivity of the engineered (*S*)-selective ancestors towards malonates **2a–5a** (**Figure 9**). When converting **2a**, the stereoselectivity of the (*S*)-selective variant ICPLLG is only moderate (89.0% *ee*). Interestingly, two of the ancestral variants, namely, N5 VCPLLG and N131 ICPLLG, were able to form (*S*)-**2b** with excellent stereoselectivity of ≥99.8% *ee*. Potentially due to a more sterically hindered hydrophobic pocket, two of the ancestral variants, N5 ICPLLG and N131 LCPLLG, display poor stereoselectivity when converting **3a**. When converting vinyl malonate **4a**, AMDase ICPLLG displays poor selectivity of only 68.8% *ee* towards (*S*)-**4b**. None of the (*S*)-selective ancestral variants can convert **4a** with higher selectivity. All (*S*)-selective ancestor variants were able to convert **5a** with complete conversion, which allowed us to determine the enantiomeric excess. AMDase ICPLLG, as well as most of the ancestral variants, displayed excellent selectivity in the production of (*S*)-**5b** (>99.2% *ee*). When converting **5a**, the only enzyme variant that could achieve higher stereoselectivity than ICPLLG is N5 VCPLLG, with >99.9% *ee* towards (*S*)-**5b**. Therefore, the stereoselectivity was indeed shown to be switchable for ancestral AMDase sequences.

**Figure 9.**
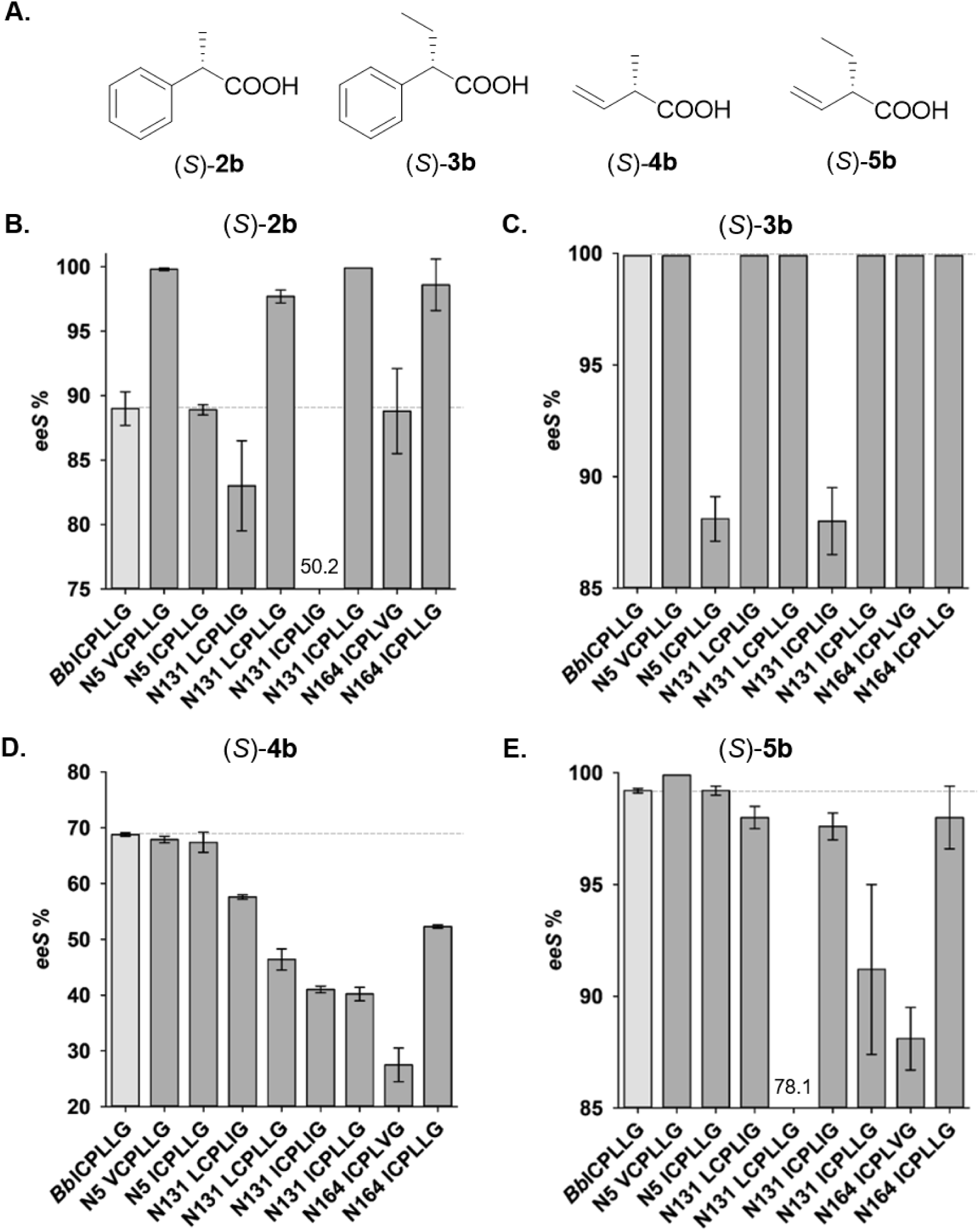
**A.** Chiral products obtained by engineered (*S*)-selective AMDase variants. **B. – E.** Stereoselectivity of engineered (*S*)-selective AMDase variants after the decarboxylation of substrates **2a**-**5a**. Reactions were performed at 30 °C using a 5 mM substrate concentration for **2a** and **3a** and 2.5 mM for **4a** and **5a**. Chiral aryl carboxylic acids (**2b** and **3b**) were derivatized to their corresponding methyl esters prior to chiral GC analysis. All reactions were performed in triplicate, and the stereoselectivity was determined by chiral GC.

In order to investigate the stereoselectivity towards a substrate with two substituents of very similar size, the acceptance of the cyclic **6a** by the ancestors as well as *Bb*AMDase was tested, using cell-free extract (4–8 mg·mL^−1^ total protein). The *ee* was only determined for ancestors that were able to reach full conversion within 20 h of reaction time (**Table S2**). Interestingly, ancestors N5, N10, and N131 converted **6a** notably faster than *Bb*AMDase, the latter leading to incomplete conversion after 16 h. The results of the decarboxylation of **5a** and **6a** show that the ancestors show a similar limitation as *Bb*AMDase towards the size of the smaller α-substituent. An optical purity of about 89–90% *ee* (+)-**6b** was reached for all tested variants.^49^

For the (*S*)-selective ancestors, the stereoselectivity in the conversion of the cyclic compound **6a** towards **6b** is moderate when carried out by AMDase ICPLLG, reaching 91.0% *ee*. All ancestral variants showed lower stereoselectivity, ranging from 2.6 to 84.5% *ee* (-)-**6b** (**Table S2**).^49^

#### Mechanistic study using pseudochiral isotope-labeled 4a

The enhancement of stereoselectivity of some of the AMDase ancestors is an unusual result, as ancestors are generally considered to show a broader substrate acceptance and higher K_M_ values, which often correlate with reduced stereoselectivity.^50,51^ In the decarboxylation of prochiral aromatic malonic acids, extant *Bb*AMDase cleaves exclusively the *pro-R* carboxylic group and forms only the (*R*)-enantiomers of the product.^52–54^ Interestingly, with 2-methyl-2-vinyl malonic acid (**4a**), formation of both enantiomers is observed.^12^ In previous work, we could show with the decarboxylation of a *pro*-*R*-^13^C-labelled **4a** that two stereochemical pathways are possible for this substrate: Cleavage of the *pro*-*R* carboxylate forms the (*R*)-product, and decarboxylation of the *pro*-*S* group forms the (*S*)-enantiomer.^12^ Both pathways proceed with inversion of the absolute configuration at the stereocenter. For decarboxylation of **4a**, a binding mode has been proposed with the *pro*-*S* carboxylate in the dioxyanion hole and the *pro*-*R* group in the hydrophobic pocket, where unfavorable interactions favor decarboxylation (**Figure 10A**).^55^ An inverse binding mode then leads to *pro*-*S* decarboxylation (**Figure 10B**).

**Figure 10:**
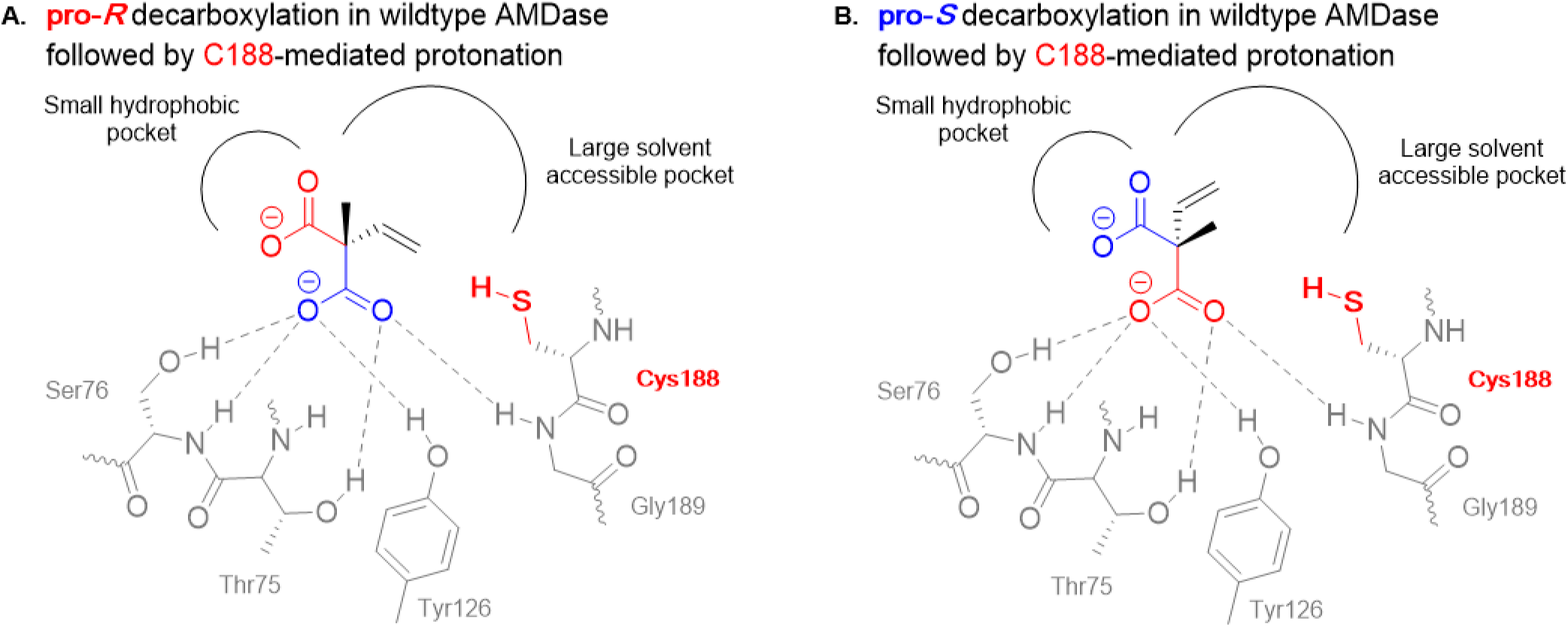
Alternative binding modes of **4a** in arylmalonate decarboxylase leading to cleavage of the *pro*-*R* (**A**) and *pro*-*S* (**B**) carboxylate groups (amino acid numbers correspond to *Bb*AMDase).

We investigated the decarboxylation of the isotope-labeled probe **4a** by ancestors N31 (99.7 *eeR*) and N131 (97.7 *eeR*). In chiral GC-MS analysis, we observed that for both ancestors, formation of the (*R*)-enantiomer resulted from cleavage of the ^13^C-labeled *pro-R* carboxylate (M = 85) (**Figures S41 and S44**). For the (*S*)-enantiomers, the (*S*)-peaks showed a higher intensity of the *m/z* 86 (M + 1) signal than that at *m/z* 85 (M), indicating loss of the unlabeled *pro-S* carboxylate (**Figures S40 and S43**). N31 showed significantly less formation of the (*S*)-enantiomer during decarboxylation of the labeled malonic acid than N131, demonstrating that the decarboxylation proceeds almost exclusively by *pro*-*R* decarboxylation. This is consistent with the higher *ee* observed. These results align with our earlier study, indicating that decarboxylation of **4a** by AMDase ancestors proceeds via the same stereochemical pathway as that of *Bb*AMDase.^12^ Lind et al. simulated both binding modes in *Bb*AMDase with QM simulations in a truncated system.^55^ For **2a**, they found an energy difference of 16.2 kcal·mol⁻¹ between the rate-limiting transition states of both pathways, explaining why only cleavage of the *pro-R* group has been observed.^53,54^ For **4a**, the calculated energy difference of 1.7 kcal·mol⁻¹ was much smaller. The observed 96.8% *ee* of (*R*)-**4b** obtained with *Bb*AMDase results in an energy difference of (ΔΔ*G*^‡^) 2.5 kcal·mol⁻¹ (**Figure S48**, **Equation S4**), which agrees well with their calculations. In contrast, the 99.7% *ee* obtained with N31 corresponds to a difference of both transition states of 3.9 kcal·mol⁻¹, demonstrating the capacity of this ancestor to suppress efficiently the stereochemical pathway initiated by decarboxylation of the *pro-S* carboxylate group. As the composition of the hydrophobic pocket of *Bb*AMDAse, N131 and N31 are highly similar, we attribute the suppression of the cleavage of the *pro*-*S* carboxylate in the latter to second-sphere amino acids.

#### Preparative scale application

Finally, we demonstrated the practical applicability of the most promising (*R*)-selective AMDase ancestor, N131, in a preparative gram-scale conversion of substrate **2a**. Compared to *Bb*AMDase (**Figure 11**), N131 catalyzed the decarboxylation reaction significantly faster, reaching nearly complete conversion within 31 minutes. This conversion translates to a space-time yield of 5.6 g·L⁻¹·h⁻¹, compared to 3.0 g·L⁻¹·h⁻¹ achieved by *Bb*AMDase. To account for the different initial catalyst loadings, specific productivity was determined. N131 yielded 96.2 g of product per gram of enzyme per hour, representing a 3.4-fold improvement over *Bb*AMDase (28.1 g_product_·g_enzyme_⁻¹·h⁻¹). This is consistent with its higher observed catalytic activity in the reactor (*Bb*AMDase: 14.3 U·mg⁻¹; N131: 25.1 U·mg⁻¹) and the previously noted 633-fold higher total turnover number (TTN). Both reactions yielded the product with excellent optical purity and isolated yields (*Bb*AMDase: 99.5% ee, 96.6%; N131: 99.4% ee, 95.8%). Given the substantially greater thermal stability of N131 relative to *Bb*AMDase, including a 294-fold increase in half-life at 45°C, its combination of high activity and stability makes it a superior biocatalyst. This improvement enables significantly higher product yields per gram of *E. coli* biomass, thereby reducing production costs and improving the overall environmental footprint.

**Figure 11.**
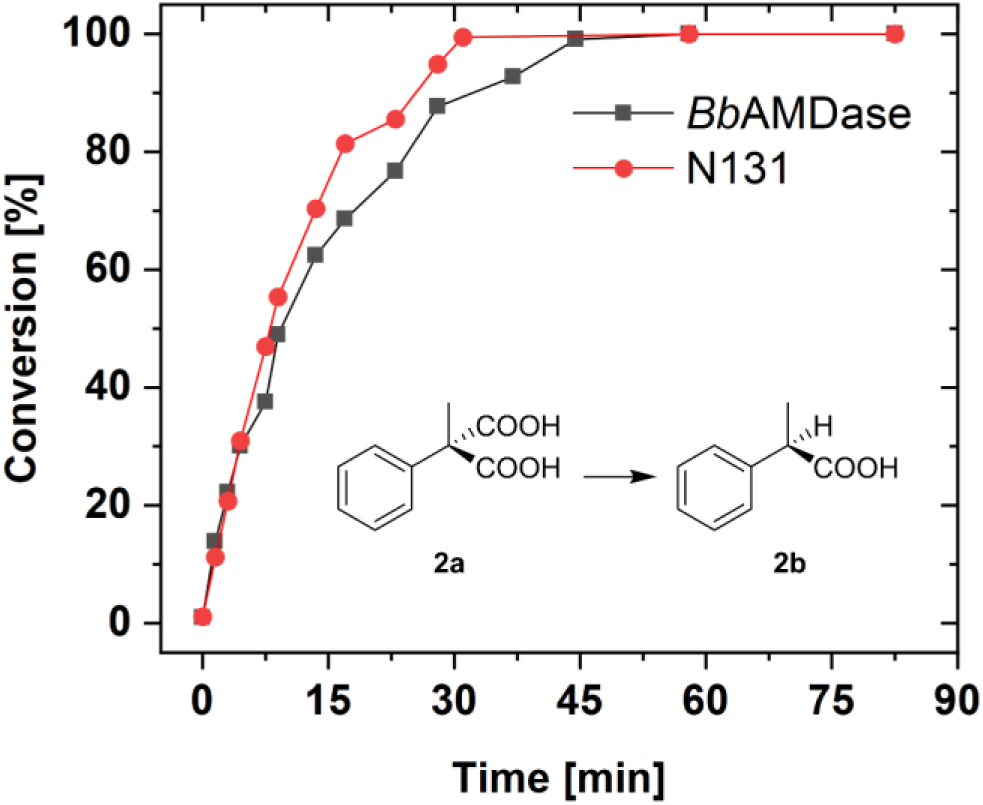
Progressive curve of biocatalytic decarboxylation of 2-methyl-2-phenylmalonate (1.0 g, 20 mM in 50 mM Tris-HCl, pH 8) of *Bb*AMDase and ancestor N131 using CFE (12 mL, *Bb*AMDase: 27.5 mg, N131: 14.9 mg). Conversion was determined using HPLC-DAD. Initial rates of the reactions: *Bb*AMDase: 14.3 U·mg⁻¹, N131: 25.1 U·mg⁻¹, mass of target enzyme determined by densitometric SDS-PAGE (**Figure S10**). Both reactions yielded quantitative isolated yields. The *ee* values of both products were determined using chiral GC (*Bb*AMDase: 99.5% *ee*, N131: 99.4% *ee*). Details shown in the Supporting Information **section 7.14**.

#### Conclusions

In this study, we demonstrate the potential of ASR to increase the catalytic diversity of biocatalysts. Key ancestral enzymes, such as N131 and N31, outperformed the extant *Bb*AMDase in activity, thermal stability, half-life time, and, in one case, stereoselectivity (up to 99.7% *eeR*). Considering an improvement of the thermal stability of N131 of up to +10 °C, the 294-fold improvement of the half-life time is an unusual result that can be rationalized by substantively increased surface polarity of this variant, which creates hydration shielding and enables the formation of temperature-induced salt-bridges that stabilize this ancestor at higher temperatures, while at the same time helping prevent aggregation. Interestingly, while the activity towards aromatic malonic acids was notably increased, the ancestors displayed similar steric limitations to *Bb*AMDase, resulting in low activity toward the cyclic malonate **6a**. Additionally, N131 fully converted α-ethyl malonate **3a**, while *Bb*AMDase achieved only a 15% conversion. Residue swaps at positions G74 and C188 enabled inversion of stereoselectivity, with some (*S*)-selective ancestors surpassing the engineered *Bb*AMDase ICPLLG regarding activity and stereoselectivity. These results highlight ASR as a powerful approach to uncover naturally evolved improvements for more robust and selective biocatalysts.

Furthermore, the ancestral variant N131 demonstrated clear practical advantages in a gram-scale decarboxylation of substrate **2a**, achieving near-complete conversion within 30 minutes, significantly faster than *Bb*AMDase. Both enzymes delivered excellent yields and optical purities, but N131’s higher activity (25.1 vs. 14.3 U·mg⁻¹) and superior stability translate to greater product output per gram of biomass. This makes N131 a more efficient and sustainable biocatalyst for the preparation of optically pure α-chiral carboxylic acids.

Finally, phylogenetic analysis of AMDase offered insights into conserved amino acid patterns and revealed that natural amino acid abundance can inspire enzyme activity optimization campaigns, as key mutations previously identified often appear in the phylogenetic tree. Consequently, knowledge provided by phylogenetic analysis can serve as guidance for further enzyme engineering studies.

## Materials and Methods

### Phylogenetic tree reconstruction

Amino acid sequences of putative AMDase enzymes were acquired via the NCBI database using the protein BLAST tool. The amino acid sequence was retrieved via its Locus Tag using IMG/M database (IMG (doe.gov)). From each cluster, the first 100 hits were taken, and their amino acid sequences were aligned using T-COFFEE Version 11.00 (M-COFFEE). The obtained amino acid sequences were further processed using the MEGAX software. Double sequences and outliers were removed from the collected sequences. We used in-house *R* scripts (*R* package *phangorn* Version 2.12.1) to construct the phylogenetic tree. The tree was pruned to only contain sequences with a maximum of 90% amino acid sequence similarity.

### Enzyme Expression

The cultivation of *E. coli* BL21 (DE3) for the production of AMDase was done by the addition of 10 µL glycerol stock to 10 mL LB-media containing Kanamycin (30 µg/mL), which was incubated overnight (200 rpm, 37 °C). The whole volume of preculture was used to inoculate 0.4 L of LB-media (OD_600_ = 0.1, 30 µg/mL Kanamycin, 120 rpm, 37 °C). The overexpression of AMDase was initiated after ∼2 h (OD_600_ = 0.9) with IPTG (1 mM) and incubated at 28 °C (22 h, 120 rpm). The cells were harvested by centrifugation (4000 rpm, 4 °C, 25 min), and the cell pellet was washed twice with Tris-HCl buffer (20 mL, 50 mM, pH 8), followed by centrifugation (4000 rpm, 4 °C, 25 min) and stored at −20 °C.

### Selectivity Screening

For the screening of selectivity, freshly prepared CFE in Tris-HCl buffer (0 °C, 3.5–5 mg/mL total protein, 50 mM, pH 8) was mixed in equal parts with the respective substrate solution with further addition of CFE after 5 h (300 µL CFE: 300 µL substrate solution + 300 µL CFE, 16 h, 30 °C, 600 rpm). The substrate solution was prepared fresh in Tris-HCl buffer (50 mM, pH 8) with pH adjustment after complete dissolution using 3 M NaOH. For the (*R*)-selective ancestors, the final concentrations of the substrate solutions were: 2-alkyl-2-phenyl-malonic acids: 3.3 mM, 2-methyl-2-vinyl-malonic acid: 1.6 mM, 2-cyclohexene malonate: 1 mM. For the (*S*)-selective ancestors, the final concentrations of the substrate solutions were: 2-alkyl-2-phenyl-malonic acids: 2.5 mM, 2-alkyl-2-vinyl-malonic acid: 2 mM, 2-cyclohexene-1,1-dicarboxylic acid: 2 mM. After complete conversion, the reaction was acidified with HCl (4 M, 200 µL (*R*)-selective and 30 µL (*S*)-selective), and the product was extracted with 300 µL EtOAc and dried with anhydrous MgSO_4_.

### Sequence-Based Analysis

The sequences of wild-type *Bb*AMDase and the N131 ancestor are provided in the Supporting Information. Sequence composition analysis was performed using the ProtParam tool on the ExPasy webserver.^41^ Protein thermostability was predicted from sequence based on propensity scores calculated using the scoring-card method based on the SCMTPP webserver.^42^ Scaled solubility was predicted from sequence using Protein-Sol.^43^

### Molecular Dynamics Simulations and Analysis

Structure modelling of the wild-type and N131 variants was conducted using Chai-1,^47^ based on the available AMDase sequences. Multiple Sequence Alignments (MSAs) were retrieved automatically from the default Chai-1 MSA server. Target representation was augmented with Evolutionary Scale Modelling (ESM) protein language embedding.^56^ The network trunk was executed with 4 recycling iterations to refine structural predictions. To sample conformational diversity, the diffusion module was executed for 200 independent samples, each one generated over 450 denoising timesteps. The per-residue RMSF values for C_α_-atoms was calculated with the CPPTRAJ^57^ module of AmberTools22.^58^

The top conformer from the Chai-1 prediction^47^ was prepared for molecular dynamics simulations. The protonation states of titratable residues (aspartic and glutamic acid, arginine, lysine and histidine residues) was evaluated using PROPKA 3.1,^59,60^ while the tautomeric form of neutral histidine residues was determined with Chimera 1.19.^61^ System preparation was carried out with the tleap module of AmberTools22.^58^ The protein atoms were described using the ff19SB forcefield,^62^ and the system was solvated by a truncated octahedral box filled with OPC (Optimal Point Charge) water molecules,^63^ with a 12.0 Å buffer layer around the protein. The exact number of water molecules was used to calculate the necessary sodium and chloride ions to keep neutrality and approximate 0.15 M NaCl concentration, in accordance with the Split charge protocol.^64^ The systems were then resolvated, applying the same water model and buffer layer, randomly replacing water molecules with the calculated number of ions. All simulations were performed using periodic boundary conditions, with long-range electrostatics treated using the Particle-Mesh Ewald (PME) algorithm.^65^

Following initial system set up, each system was initially minimized using 2,500 steps of steepest descent minimization, followed by 7,500 steps of conjugate gradient minimization, applying a 10 kcal mol^−1^ Å^−2^ positional restraint to all protein heavy atoms. Following this, the systems were heated under canonical (NVT) ensemble conditions from 0 to 100 K over 250 ps simulation time with 1 fs timescales, followed by heating to 300 K during a 250 ps simulation, under isothermal– isobaric (NPT) conditions at 1 bar, maintaining the initial 10 kcal mol^−1^ Å^−2^ heavy-atom restraint. A second cycle of minimization and two-step heating (0–100 K NVT, 100–300 K NPT) was performed under identical conditions, with the positional restraint reduced to 10 kcal mol^−1^ Å^−2^ and restricted only to backbone C_α_-atoms. Equilibration was performed in an NPT ensemble at 300 K over six consecutive 100 ps stages, during which the C_α_-atom restraints were decreased by 1 kcal mol^−1^ Å^−2^ per stage (from 5 to 1 kcal mol^−1^ Å^−2^), with a final 100 ps unrestrained equilibration. Following, unrestrained NPT production simulations were performed at 300 K and 1 bar for 250 ns using a 2-fs time step, saving coordinates every 50 ps. Temperature was maintained using a Langevin thermostat (collision frequency of 1 ps^−1^),^66^ pressure was controlled using a Berendsen barostat (pressure relaxation time of 1 ps^−1^)^67^ and hydrogen bonds were constrained using the SHAKE algorithm.^68^ For higher temperature simulations, instead of 300 K, 360 K was applied as the target temperature of heating, and carrying out the production run. The entire molecular dynamics protocol was carried out with 10 x 0.5 μs independent replicates for each system (10 μs total simulation time over all systems). Convergence of our simulations is shown in **Figure S54**. All simulations were performed using the GPU-accelerated version of the pmemd module of Amber22.^69^ All raw data necessary to reproduce our simulations has been deposited to Zenodo, DOI: 10.5281/zenodo.21894315

Analysis of root mean square deviation (RMSD) values, hydrogen bond interactions, solvent accessible surface areas (SASA), and radial distribution functions (RDF) were performed using the CPPTRAJ^57,70^ module of AmberTools22.^58^ SASA analysis was carried out with the “surf” command, for total SASA, the entirety of the protein was included, while for hydrophobic and hydrophilic SASA values, specific masks were applied, covering hydrophobic and hydrophilic atoms, respectively. Hydrogen bonds were determined with the “hbond” command of CPPTRAJ.^57^ Radial distribution functions were calculated between water oxygen atoms and the carboxylate oxygen atoms of acidic side chains (Asp and Glu residues), with a cutoff distance of 10.0 Å and a resolution of 0.05 Å. Salt bridges and hydrogen bonds were analyzed using MDAnalysis,^71^ applying a 4.0 Å cutoff for the definition of interaction between the terminal oxygen and nitrogen atoms of aspartic and glutamic acid and lysine and arginine residues, respectively, and a 4.5 Å cutoff for defining interaction presence between sidechain carbon atoms of alanine, valine, leucine, isoleucine, methionine, phenylalanine, tryptophane and proline residues.

Time-averaged electrostatic surface analysis was carried out by subsampling the trajectories, aligning every 100^th^ frame to the first frame of the first replica with CPPTRAJ,^57,70^ based on C_α_-atoms (in total, 1000 frames). The frames were saved as individual pdb files, and converted to pqr files with the PDB2PQR software,^72–75^ applying AMBER force field parameters. Electrostatic potential distributions were determined with the Adaptive Poisson-Boltzmann Solver (APBS).^76^ Calculations were carried out on the respective temperature, with an implicit solvent model simulating an ionic strength of 0.15 M NaCl. The resulting per-frame values were averaged, and the surface was depicted in PyMOL.^77^

## ASSOCIATED CONTENT

### Supporting Information

Experimental details, chiral GC chromatograms, and ASR details can be found in the Supporting Information. Raw data to support our simulation analysis have been deposited to Zenodo, DOI: 10.5281/zenodo.21894315, under a CC-BY license.

## AUTHOR INFORMATION

### Author Contributions

The manuscript was written through contributions of all authors. All authors have given approval to the final version of the manuscript. # These authors contributed equally.

## ACKNOWLEDGMENT

We would like to thank Dr. Bernhard Loll, FU Berlin, for fruitful discussions on protein crystallization. We would like to thank the laboratory technician Martina Weiß, from the Institute of Technical Chemistry TCI (Leibniz University Hannover) for her kind technical help in analytics.

