## Supplemental Information for "Ancestors of Arylmalonate Decarboxylase show increased Activity, Stability and Stereoselectivity"

#shared contribution

*Correspondence to:

### 1. ADDITIONAL FIGURES AND TABLES

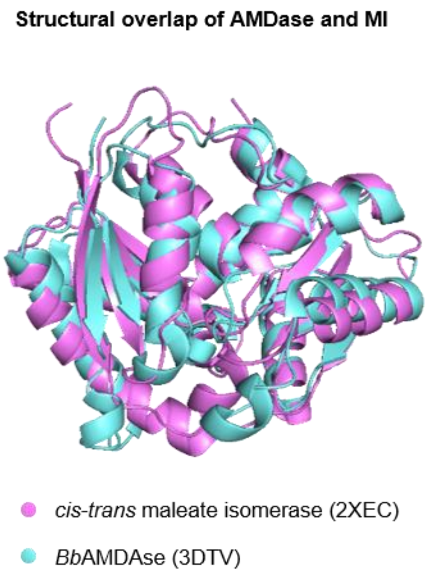

***Figure S1.*** *Overlay of crystal structures of a monomer of Nocardia farcinica maleate cis-trans isomerase (pink) and Bordetella bronchiseptica arylmalonate decarboxylase (blue). PDB entries: 2XEC and 3DTV, respectively. Both enzymes do not have bound ligands and are, therefore, in their open conformation.*

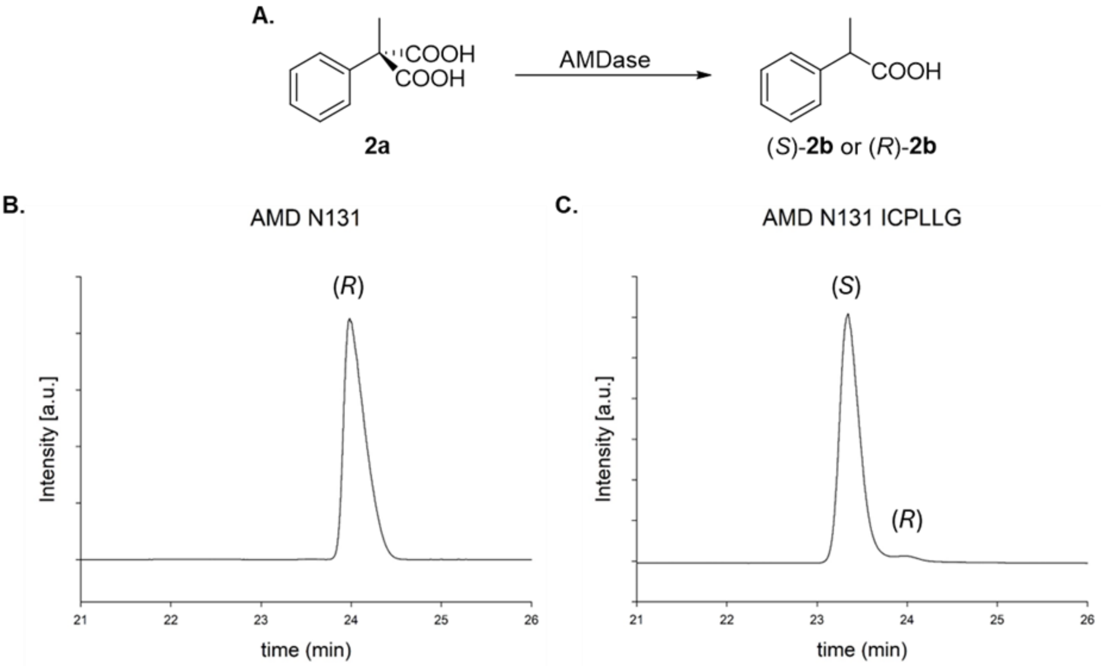

***Figure S2. A.*** *Reaction scheme of AMDase converting **2a** into (R)-**2b** or (S)-**2b**. **B.** and **C.** Chiral GC chromatograms of (R)-**2b** and (S)-**2b** produced by AMDase N131 and AMDase N131 ICPLLG, respectively.*

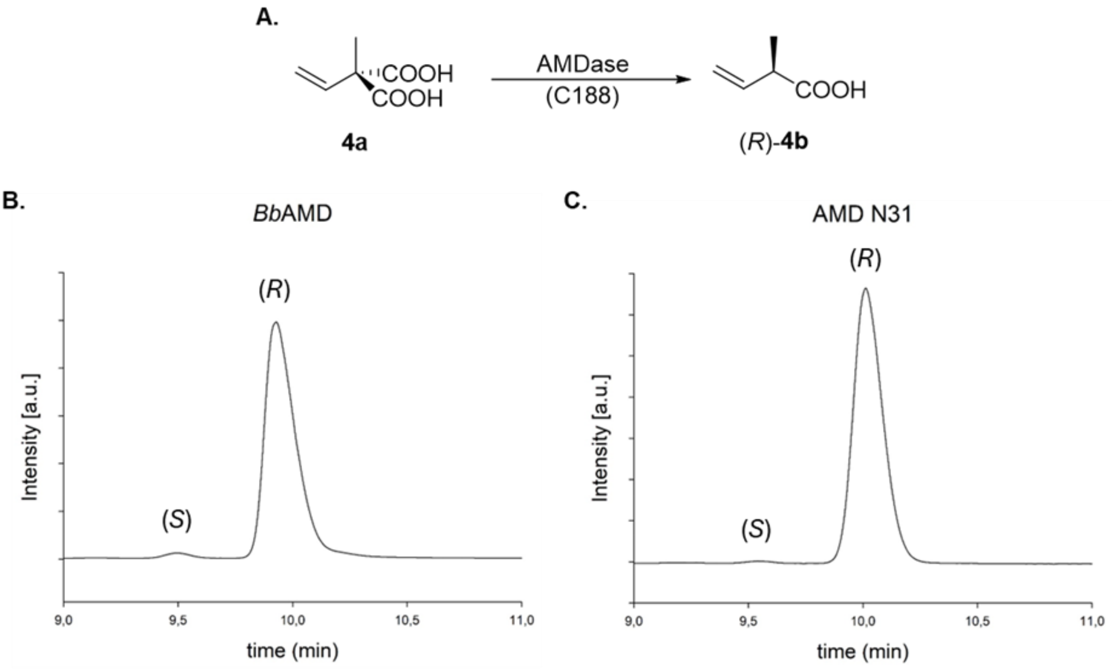

***Figure S3. A.*** *Reaction scheme of AMDase converting 2-methyl-2-vinyl malonate (**4a**) into (R)-2-methyl-3-butenoic acid ((R)-**4b**). **B.** and **C.** Chiral GC chromatograms of **4b** produced by BbAMDase and AMD N31, respectively.*

***Table S1.*** *Conversion of **3a** (5 mM) by (R)-selective AMDase ancestors. The reactions were performed at 30 °C, and the conversions were measured after 16 h.*

| **AMDase** | **Conversion (%)[a]** |
|---|---|
| *Bb*AMDase | 15 |
| N5 | 26 |
| N10 | 18 |
| N31 | 11 |
| N87 | 94 |
| N107 | 27 |
| N109 | 23 |
| N131 | >99 |
| N151[b] | n.d. |
| N164 | 31 |
| N200 | 42 |

[a] determined by HPLC; [b] AMDase N151 showed batch-to-batch deviations.

***Table S2.*** *Decarboxylation of 2-cyclohexene-1,1-dicarboxylic acid (**6a**) by predicted ancestors of AMDase. The reactions were performed at 30 °C, in triplicate.*

| AMDase | Substrate | Sub. conc. (mM) | Reaction time (h) | Product[1] | Conversion (%)[a] | Enantiomeric excess (%)[b] |
|---|---|---|---|---|---|---|
| *Bb*AMDase[c] | **6a** | 1 | 29 | (+)-**6b** | >99 | 89.1 ± 0.5 |
| N5 | **6a** | 1 | 16 | (+)-**6b** | >99 | 89.5 ± 0.0 |
| N10 | **6a** | 1 | 16 | (+)-**6b** | >99 | 89.6 ± 0.4 |
| N31 | **6a** | 2 | 20 | **6b** | 57 | n.d. |
| N87 | **6a** | 2 | 20 | **6b** | 28 | n.d. |
| N107 | **6a** | 2 | 20 | **6b** | 78 | n.d. |
| N109 | **6a** | 2 | 20 | **6b** | 56 | n.d. |
| N131 | **6a** | 1 | 16 | (+)-**6b** | >99 | 89.7 ± 0.7 |
| N151 | **6a** | 2 | 20 | **6b** | 19 | n.d. |
| N164 | **6a** | 2 | 20 | **6b** | 61 | n.d. |
| N200 | **6a** | 2 | 20 | **6b** | 78 | n.d. |
| *Bb*ICPLLG | **6a** | 2 | 24 | (-)-**6b** | >99 | 91.0 ± 3.0 |
| N5 VCPLLG | **6a** | 2 | 24 | (-)-**6b** | >99 | 75.8 ± 0.8 |
| N5 ICPLLG | **6a** | 2 | 24 | (-)-**6b** | >99 | 82.1 ± 1.0 |
| N131 LCPLIG | **6a** | 2 | 24 | (-)-**6b** | >99 | 51.4 ± 2.6 |
| N131 LCPLLG | **6a** | 2 | 24 | (-)-**6b** | >99 | 2.6 ± 1.1 |
| N131 ICPLIG | **6a** | 2 | 24 | (-)-**6b** | >99 | 84.5 ± 1.3 |
| N131 ICPLLG | **6a** | 2 | 24 | (-)-**6b** | >99 | 44.2 ± 3.5 |
| N164 ICPLVG | **6a** | 2 | 24 | (-)-**6b** | >99 | 31.2 ± 2.6 |
| N164 ICPLLG | **6a** | 2 | 24 | (-)-**6b** | >99 | 70.8 ± 2.3 |

[a] determined by HPLC; [b] determined by chiral GC; [c] In order to achieve full conversion and avoid interference of the substrate in the chiral analytics, additional *Bb*AMDase to a total of 7.6 mg protein CFE were added during the course of the reaction. n.d.: not determined.

#### 2. GENERAL INFORMATION

The following *R* packages were used to generate the phylogenetic AMDase tree reconstruction and ASR: *R phangorn* for processing the database and generating the phylogenetic tree, *R* package *APE*, and *phytools* for visualization of the tree, and *R* package *ggmsa* was used to show the HMM plots.[2,3,4]

##### 2.1 Chemicals

All commercially available chemicals and solvents were purchased from Acros Organics, Alfa Aesar, Fisher, Fluka, Merck, Roth, Sigma-Aldrich, TCI, VWR, and used without further purification, unless otherwise stated.

##### 2.2 Strains

The following strains have been used for cloning experiments and production of recombinant proteins, respectively; *E. coli* TOP10, genotype: F- mcrA Δ(mrr-hsdRMS-mcrBC) Φ80(lacZ)ΔM15 ΔlacX74 recA1 araD139 Δ(ara-leu)7697 galU galK rpsL(StrR) endA1 nupG and *E. coli* BL21(DE3), genotype: F- ompT hsdSB ($r_B^- m_B^-$ ) gal dcm (DE3). Both strains have been purchased from Thermo Scientific.

##### 2.3 Isolation of plasmid DNA

Cell material for plasmid isolation was either obtained from liquid culture (5–10 mL overnight culture, ONC) by centrifugation or from densely grown LB-agar plates. Commercially available GeneJET Plasmid Miniprep Kit (Thermo Scientific) or Wizard Plus SV Minipreps DNA Purification System (Promega) was used to isolate the plasmid DNA according to the manufacturer's instructions. Plasmid DNA was typically eluted in pure dd$H_2O$.

##### 2.4 Determination of DNA concentration

The concentration of isolated plasmid DNA or purified PCR products was determined spectrophotometrically with a NanoDrop 2000c spectrophotometer (Peqlab) at 260 nm. Additionally, the absorption at 280 and 230 nm was measured in parallel to assess purity of the DNA sample.

##### 2.5 DNA sequencing and sequence analysis

Samples for Sanger sequencing were prepared according to the instructions provided by the company and sent either to GATC Biotech AG or Microsynth Austria GmbH. Respective sequencing primers were either available from the in-house primer collection or provided by the sequencing company. DNA sequences were aligned and analyzed with the online-tool Benchling.

#### 2.6 Buffer preparation

Reaction buffer (Tris HCl, 50 mM, pH 8): For the reaction buffer, 6.06 g Tris was dissolved in 1 L $ddH_2O$ and acidified with 4 M HCl to pH 8.

Binding buffer (Tris HCl (20 mM), NaCl (300 mM), Imidazole (20 mM, pH 7.4): For the binding buffer, NaCl (17.52 g) and imidazole (1.36 g) were dissolved in 930 mL $ddH_2O$. After that, 20 mL of Tris HCl (1 M stock, pH 8.5) was added, and the solution was acidified to pH 7.4 with 4 M HCl and filled up to 1000 mL. The buffer was filtered under vacuum through a cellulose acetate filter (0.2 µm, Sartorius Stedim Biotech GmbH).

Elution buffer (Tris HCl (20 mM), NaCl (300 mM), Imidazole (300 mM), pH 7.4): For the elution buffer, NaCl (8.76 g) and imidazole (10.20 g) were dissolved in 440 mL $ddH_2O$. After that, 10 mL of Tris HCl (1 M stock, pH 8.5) was added, and the solution was acidified to pH 7.4 with 4 M HCl and filled up to 500 mL. The buffer was filtered under vacuum through a cellulose acetate filter (0.2 µm, Sartorius Stedim Biotech GmbH).

Reverse-phase HPLC mobile phase ($H_3PO_4$, 20 mM): Concentrated Phosphoric acid (2.32 mL) was diluted in 2 L $ddH_2O$ and filtered under vacuum through a cellulose acetate filter (0.2 µm, Sartorius Stedim Biotech GmbH).

#### 2.7 Transformation of competent *E. coli* cells

For the transformation, an aliquot of competent cells (50 µL) was thawed on ice, and 2 µL of plasmid DNA was added to the cell suspension. The mixture was incubated on ice for 30 min, before a heat shock at 42 °C for 30 s was applied. LB-SOC medium (400 µL) was added, and the mixture was regenerated at 37 °C and 300 rpm for 1 h. Suitable aliquots were spread on antibiotic-containing LB agar plates as required and incubated at 37 °C overnight.

#### 2.8 Glycerol stocks

Cultures were maintained in 25% (v/v) glycerol stocks at -20 °C and -80 °C. For this purpose, 50% (v/v) sterile glycerol was thoroughly mixed with the respective overnight culture in a 1:1 ratio in cryogenic tubes. Afterwards, glycerol stocks were frozen and replaced after repeated use.

### 3. ANALYTICAL METHODS

#### 3.1 Gas chromatography

Chiral GC samples were measured on a Shimadzu Nexis GC-2030 device with flame-ionization detector (FID) equipped with an AOC-20i/s 208 autosampler using a Macherey-Nagel cyclodextrin column (Hydrodex-β-6TBDM. Chiral stationary phase, 25 m, 0.25 mm ID, 0.25 μm df). An isothermal method was developed for each product to achieve baseline separation.

**SPL1**: Injection volume: 1 μL, Injection mode: split, Injection temperature: 250 °C, Carrier Gas: $N_2$, Flow control mode: Linear velocity, Linear Velocity: 36 cm/s; **FID**: Sampling Rate 40 ms, Makeup Gas: $N_2$, Detection Temperature: 230 °C, Makeup Flow 24.0 mL/min, $H_2$ Flow: 32.0 mL/min, Air Flow 200.0 mL/min;

**Phenylpropionic acid: SPL1**: Linear Velocity: 20 cm/s **Column:** Temperature: 100 °C, Hold Time: 35 min.

**Phenylbutyric acid: SPL1**: Linear Velocity: 20 cm/s **Column:** Temperature: 103 °C, Hold Time: 35 min Sampling Rate 80ms, Heating rate: 15 °C/min up to 190 °C, hold 1 min.

**2-methylbut-3-enoic acid/ 2-ethylbut-3-enoic acid: Column:** Temperature: 100 °C, Hold Time: 30 min.

**2-cyclohexene acetic acid:** Injection volume: 1 μL (*S*)-selective ancestors and 3 μL (*R*)-selective, **Column:** Temperature: 159 °C, Hold Time: 10 min.

#### 3.2 Gas-chromatography-mass spectrometry

Chiral GC-MS analyses were performed on a GC-2010 plus Shimadzu with an AOC-20s autosampler and an AOC-20i auto injector. The separations were carried out on a Hydrodex-β-6TBDM column, (Chiral stationary phase, 25 m, 0.25 mm ID, 0.25 μm df). The following isothermal method was used: Chiral_GC-MS_M1: Isothermal 100 °C for 30 min.

**SPL1**: Injection volume: 2 μL, Injection mode: split, Injection temperature: 230 °C, Split ratio: 10, Carrier Gas: He, Flow control mode: Linear velocity, Linear Velocity: 47.4 cm/s, Pressure: 83.8 kPa, Total flow: 17.9 mL/min, Column flow: 1.35 mL/min, Purge Flow: 3 mL/min; **MS**: Ion Source temperature: 200°C, Interface temperature: 200°C, Solvent Cut time: 4.01 min, Event time: 0.13.

#### 3.3 High Performance Liquid Chromatography

Analytical HPLC measurements were performed on a Shimadzu HPLC SLC-40. The device was equipped with the following detectors: PDA (SDP-M40) and SPD-40 UV-Vis. The PDA detector was used for the (*R*)-selective ancestors, and SPD-40 UV-Vis detector for the (*S*)-selective ancestors. All separations were carried out on a reversed-phase EC 250/4.6 NUCLEODUR C18 Pyramid (5 μm, particle size: 5.0 μm, length: 250 mm, internal diameter: 4.6 mm) column. The following method was used for the initial rates with CFE on phenylmalonic acid and 2-methyl-2-phenylmalonate with the (*R*)-selective ancestors: Isocratic (20 mM $H_3PO_4$ aq./ACN, 80/20, 1 mL flow rate, 40 °C column temperature, 18 min). The following method was used for the initial rates with CFE on phenylmalonic acid with the (*S*)-selective ancestors: Isocratic (20 mM $H_3PO_4$ aq./ACN, 80/20, 1 mL flow rate, 25 °C column temperature, 25 min). For the initial rates with ancestor N31 with 2-methyl-2-phenylmalonate the following method was used: Isocratic (20 mM $H_3PO_4$ aq./ACN, 70/30, 1 mL flow rate, 40 °C column temperature, 25 min). For the initial rates with ancestor N31 with 2-methyl-2-vinylmalonic acid the following method was used: Isocratic (20 mM $H_3PO_4$ aq./ACN, 85/15, 1 mL flow rate, 40 °C column temperature, 20 min). The conversion data of 2-alkyl-2-phenylmalonates was obtained at 254nm, for 2-alkyl-2-vinyl malonic acids at 210nm and 2-cyclohexene malonate at 190 nm.

Analytical HPLC measurements for the kinetic– and half-life characterization of AMDase ancestor N131 were carried out using a VWR® Hitachi Chromaster equipped with an diode-array detector (Product no. 5430). The product concentration was quantified at 210 nm after separation  by a Kinetex® 2.6 µm Polar C18 (100 Å, inner diameter: 4.6 mm and length: 150 mm) column. A flow rate of 1 mL/min was used and a gradient of $H_3PO_4$ (20 mM) in dd$H_2O$ and acetonitrile (95:5 → 20:80%, 15 min).

#### 4. SEQUENCE DATABASE

Amino acid sequences of putative AMDase enzymes were acquired via the NCBI database using the protein BLAST tool. From each AMDase cluster described by Maimanakos *et al.*[5], one AMDase representative sequence (***Table S1***) was arbitrarily chosen and used for the BLAST function. The amino acid sequence was retrieved via its Locus Tag using IMG/M database (IMG (doe.gov)). From each cluster, the first 100 hits were taken, and their amino acid sequences were aligned using T-COFFEE Version 11.00 (M-COFFEE).

***Table S1:*** *Amino acid sequences from each cluster*[5] *and used for the BLAST tool on the NCBI database.*

| Cluster | Locus Tag (IMG/NCBI) | Organism | Isolation source |
|---|---|---|---|
| **ABC-I** | G489DRAFT_03382 | *Cucumibacter marinus* DSM 18995 | Costal seawater |
| **TRAP-I** | H610DRAFT_00387 | *Pseudomonas azotifigens* DSM 17556 | Hyperthermal compost material |
| **TRAP-II** | NA2_08029 | *Nitratireductor pacificus* pht-3B | Pyrene-degrading consortium of an enriched sediment from the Pacific Ocean |
| **TTT-I** | Ga0077226_10648 | *Bordetella bronchiseptica* KU1201 | Soil enriched with phenylmalonate as sole carbon source |
| **TTT-II** | G456DRAFT_02398 | *Alcaligenes faecalis phenolicus* DSM 16503 | Wastewater bioprocessor |
| **TTT-III** | Ga0058957_03218 | *Variovorax* sp. NFACC27 | Bacterial root endophytes of switchgrass |
| **TTT-IV** | AmiJ41DRAFT_00035970 | *Aminobacter* sp. J41 | Lignin-degrading bacterial isolates |
| **TTT-V** | C507DRAFT_01657 | *Amorphus coralli* DSM 19760 | Coral mucus, host *Pleuractis granulosa* |

The obtained amino acid sequences were further processed using the MEGAX software. Double sequences and outliers were removed from the collected sequences. We used in-house *R* scripts to have sequences with at least four amino acid differences. These remaining 222 AMDase sequences were used to construct the phylogenetic tree.

Four sequences from Maleate *cis-trans* Isomerase (MI) published by F. Fisch et al.[6] (accession numbers: Q5YXQ1, Q9KWI0, O24766, and Q9WX57) were used to generate an outgroup.

To further process the data and generate the phylogenetic AMDase tree, the sequence database was transferred to the *R phangorn* software. ***Figure S4*** depicts the aligned AMDase (222) and Maleate *cis-trans* isomerase (4) amino acid sequences. AMDase and MI share the same fold, and the catalytic residue C188 and also G74 (being the catalytic residue in the mutant G74C of AMDase and its derivatives) on the opposite side of the active center, and the two catalytic Cys residues C76 and C190 of MI are structurally well-conserved. Therefore, these residues were aligned for the phylogenetic analysis.

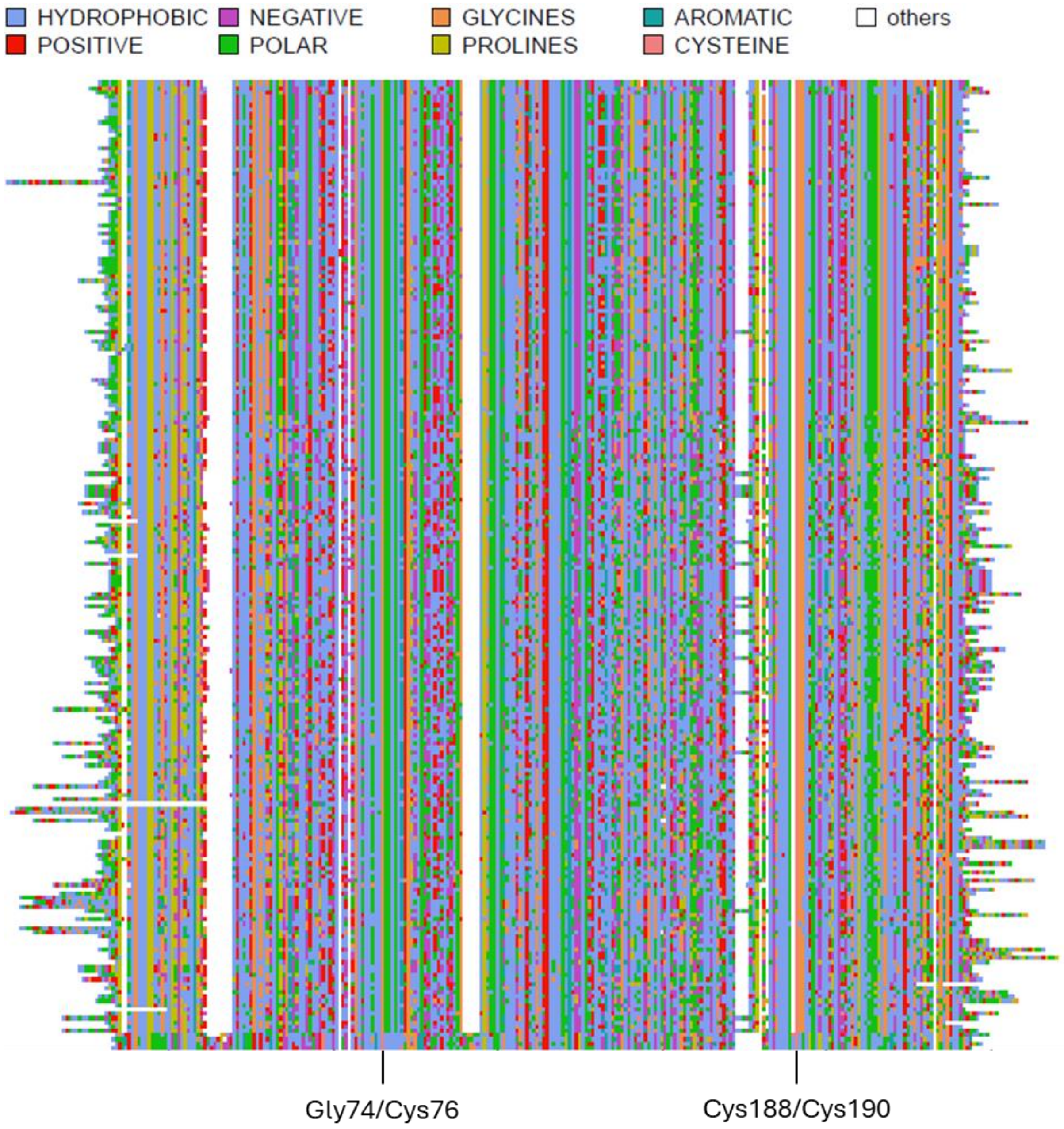

***Figure S4.*** *Aligned AMDase and MI amino acid sequences that were used to generate the rooted phylogenetic tree. The bottom four sequences indicate the MI outgroup. The alignment is shown based on the properties of the amino acid site residues. Gly74 (AMDase) and Cys76 (MI) were aligned, as well as Cys188 (AMDase) and Cys190 (MI). The white "other" indicates a gap state.*

#### 5. PHYLOGENETIC TREE

We chose WAG+G(4)+I as a transition model after performing model tests to derive the best-fitting amino acid model.[7] We coded gaps as their own state, not as an ambiguous state, and performed stochastic rearrangements using 100 iterations to optimize the topology. We rooted the tree using the 4 outgroup sequences. The resulting phylogenetic tree was employed to select ancestors. We chose 10 nodes (N5, N10, N31, N87, N107, N109, N131, N151, N164, and N200), so that all major clades were covered, see ***Figure S5***.

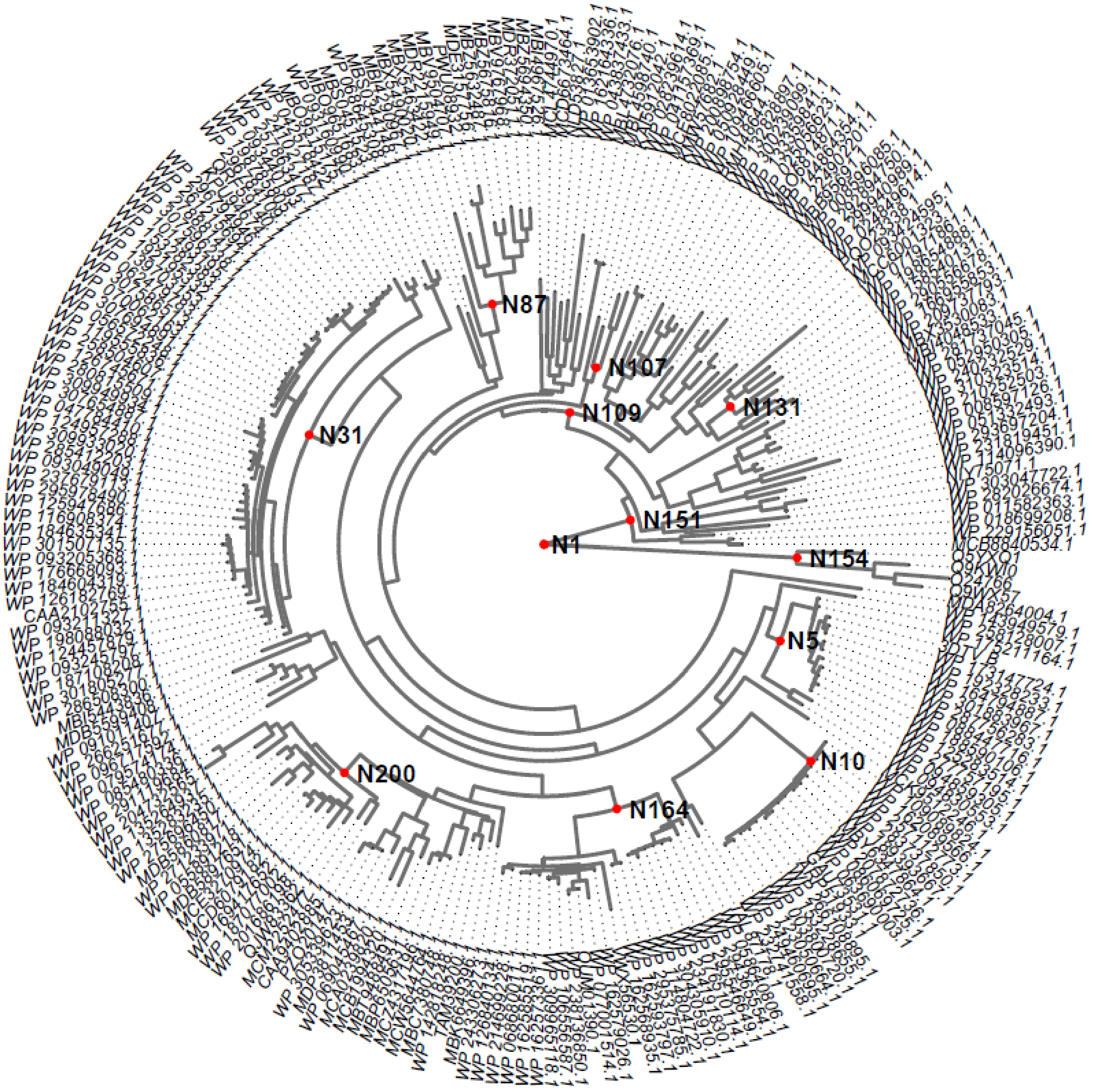

***Figure S5.*** *Rooted maximum-likelihood (ML) phylogenetic AMDase tree based on 222 AMDase and 4 MI sequences.*

We performed marginal ancestral sequence reconstruction to get the sequences of the nodes with the *R* package *Phangorn*.[2] Marginal sequence reconstruction gives a probability for each state (amino acid and gap). As the ancestral state, we chose the one with the highest probability.[8,4]

The tree was further pruned to only contain sequences with a maximum of 90% amino acid sequence similarity, see ***Figure S6***. The scripts prune the tree, gap as a state, and improved ancestral reconstruction have been incorporated in the *R* package *phangorn* Version 2.12.1.[2]

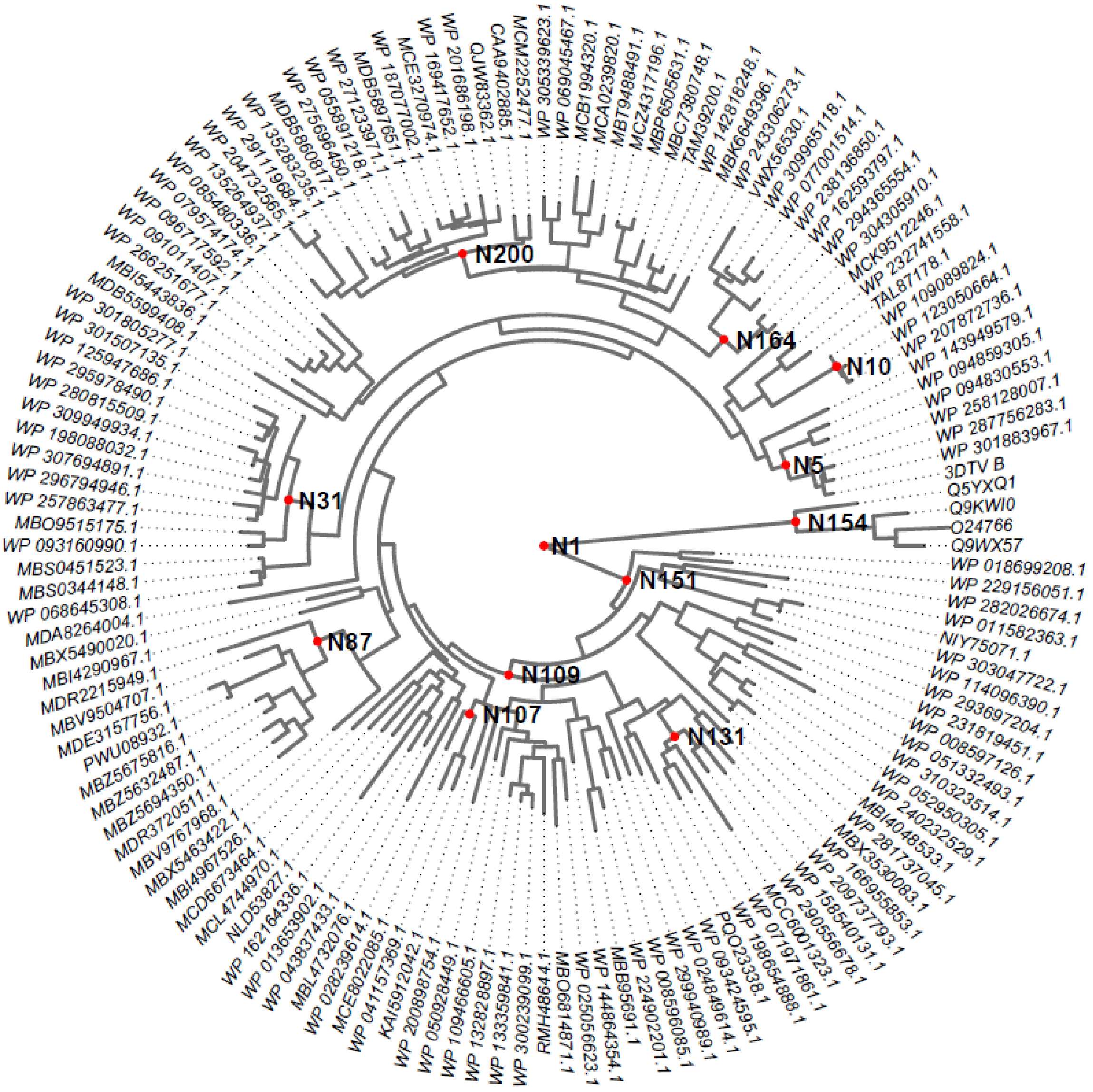

***Figure S6.*** *Rooted maximum-likelihood (ML) phylogenetic AMDase tree based on 136 AMDase and 4 MI sequences.*

***Table S2.*** *Overview of the origin of extant enzymes from early ancestors bearing a higher thermal stability (green) and showing a lower thermal stability (red) than BbAMDase. The majority of the species were isolated from soil, marine sediment, and/or from fresh waters.*

| Ancestor | Based on extant enzymes | Organism | Type of species |
|---|---|---|---|
| **N5** (6 extant enzymes) | 3DTV (*Bb*AMDase) | *Bordetella bronchiseptica* | Bacterium |
| | WP 301883976.1 | *Dialister invisus* | Bacterium |
| | WP 287756283.1 | *Achromobacter sp.* | Bacterium |
| | WP 258128007.1 | *Achromobacter anxifer* | Bacterium |
| | WP 094830553.1 | *Bordetella genomosp. 1* | Bacterium |
| | WP 094859305.1 | *Bordetella genomosp. 5* | Bacterium |
| **N131** (5 extant enzymes) | WP 158540131.1 | *Rhodosalinus halophilus* | Bacterium |
| | WP 290556678.1 | *Aestuariivita sp.* | Bacterium |
| | MCC6001323.1 | *Pararhodobacter sp.* | Bacterium |
| | WP 071971861.1 | *Sulfitobacter alexandrii* | Bacterium |
| | WP 198654888.1 | *Albibacillus kandeliae* | Bacterium |
| **N164** (7 extant enzymes) | WP 304305910.1 | *Pseudacidovorax intermedius* | Bacterium |
| | WP 294365554.1 | *Pseudacidovorax sp.* | Bacterium |
| | WP 162593797.1 | *Variovorax sp. PBL-E5* | Bacterium |
| | WP 238136850.1 | *Variovorax sp. JS1663* | Bacterium |
| | WP 077001514.1 | *Variovorax sp. KK3* | Bacterium |
| | WP 309965118.1 | *Variovorax guangxiensis* | Bacterium |
| | VWX56530.1 | *Burkholderiales bacterium 8X* | Bacterium |
| **N200** (16 extant enzymes) | WP 135264937.1 | *Ramlibacter henchirensis* | Bacterium |
| | WP 204732565.1 | *Hydrogenophaga laconesensis* | Bacterium |
| | WP 291119684.1 | *Hydrogenophaga sp.* | Bacterium |
| | WP 135283235.1 | *Ramlibacter rhizophilus* | Bacterium |
| | MDB5860817.1 | *Ramlibacter sp.* | Bacterium |
| | WP 275696450.1 | *Ramlibacter paludis* | Bacterium |
| | WP 055891218.1 | *Ramlibacter sp. Leaf400* | Bacterium |
| | WP 271233971.1 | *Ramlibacter tataouinensis* | Bacterium |
| | MDB5897651.1 | *Ramlibacter sp.* | Bacterium |
| | WP 187077002.1 | *Ramlibacter cellulosilyticus* | Bacterium |
| | MCE3270974.1 | *Ramlibacter sp.* | Bacterium |
| | WP 169417652.1 | *Ramlibacter agri* | Bacterium |
| | WP 201686198.1 | *Ramlibacter aurantiacus* | Bacterium |
| | QJW83362.1 | *Ramlibacter terrae* | Bacterium |
| | CAA9402885.1 | *uncultured Ramlibacter sp.* | Bacterium |
| | MCM2252477.1 | *Ramlibacter sp.* | Bacterium |

| | | | |
|---|---|---|---|
| **N10** (3 extant enzymes) | WP 109089824.1 | *Alcaligenes faecalis* | Bacterium |
| | WP 123050664.1 | *Alcaligenes faecalis* | Bacterium |
| | WP 207872736.1 | *Alcaligenes* | Bacterium |
| **N31** (12 extant enzymes) | WP 301805277.1 | *Variovorax ginsengisoli* | Bacterium |
| | WP 301507135.1 | *Variovorax sp. CAN15* | Bacterium |
| | WP 125947686.1 | *Variovorax sp.* | Bacterium |
| | WP 295978490.1 | *Variovorax sp.* | Bacterium |
| | WP 280815509.1 | *Variovorax sp. TBS-050B* | Bacterium |
| | WP 309949934.1 | *Variovorax sp.* | Bacterium |
| | WP 198088032.1 | *Variovorax sp. E3* | Bacterium |
| | WP 307694891.1 | *Variovorax boronicumulans* | Bacterium |
| | WP 296794946.1 | *Variovorax sp.* | Bacterium |
| | WP 257863477.1 | *Variovorax sp. ZS18.2.2* | Bacterium |
| | MBO9515175.1 | *Variovorax sp.* | Bacterium |
| | WP 093160990.1 | *Variovorax sp. YR216* | Bacterium |
| **N87** (9 extant enzymes) | MDR2215949.1 | *Nevskiaceae bacterium* | Bacterium |
| | MBV9504707.1 | *Terriglobia bacterium* | Bacterium |
| | MDE3157756.1 | *Acidobacteriota bacterium* | Bacterium |
| | PWU08932.1 | *Terriglobia bacterium* | Bacterium |
| | MBZ5675816.1 | *Terriglobia bacterium* | Bacterium |
| | MBZ5632487.1 | *Terriglobia bacterium* | Bacterium |
| | MBZ5694350.1 | *Terriglobia bacterium* | Bacterium |
| | MDR3720511.1 | *Candidatus Acidiferrales bacterium* | Bacterium |
| | MBV9767968.1 | *Bryobacterales bacterium* | Bacterium |
| **N107** (3 extant enzymes) | WP 028239614.1 | *Stutzerimonas azotifigens* | Bacterium |
| | MCE8022085.1 | *Billgrantia zhangzhouensis* | Bacterium |
| | WP 041157369.1 | *Halomonadaceae* | Bacterium |

#### 6. ANCESTOR SEQUENCES OF ARYLMALONATE DECARBOXYLASE

The bases AUG (coding for methionine) were added to ensure expression of the AMDase ancestors. The following sequences have been ordered from *Twist Bioscience* into the pET-28a(+) vector with a C-terminus His-Tag. The codons were optimized for *E. coli*.

##### 6.1 *(R)*-selective AMDase variants

*Bb*AMDase (from *Bordetella bronchiseptica*)

MGQMQQASTPTIGMIVPPAAGLVPADGARLYPDLPFIASGLGLGSVTPEGYDAVIESVVDHARRLQKQ
GAAVVSLMGTSLSFYRGAAFNAALTVAMREATGLPCTTMSTAVLNGLRALGVRRVALATAYIDDVNERL
AAFLAEESLVPTGCRSLGITGVEAMARVDTATLVDLCVRAFEAAPDSDGILLSCGGLLTLDAIPEVERRL
GVPVVSSSPAGFWDAVRLAGGGAKARPGYGRLFDESGGSHHHHHH*

AMDase_N5

MAPQSSTPVIGLIVPPAAGEVPADGARLYPGLRFIASGLGLGSVTPEGYDAVIDSVVDHARKLADQGAS
AISLMGTSLSFYRGAAFNRALTEAMREATGLPCTTMSHAVLRGLRALGLRRVALATAYIDDVNERLAAF
LAEEGFDVTACRSLGITGVEAMARVSTDTLVDLCERAFEAAPDSDGILLSCGGLLTLDAIPQVERRLGV
PVVSSSPAGFWDVVRLAGHDPQAPGYGRLFGPSHHHHHH*

AMDase_N10

MATTDNSDTKRPVLGLIVPPAAGLVPPEGPEMYPEIDFIAQGLALSSVDKEGYDQVIDQVVDAAQKLAA
RGAQAVSLMGTSLSFYRGSDFNEELVARLRESTGLPCSTMSHAILRGLRASGIERVAVASSYIDDVNQ
RLVRFLAQNQIQAVCAYGLGVNDVTAMSQISTQELVDLCLKTWDIAQEQAPGQAQGLLLSCGGLVSLE
AVRQVEDKLGVTVVSSSPAGFWDLVATAGLDLFPRGMGRLAQERQTAHHHHHH*

AMDase_N31

MATQPHLGLIVPPAAGAVPVDGPLLYGERIRFSARGLGLGEISTRGYTEVIDSVIEKARELAAEGVQAVS
LMGTSLSFFRGAAFNRELQQEMARATGLPCSTMSDAIVGGLRSLGVRRVAVATAYIDEVNRQLRRYLE
DSGFEPLALQGLSISDVQAVGEVPTAVLVDLCRTVFEASPGADGILISCGGLVTLDAVREVEARLGVPV
VSSSPAGFWDLVRCAGLDARSPGHGRLFDQGRHHHHHH*

AMDase_N87

MARGFRRTFLAGAAAGAAGAAAGAAAEPVLGLIFPPANRAVPEEGLAMYPDRVRFLAEGLGLERMTP
EGYDAVIDRIAPAAKKLAEQGAQAISLMGTSLTFYKGAAFNQQLIETVHKATGLPATTMSTGIVEGLKAV
GARRVAVATAYNDEVNERLRAFLEEHGFEVLVVKGLGIEAVEDIPAVTQDELLDFGARVYESAPKADALL
ISCGGLRTLELLAPLEKRCKVPVVSSTPHALWAAVRLVGLSGRVPGYGRLLSRAHHHHHH*

AMDase_N107

MATPNRPTVGLIVPPAAGEVPPEPPALYGDEVNFIAAGLGLRKLTPEGYDAVIDRVGELSRELAADGAD
AVSLMGTSLSFYRGPEFNAQLIEIMQEASGLPATTMSNAVIEALNAVGARRIAVATAYVDSVNQRLADFL
AESGFEVASLESLDIERVEDIAAVTQDELLELGRRALTAAPADALFISCGGLRTQEVTLQLEAEFGIPVIS
SAMAGAWAAVRLVGHSGRAPGSGRLFEAGPHHHHHH*

AMDase_N109
MATRPTIGLIVPPAAGEVPPEAPALYPDRIRFIARGLGLKEMSPEGYDAVIDRVAELARRLAEQGAQAVS
LMGTSLSFYRGPEFNDQLIETMREATGLPATTMSNAIIEALRAVGARRIAVATAYTDEVNQRLRDYLEES
GFEVASLESLGITAVEDVHAVTEDELIDLGRRAFEAAPGADALLISCGGLRTLDVTPPLEARLGIPVISSA
MAGAWAAVRLVGHDGRAPGFGRLFEQAPHHHHHH*

AMDase_N131
MATTPVIGMIVPPAAGEVPPEAYGLYPEGVRFAARGLALKELSMEDYSEAIERVAELARELREEEGADA
ISLMGTSLSFFRGGAFNDELVEIMQQETGVPATTMSNSIRDALRAVGARRIAVGTAYTDEVNDRLRGFL
EASGFEVASLTGMGLTAVEDILAVTEDEVTDLGLRAFEAADGPADAVLISCGGLPALHLADALEPEIGVP
VVASSTAGVWGAVRLLGLSGESPALGRLGRTPPSRESAVGHHHHHH*

AMDase_N151
MATRPTIGLIVPPAAGEVPPEAPAMYPDLRFIARGLGLKEMSPEGYARVIDRVGDAAKRLAARGADVVS
LMGTSLSFFRGPDFNDQLIETMAERSGLPATTMSSAIIEALRAVGARRIAVLTAYEDDVNQRLRDYLEEH
GFEVASLQSLGIRAVEDVAGVSEDQLIDLGRRAFEAAPGADALLISCGGLQTLDVVPPLEAKTGIPVVS
SATAGVWAAVRLVGHDARAPGFGRLFEVAPHHHHHH*

AMDase_N164
MATDSATPTLGLIVPPAAGEVPPDGPALYGGRVRFIARGLGLAGISPEGFDAVVDRILDLARELRDAGA
QAISLMGTSLSFYRGAAFTEDLRARMQEATGLPCTTMSHAIVRSLRQLGIRRVAVATAYIDTLNDRLVAY
LAGEGFEVTAIRGLSITGVEAVGQVPAETLMELAERVVAADPSADGLLISCGGLLTLDLHPPLERRLGLP
VTSSSPAGFWDLMRTAGLDPASPGHGRLFEPARHHHHHH*

AMDase_N200
MAMSSTPCIGLIVPPAAGQVPQDGPMLYGDRVRFIARGLGIGAVSPEGFNPVIDTILDRARELRDAGAQ
VISLMGTSISFYRGAAFTESLRAAMQEATGVPCTTMSHAIVAALRQLGIQRVAVATSYIDELNDRLVDYL
THAGFTVTAIRGMSITGVKEVGEVPTQALVELSKSVYAQDPSADGIFISCGGLLTLDAIRQLEDRLGLPV
TASSPAGFWDVVRLAGWDPSSPGFGRLFAPSHHHHHH*

#### 6.2 *(S)*-selective AMDase variants

*Bb*AMDase_ICPLLG
MGQMQQASTPTIGMIVPPAAGLVPADGARLYPDLPFIASGLGLGSITPEGYDAVIESVVDHARRLQKQG
AAVVSLMCTSLSFYRGAAFNAALTVAMREATGLPCTTMSTAVLNGLRALGVRRVALATPYIDDVNERLA
AFLAEESLVPTGCRSLGITGLEALARVDTATLVDLCVRAFEAAPDSDGILLSGGGLLTLDAIPEVERRLG
VPVVSSSPAGFWDAVRLAGGGAKARPGYGRLFDESGGSHHHHHH*

AMDase_N5_ICPLLG
MAPQSSTPVIGLIVPPAAGEVPADGARLYPGLRFIASGLGLGSITPEGYDAVIDSVVDHARKLADQGAS
AISLMCTSLSFYRGAAFNRALTEAMREATGLPCTTMSHAVLRGLRALGLRRVALATPYIDDVNERLAAF
LAEEGFDVTACRSLGITGLEALARVSTDTLVDLCERAFEAAPDSDGILLSGGGLLTLDAIPQVERRLGVP
VVSSSPAGFWDVVRLAGHDPQAPGYGRLFGPSHHHHHH*

AMDase_N5_VCPLLG
MAPQSSTPVIGLIVPPAAGEVPADGARLYPGLRFIASGLGLGSVTPEGYDAVIDSVVDHARKLADQGAS
AISLMCTSLSFYRGAAFNRALTEAMREATGLPCTTMSHAVLRGLRALGLRRVALATPYIDDVNERLAAF
LAEEGFDVTACRSLGITGLEALARVSTDTLVDLCERAFEAAPDSDGILLSGGGLLTLDAIPQVERRLGVP
VVSSSPAGFWDVVRLAGHDPQAPGYGRLFGPSHHHHHH*

AMD_N131_ICPLIG
MATTPVIGMIVPPAAGEVPPEAYGLYPEGVRFAARGLALKEISMEDYSEAIERVAELARELREEEGADAI
SLMCTSLSFFRGGAFNDELVEIMQQETGVPATTMSNSIRDALRAVGARRIAVGTPYTDEVNDRLRGFL
EASGFEVASLTGMGLTALEDILAVTEDEVTDLGLRAFEAADGPADAVLISGGGLPALHLADALEPEIGVP
VVASSTAGVWGAVRLLGLSGESPALGRLGRTPPSRESAVGHHHHHH*

AMD_N131_LCPLIG
MATTPVIGMIVPPAAGEVPPEAYGLYPEGVRFAARGLALKELSMEDYSEAIERVAELARELREEEGADA
ISLMCTSLSFFRGGAFNDELVEIMQQETGVPATTMSNSIRDALRAVGARRIAVGTPYTDEVNDRLRGFL
EASGFEVASLTGMGLTALEDILAVTEDEVTDLGLRAFEAADGPADAVLISGGGLPALHLADALEPEIGVP
VVASSTAGVWGAVRLLGLSGESPALGRLGRTPPSRESAVGHHHHHH*

AMD_N131_ICPLLG
MATTPVIGMIVPPAAGEVPPEAYGLYPEGVRFAARGLALKEISMEDYSEAIERVAELARELREEEGADAI
SLMCTSLSFFRGGAFNDELVEIMQQETGVPATTMSNSIRDALRAVGARRIAVGTPYTDEVNDRLRGFL
EASGFEVASLTGMGLTALEDLLAVTEDEVTDLGLRAFEAADGPADAVLISGGGLPALHLADALEPEIGVP
VVASSTAGVWGAVRLLGLSGESPALGRLGRTPPSRESAVGHHHHHH*

AMD_N131_LCPLLG
MATTPVIGMIVPPAAGEVPPEAYGLYPEGVRFAARGLALKELSMEDYSEAIERVAELARELREEEGADA
ISLMCTSLSFFRGGAFNDELVEIMQQETGVPATTMSNSIRDALRAVGARRIAVGTPYTDEVNDRLRGFL
EASGFEVASLTGMGLTALEDLLAVTEDEVTDLGLRAFEAADGPADAVLISGGGLPALHLADALEPEIGVP
VVASSTAGVWGAVRLLGLSGESPALGRLGRTPPSRESAVGHHHHHH*

AMD_N164_ICPLLG
MATDSATPTLGLIVPPAAGEVPPDGPALYGGRVRFIARGLGLAGISPEGFDAVVDRILDLARELRDAGA
QAISLMCTSLSFYRGAAFTEDLRARMQEATGLPCTTMSHAIVRSLRQLGIRRVAVATPYIDTLNDRLVAY
LAGEGFEVTAIRGLSITGLEALGQVPAETLMELAERVVAADPSADGLLISGGGLLTLDLHPPLERRLGLP
VTSSSPAGFWDLMRTAGLDPASPGHGRLFEPARHHHHHH*

AMD_N164_ICPLVG
MATDSATPTLGLIVPPAAGEVPPDGPALYGGRVRFIARGLGLAGISPEGFDAVVDRILDLARELRDAGA
QAISLMCTSLSFYRGAAFTEDLRARMQEATGLPCTTMSHAIVRSLRQLGIRRVAVATPYIDTLNDRLVAY
LAGEGFEVTAIRGLSITGLEALGQVPAETLMELAERVVAADPSADGLLISGGGLLTLDLHPPLERRLGLP
VTSSSPAGFWDLMRTAGLDPASPGHGRLFEPARHHHHHH*

#### 7. EXPERIMENTAL PROCEDURES

##### 7.1 Cultivation of *E. coli* for AMDase production

The cultivation of *E. coli* BL21 (DE3) for the production of AMDase was done by the addition of 10 µL glycerol stock to 10 mL LB-media containing Kanamycin (30 µg/mL), which was incubated overnight (200 rpm, 37 °C). The whole volume of preculture was used to inoculate 0.4 L of LB-media ($OD_{600}$ = 0.1, 30 µg/mL Kanamycin, 120 rpm, 37 °C). The overexpression of AMDase was initiated after ~2 h ($OD_{600}$ = 0.9) with IPTG (1 mM) and incubated at 28 °C (22 h, 120 rpm). The cells were harvested by centrifugation (4000 rpm, 4 °C, 25 min), and the cell pellet was washed twice with Tris-HCl buffer (20 mL, 50 mM, pH 8), followed by centrifugation (4000 rpm, 4 °C, 25 min) and stored at -20 °C.

##### 7.2 Preparation of cell-free extract (CFE)

The frozen cell pellet was thawed on ice and suspended in Tris-HCl buffer (0 °C, 75 mg/mL, 50 mM, pH 8) or binding buffer (0 °C, 75 mg/mL, 50 mM Tris-HCl, 300 mM NaCl, 10 mM imidazole, pH 7.4) prior sonication on ice (3 x 2 min cycle, 1 min pause, 50% amplitude, duty cycle 5). The lysed cell suspension was centrifuged (10,000 rpm, 4 °C, 30 min) and placed on ice until use.

##### 7.3 Bromothymol blue assay (BTB-assay)

The Bromothymol blue-assay (BTB-assay) was done by a modified procedure of Okrasa *et. al*.[9] The assay solution was prepared by adding Bromothymol blue (0.5 µg/mL) into a solution of 10 mM phenylmalonic acid (2.5 mM MOPS buffer). The pH was adjusted to approximately 6.9 to a green solution and was split into 300 µL aliquots. The conversion was observed by adding 2 µL of CFE with comparison to the respective buffered blank solution.

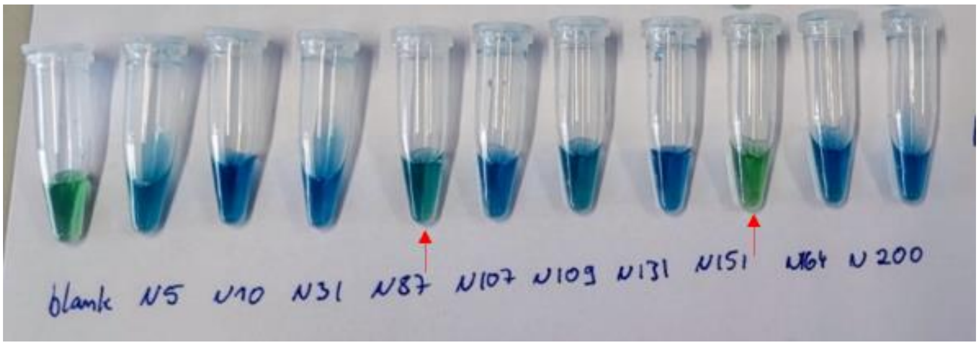

***Figure S7.*** *Bromothymol blue-assay of the AMDase ancestors for direct indication of active enzyme expression.*

##### 7.4 Qualitative observation of inclusion bodies using SDS-PAGE

After sonication, the lysed cells were centrifuged (1 mL sample, 13000 rpm, 4 °C, 15 min), and the total protein concentration of the cell-free extract (CFE) was determined by BCA. The samples were diluted to normalize the protein content for the SDS-PAGE analysis. For the (*R*)-selective ancestors, 10 µg of total protein was loaded per pocket (***Figure S8***), whereas for the (*S*)-selective ancestors, 3.75 µg of total protein was loaded per pocket (***Figure S9***). The remaining pellets of inclusion bodies and cell debris were washed three times with dd$H_2O$ (1 mL, 13000 rpm, 4 °C, 15 min). The hypothetical protein concentration was normalized to the respective sample and incorporated as a suspension in the preparation of the SDS-PAGE sample.

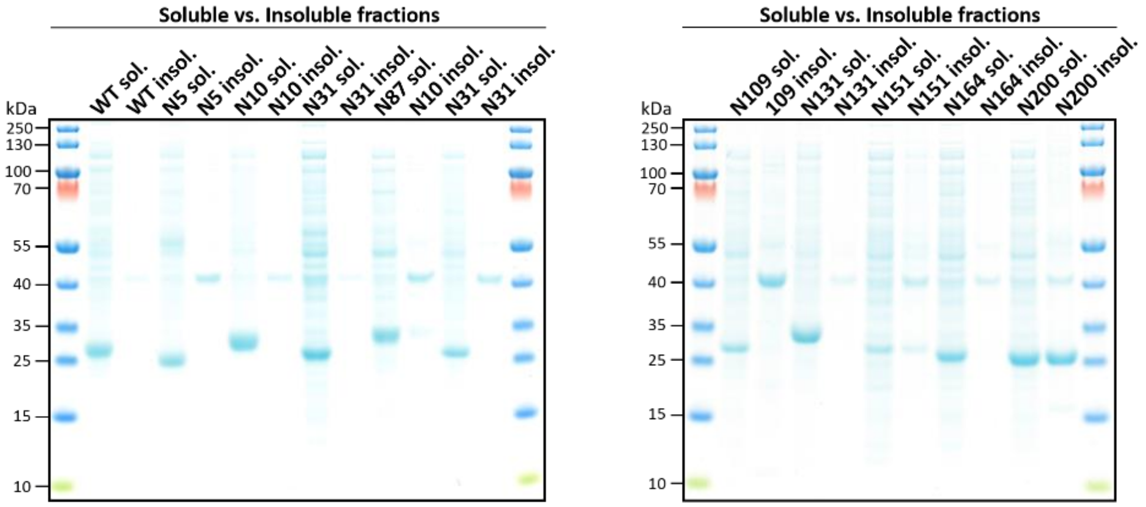

***Figure S8.*** *Comparison of soluble and insoluble fractions of the cell-free extract of overexpressed AMDase ancestors. SDS-PAGE samples were normalized to 10 µg total protein per pocket.*

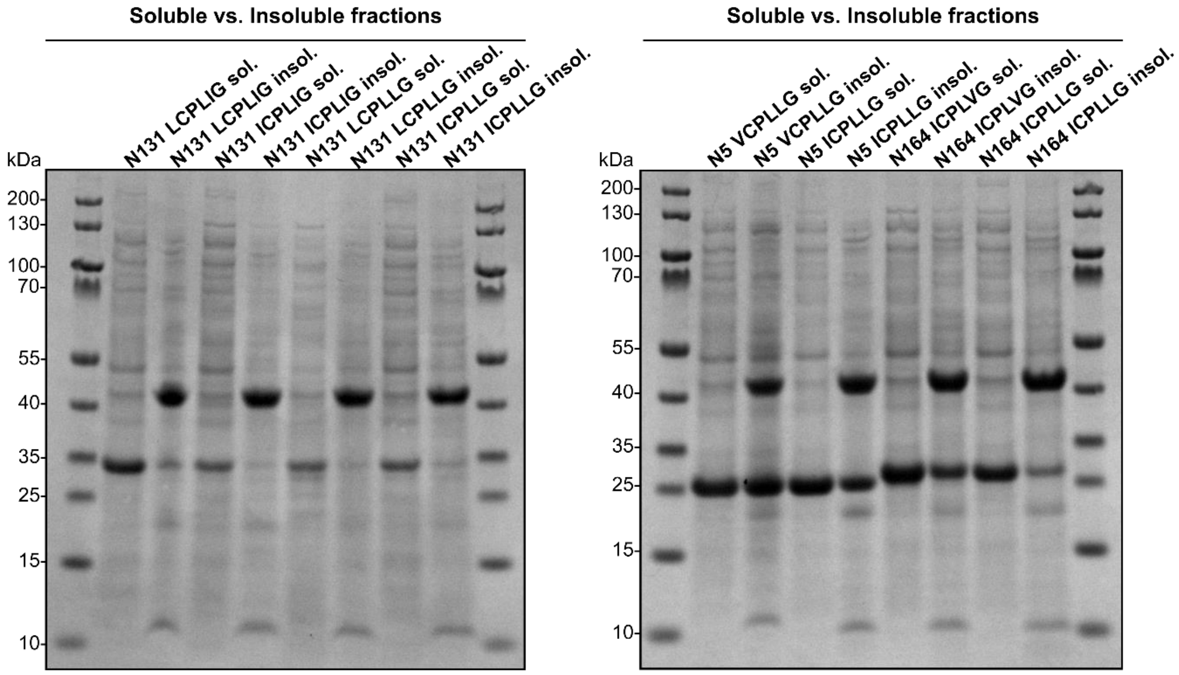

***Figure S9.*** *SDS-PAGE of the soluble and insoluble fractions, using the cell-free extract of the overexpressed ancestral (S)-selective AMDase variants. Per pocket, 3.75 µg of total protein was loaded.*

#### 7.5 Selectivity screening

For the screening of selectivity, freshly prepared CFE in Tris-HCl buffer (0 °C, 3.5–5 mg/mL total protein, 50 mM, pH 8) was mixed in equal parts with the respective substrate solution with further addition of CFE after 5 h (300 µL CFE: 300 µL substrate solution + 300 µL CFE, 16 h, 30 °C, 600 rpm). The substrate solution was prepared fresh in Tris-HCl buffer (50 mM, pH 8) with pH adjustment after complete dissolution using 3 M NaOH. For the (*R*)-selective ancestors, the final concentrations of the substrate solutions were: 2-alkyl-2-phenyl-malonic acids: 3.3 mM, 2-methyl-2-vinyl-malonic acid: 1.6 mM, 2-cyclohexene malonate: 1 mM. For the (*S*)-selective ancestors, the final concentrations of the substrate solutions were: 2-alkyl-2-phenyl-malonic acids: 2.5 mM, 2-alkyl-2-vinyl-malonic acid: 2 mM, 2-cyclohexene-1,1-dicarboxylic acid: 2 mM. After complete conversion, the reaction was acidified with HCl (4 M, 200 µL (*R*)-selective and 30 µL (*S*)-selective), and the product was extracted with 300 µL EtOAc and dried with anhydrous $MgSO_4$.

To ensure complete conversion of 2-cyclohexene-1,1-dicarboxylic acid, by the (*R*)-selective ancestors, and 2-ethyl-3-butenoic acid, by the (*S*)-selective AMDase ancestors, after 18 h of incubation, a 150 µL aliquot was analyzed by HPLC. For almost all variants, 300 µL of freshly prepared CFE was then added to the mixture and incubated for 3 h before proceeding with extraction. In the case of 2-cyclohexene malonic acid, when converted by *Bb*AMDase, in total 5 additions of fresh enzyme, which was freshly prepared or CFE that was stored on ice for up to 4 h, were necessary. 2-methyl-but-3-enoic acid was analyzed directly, and 2-alkyl-propionic acids were derivatized before chiral GC analysis. Therefore, the sample was incubated (30 min, 22 °C) with 100 µL MeOH and 25 µL TMS-$CH_2$-$N_2$. The reaction was terminated with 2.5 µL anhydrous acetic acid and was concentrated in a stream of nitrogen afterwards, the product was dissolved in 200 µL EtOAc for Chiral GC analysis. The samples were measured in triplicate of different split ratios (10, 20, 60, 70), and the conversion was reproduced once.

#### 7.6 Determination of initial AMDase activity on CFE

The initial activity of the AMDase ancestors was determined in a range of 10% substrate conversion for 5 minutes. Therefore, freshly prepared CFE (100 µL, 3.5–5 mg/mL total protein, 50 mM Tris-HCl, pH 8) was added to the incubated (30 °C, 600 rpm) substrate solution (final concentration: phenylmalonate and 2-methyl-2-phenyl malonate: 20 mM in 50 mM Tris-HCl buffer, pH 8). The conversion was determined by taking samples (100 µL at 0.15, 1.15, 2.15, 3.15, and 4.15 minutes), with the enzymatic reaction terminated in ACN: $H_2O$, 1:1 (900 µL). The samples were centrifuged (13,000 rpm, 4 °C, 20 min) and measured by HPLC. The mass-specific activity (U/mg) was calculated based on the quantified enzyme concentration by densitometric SDS-PAGE (BSA calibration: 83–667 µg/mL, evaluated with N131Analyzer software).

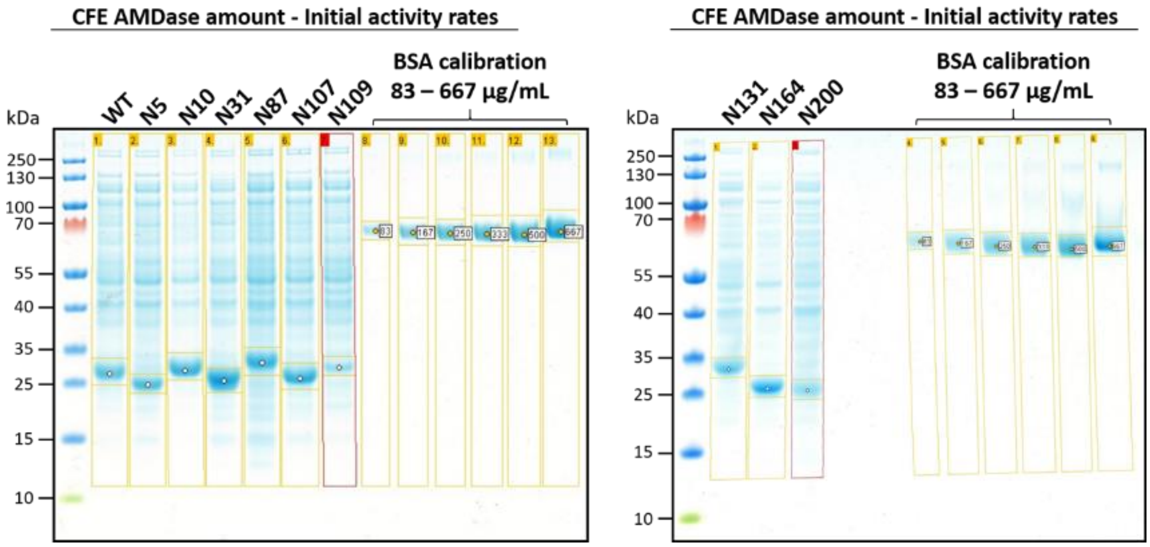

***Figure S10.*** *Densitometric quantification of AMDase in the cell-free extract (CFE) used to determine initial activity on phenylmalonate (PMA), and 2-methyl-2-phenyl malonate (Me-PMA). SDS-PAGE samples of CFE were normalized to 10 µg total protein and quantified to a BSA calibration (83–667 µg/mL) using GelAnalyzer.*

##### 7.7 Determination of kinetic parameters for AMDase ancestors N31 and N131, with purified enzyme.

The initial rates of AMDase ancestors N31 and N131 were determined with freshly prepared purified enzyme towards 2-methyl-2-phenyl malonate (**2a**) and 2-methyl-2-vinylmalonic acid (**4a**) (r=3). The rates were determined below 10% substrate conversion. The initial rates for (**2a**) were determined at 12 substrate concentrations: 0.111, 0.25, 0.5, 1, 2, 3.5, 5, 7, 15, 30, 50 and 70 mM. The initial rates for (**4a**) were determined at 16 substrate concentrations: 0.111, 0.25, 0.5, 0.8, 1, 1.5, 2, 3, 4, 5, 8, 10, 15, 20, 30 and 50 mM.

AMDase ancestor N31 (100 µL, 0,05 mg/mL for (**2a**) and 1 mg/mL for (**4a**), 50 mM Tris-HCl, pH 8) and ancestor N131 (100 µL, 0,03 mg/mL for (**2a**) and 0.3 mg/mL for (**4a**), 50 mM Tris-HCl, pH 8) were incubated for 30 sec and the enzyme solution was then added to the 2-3 min incubated (30 °C, 600 rpm) respective substrate solution (900 µL). The conversion was determined by taking samples (5 time points of 100 µL, every 1 min for (**2a**) and 5/10 min >0.5 mM/<0.5 mM for (**4a**), with the enzymatic reaction terminated in 4 M HCl (10 µL) in case for N31 and with 200 µL ACN/$H_2O$ (9/1) in case for N131. The different termination procedure was assessed to not influence the initial rate determination. In both cases the samples were centrifuged (13,000 rpm, 4 °C, 30 min) and the product concentration determined by HPLC. The samples were either analyzed directly or stored at -20°C for up to 2 days.

***Table S3.*** *Summarized data of the kinetic characterization of AMDase ancestors N31 and N131 towards 2-methyl-2-phenylmalonic acid (**2a**) and 2-vinyl-2-methyl malonic acid (**4a**). n.d. = not determined.*

| AMDase ancestor | Substrate | $k_{cat}$ [$s^{-1}$] | $K_M$ [mM] | $k_{cat}/K_M$ [$s^{-1}$ $mM^{-1}$] | $K_i$ [mM] |
|---|---|---|---|---|---|
| **N31** | **2a** | 10.0 ± 0.5 | 6.0 ± 0.4 | 1.7 | n.d. |
| | **4a** | 0.1 ± 0.0 | 9.9 ± 0.4 | 0.0 | n.d. |
| **N131** | **2a** | 33.9 ± 0.5 | 0.3 ± 0.0 | 103.9 | n.d. |
| | **4a** | 0.8 ± 0.1 | 1.0 ± 0.3 | 0.8 | 17.8 ± 6.2 |

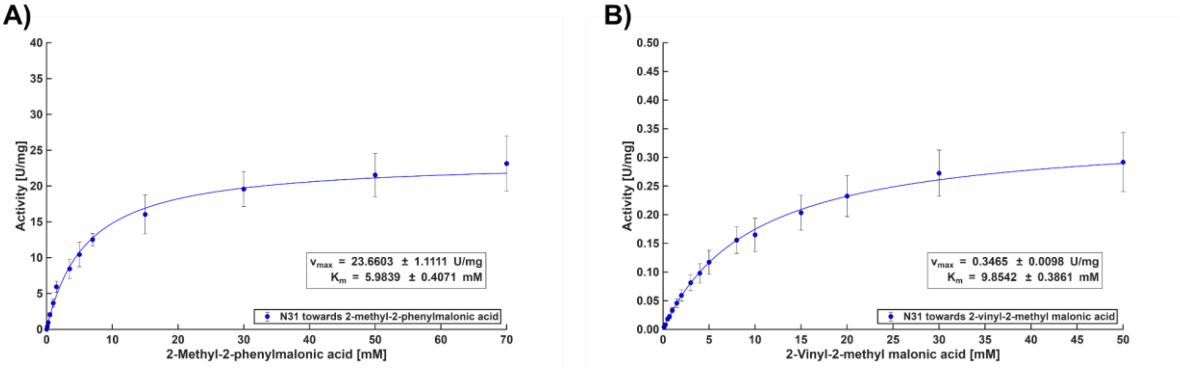

***Figure S11.*** *Michaelis-Menten kinetic characterization of AMDase N31. The data were fitted in MATLAB.* ***A)*** *Towards 2-methyl-2-phenylmalonic acid (**2a**).* ***B)*** *Towards 2-vinyl-2-methyl malonic acid (**4a**).*

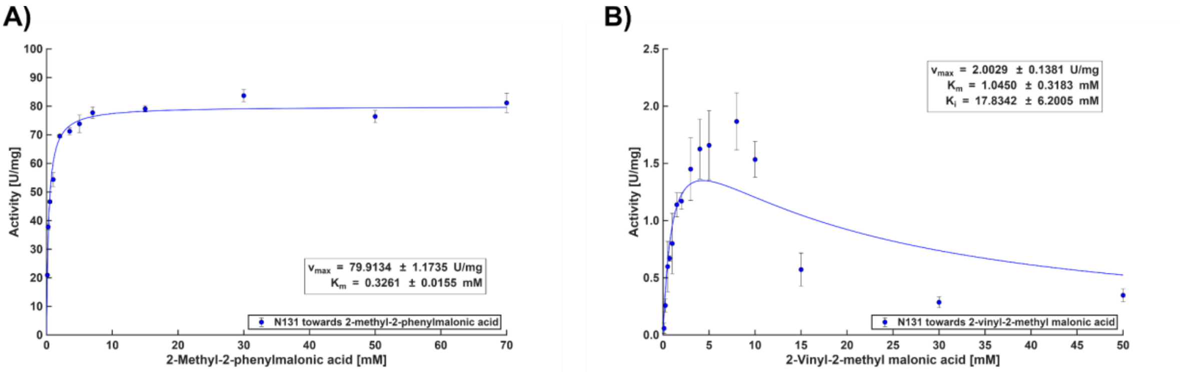

***Figure S12.*** *Michaelis-Menten kinetic characterization of AMDase N131. The data were fitted in MATLAB.* ***A)*** *Towards 2-methyl-2-phenylmalonic acid (**2a**).* ***B)*** *Towards 2-vinyl-2-methyl malonic acid (**4a**).*

#### 7.8 HPLC chromatograms

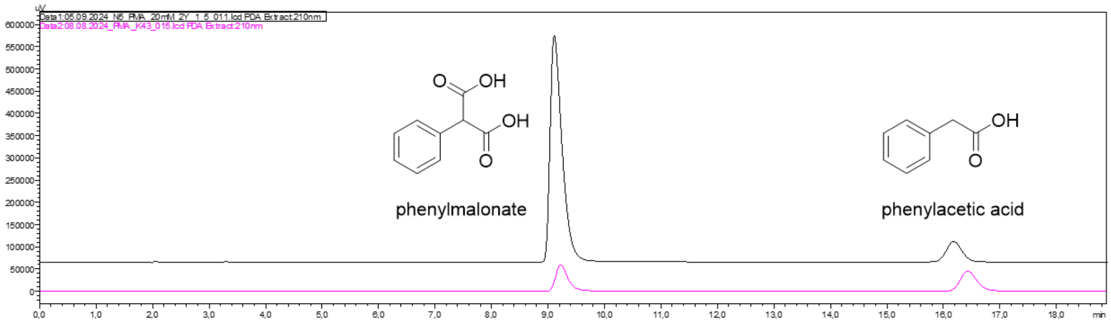

***Figure S13.*** *Exemplary HPLC chromatogram of the conversion of phenylmalonate with AMDase N5 (black) and the measurement of one calibration standard (pink) at 210 nm. Retention times: phenylmalonate = 9.1 min, phenylacetic acid = 16.2 min.*

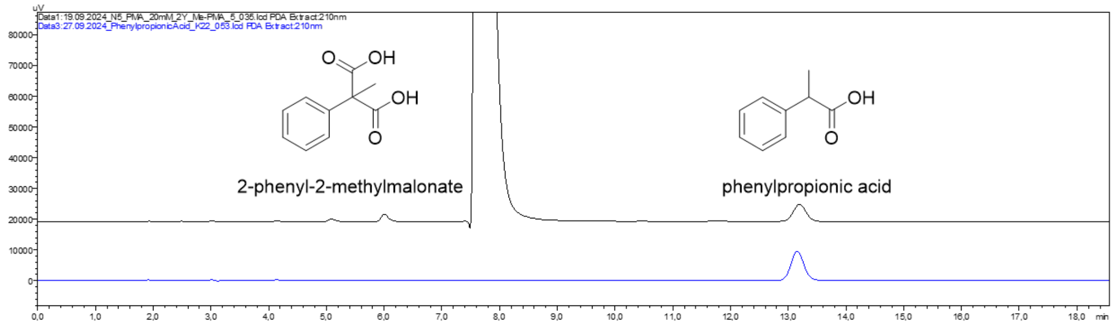

***Figure S14.*** *Exemplary HPLC chromatogram of the conversion of 2-methyl-2-phenylmalonic acid with AMDase N5 (black) and the measurement of one calibration standard (blue) at 210 nm. Retention times: 2-methyl-2-phenyl-malonic acid = 7.9 min, 2-phenyl propionic acid = 13.1 min.*

#### 7.9 Chiral GC chromatograms

In the following section, chromatograms from the better-performing ancestral variants will be displayed, compared to the performance of the corresponding reference extant enzyme. For the (*R*)-selective AMDases, the reference enzyme is *Bb*AMDase, whereas for the (*S*)-selective ancestral AMDase variants, *Bb*AMDase ICPLLG. For the substrates for which none of the ancestral variants had improved performance, an exemplary chromatogram will be shown.

##### 7.9.1 (*R*)-selective AMDases

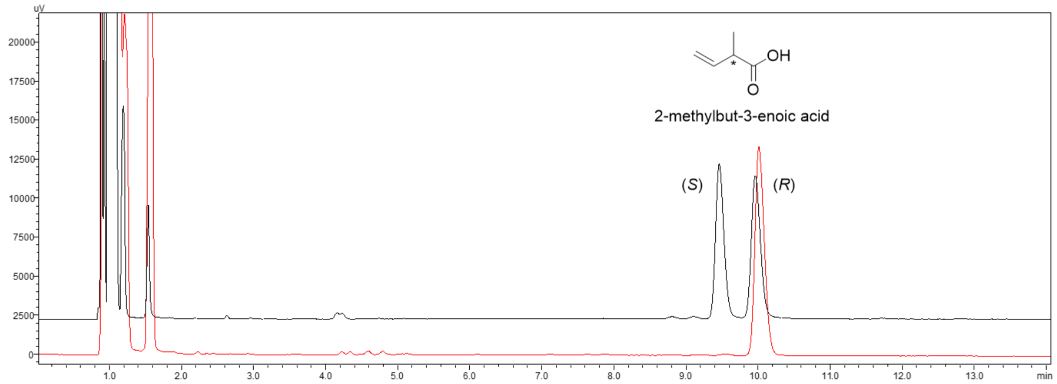

***Figure S15.*** *GC-FID chromatogram of 2-methylbut-3-enoic acid after the conversion of 2-vinyl-2-methyl malonate with AMDase N5 (red) superimposed with the racemic standard (black). Identification of the enantiomer was based on literature.*[10] *Retention times: (S)-2-methylbut-3-enoic acid = 9.3 min, (R)-2-methylbut-3-enoic acid = 9.8 min.*

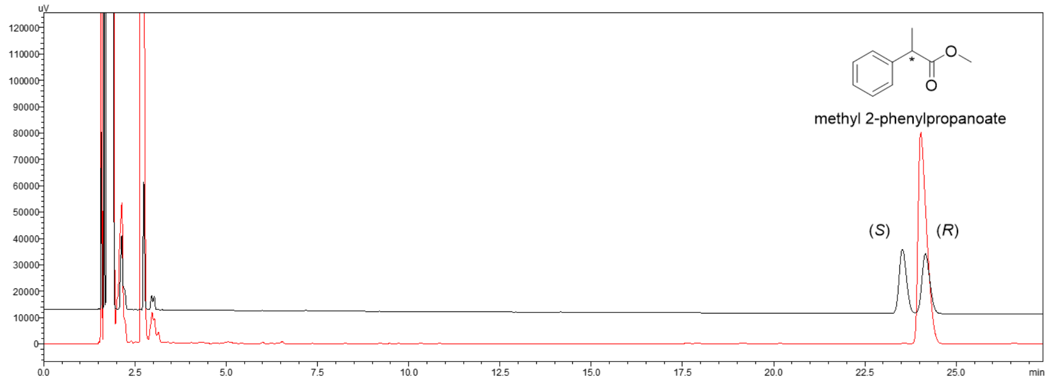

***Figure S16.*** *GC-FID chromatogram of methyl-2-phenylpropanoate after the conversion of 2-methyl-2-phenylmalonate with AMDase N5 (red) and subsequent derivatization with TMS-$CH_2$-$N_2$. The chromatogram is superimposed with the racemic standard (black). Identification of the enantiomer was based on literature.*[11] *Retention times: (S)-methyl-2-phenylpropanoate = 23.4 min, (R)-methyl-2-phenylpropanoate = 24.2 min.*

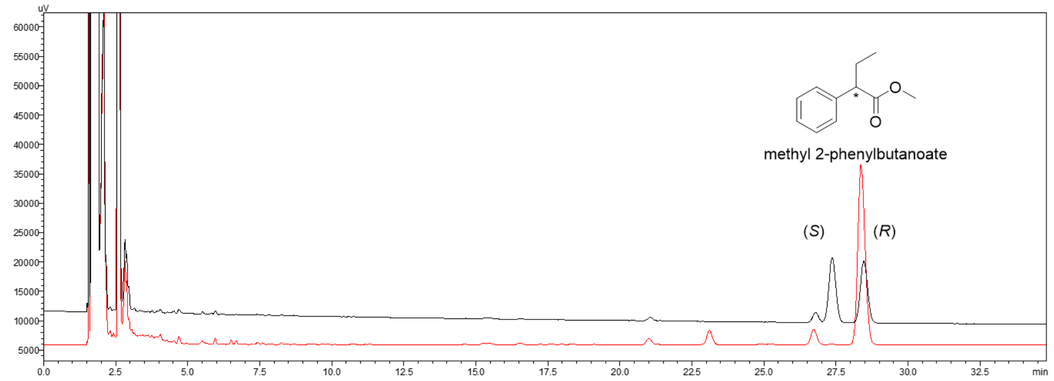

***Figure S17.*** *GC-FID chromatogram of methyl-2-phenylbutanoate after the conversion of 2-ethyl-2-phenylmalonate with AMDase N5 (red) and subsequent derivatization with TMS-$CH_2$-$N_2$. The chromatogram is superimposed with the racemic standard (black). Retention times: (S)-methyl-2-phenylbutanoate = 27.2 min, (R)-methyl-2-phenylbutanoate = 28.5 min.*

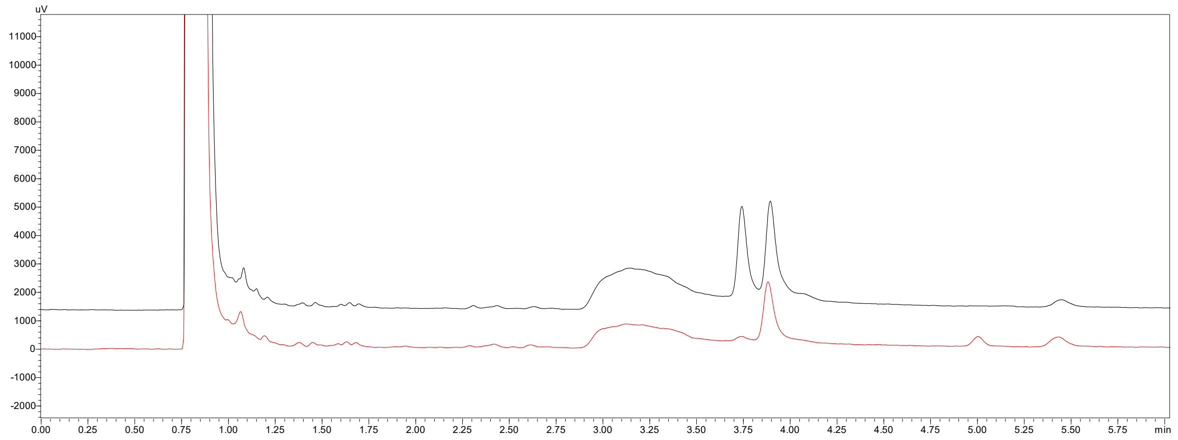

***Figure S18.*** *GC-FID chromatogram of cyclohex-2-ene-1-carboxylic acid after the conversion of 2-cyclohexene-1,1-dicarboxylate with AMDase N131 (red). The chromatogram is superimposed with the racemic standard (black). Retention times: 1st eluting peak = 3.7 min, 2nd eluting peak = 3.9 min.*

##### 7.9.2 (*S*)-selective AMDases

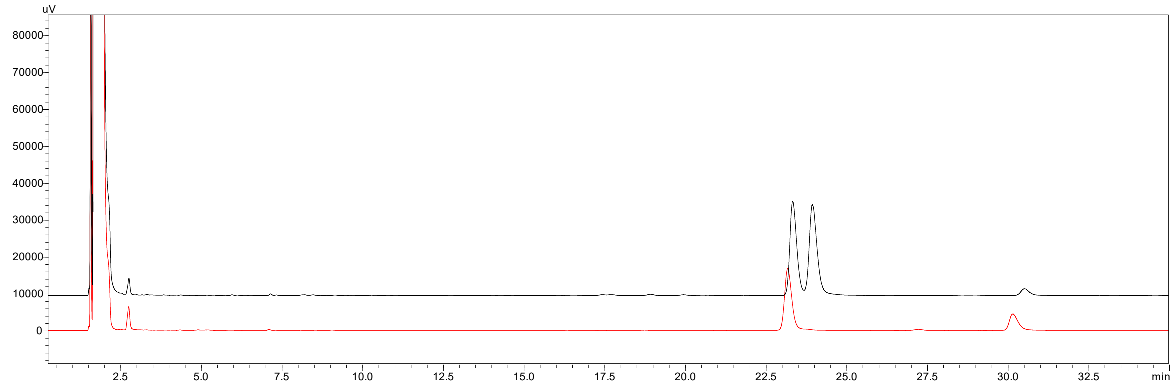

***Figure S19.*** *GC-FID chromatogram of methyl-2-phenylpropanoate after the conversion of 2-methyl-2-phenylmalonate with AMDase N5 VCPLLG (red) and subsequent derivatization with TMS-CH$_2$-N$_2$. The chromatogram is superimposed with the racemic standard (black). Retention times: (S)-methyl-2-phenylpropanoate = 23.4 min, (R)-methyl-2-phenylpropanoate = 24 min.*

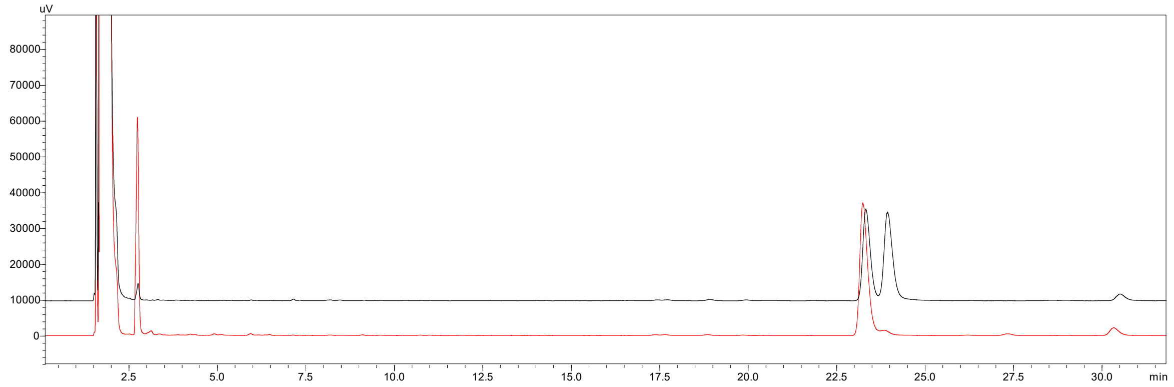

***Figure S20.*** *GC-FID chromatogram of methyl-2-phenylpropanoate after the conversion of 2-methyl-2-phenylmalonate with AMDase N131 LCPLLG (red) and subsequent derivatization with TMS-CH$_2$-N$_2$. The chromatogram is superimposed with the racemic standard (black). Retention times: (S)-methyl-2-phenylpropanoate = 23.4 min, (R)-methyl-2-phenylpropanoate = 24 min.*

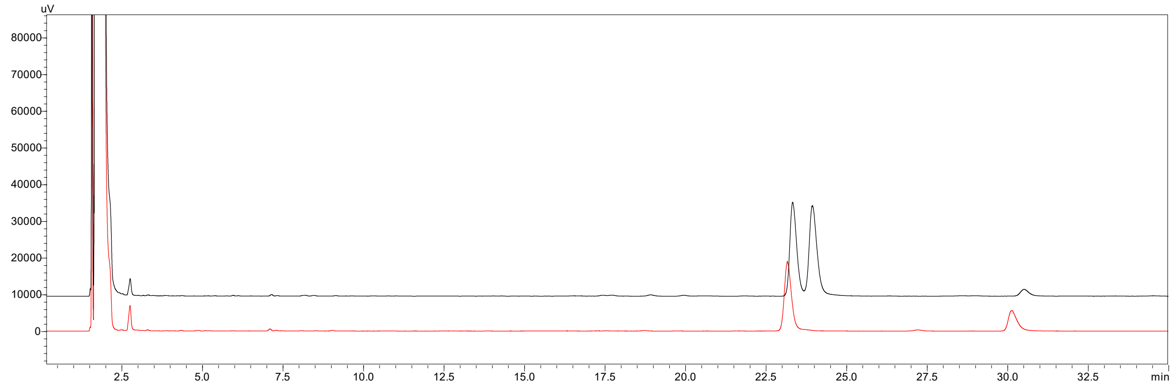

***Figure S21.*** *GC-FID chromatogram of methyl-2-phenylpropanoate after the conversion of 2-methyl-2-phenylmalonate with AMDase N131 ICPLLG (red) and subsequent derivatization with TMS-CH$_2$-N$_2$. The chromatogram is superimposed with the racemic standard (black). Retention times: (S)-methyl-2-phenylpropanoate = 23.4 min, (R)-methyl-2-phenylpropanoate = 24 min.*

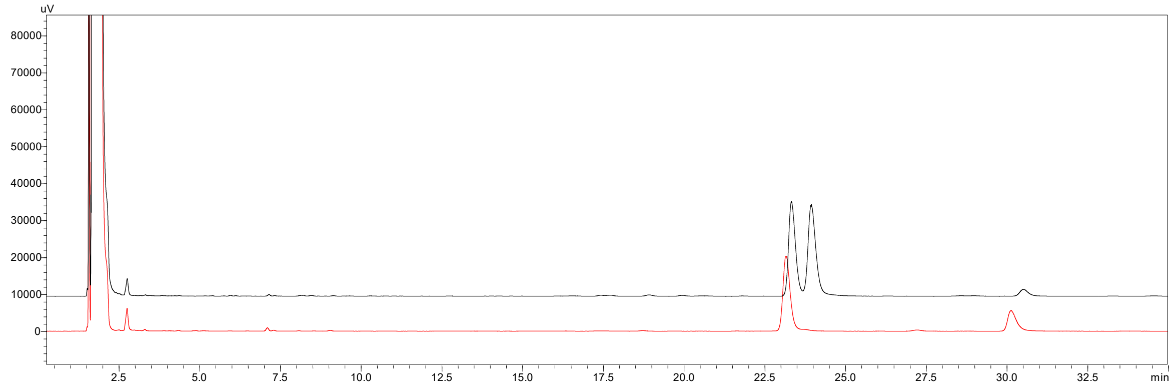

***Figure S22.*** *GC-FID chromatogram of methyl-2-phenylpropanoate after the conversion of 2-methyl-2-phenylmalonate with AMDase N164 ICPLLG (red) and subsequent derivatization with TMS-CH$_2$-N$_2$. The chromatogram is superimposed with the racemic standard (black). Retention times: (S)-methyl-2-phenylpropanoate = 23.4 min, (R)-methyl-2-phenylpropanoate = 24 min.*

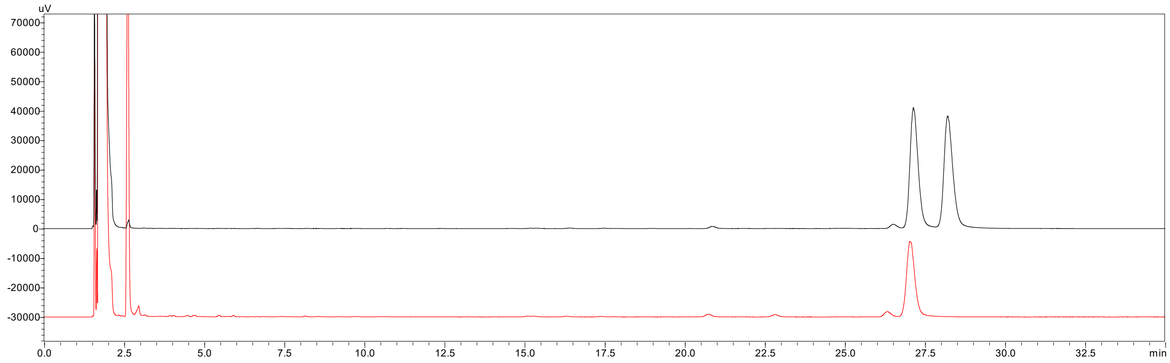

***Figure S23.*** *GC-FID chromatogram of methyl-2-phenylbutanoate after the conversion of 2-ethyl-2-phenylmalonate with AMDase N164 ICPLVG (red) and subsequent derivatization with TMS-CH$_2$-N$_2$.*

*The chromatogram is superimposed with the racemic standard (black). Retention times: (S)-methyl-2-phenylbutanoate = 27.2 min, (R)-methyl-2-phenylbutanoate = 28.5 min.*

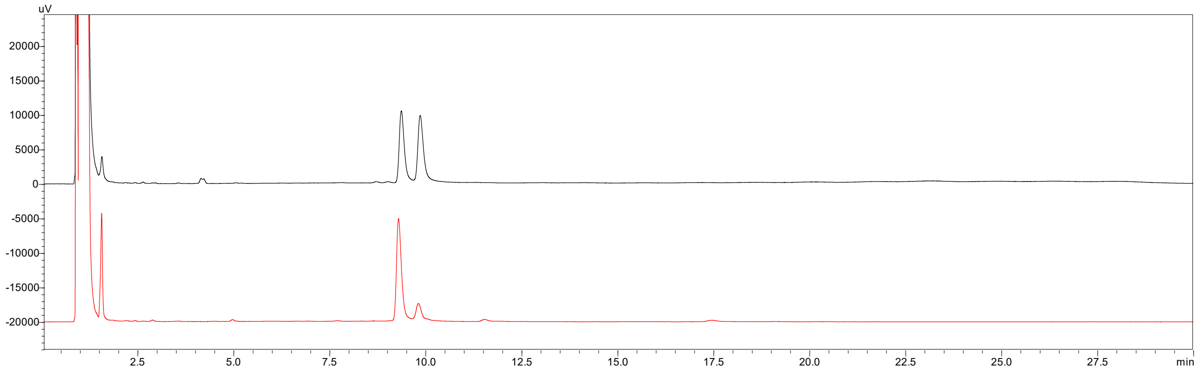

***Figure S24.*** *GC-FID chromatogram 2-methylbut-3-enoic acid after the conversion of 2-vinyl-2-methyl malonate with AMDase N5 VCPLLG (red). The chromatogram is superimposed with the racemic standard (black). Retention times: (S)-2-methylbut-3-enoic acid = 9.3 min, (R)-2-methylbut-3-enoic acid = 9.9 min.*

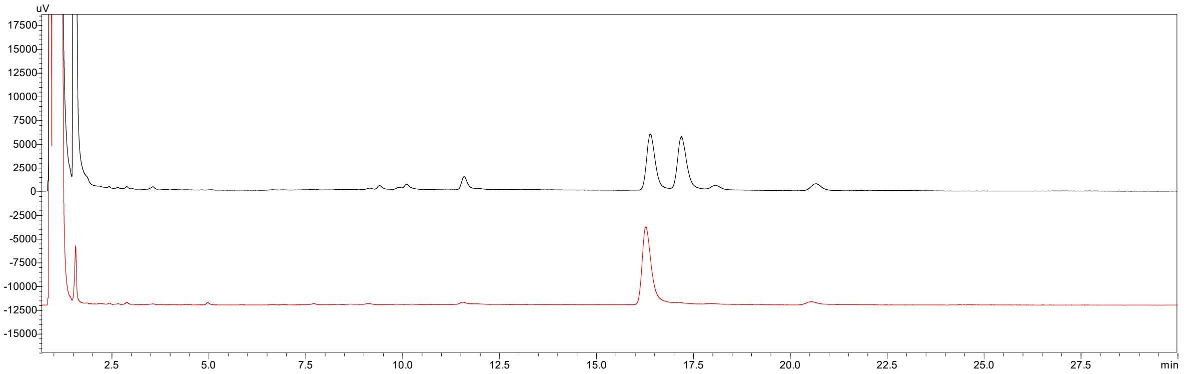

***Figure S25.*** *GC-FID chromatogram 2-ethylbut-3-enoic acid after the conversion of 2-ethyl-2-vinyl malonate with AMDase N5 VCPLLG (red). The chromatogram is superimposed with the racemic standard (black). Retention times: (S)-2-ethylbut-3-enoic acid = 16.4 min, (R)-2-ethylbut-3-enoic acid = 17.2 min.*

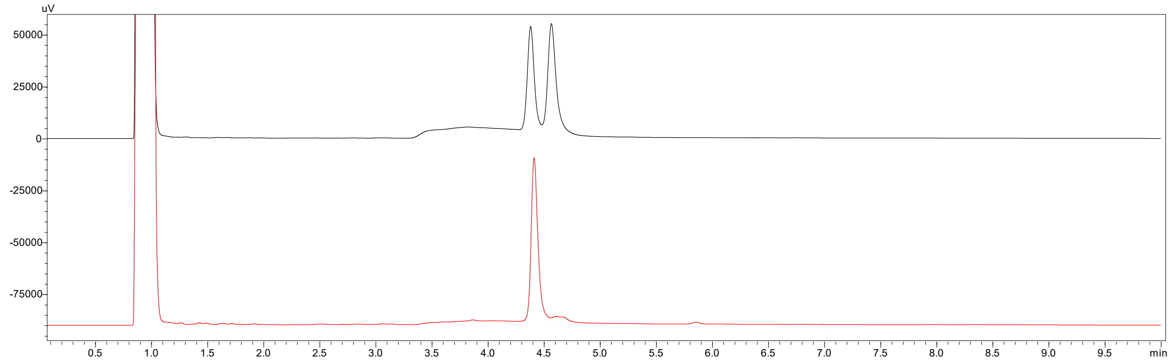

***Figure S26.*** *GC-FID chromatogram of cyclohex-2-ene-1-carboxylic acid after the conversion of 2-cyclohexene-1,1-dicarboxylate with AMDase N131 LCPLLG (red). The chromatogram is superimposed with the racemic standard (black). Retention times: first eluting peak = 4.4 min, second eluting peak = 4.6 min.*

#### 7.10 Enzyme purification

For the first screening of half-life time and unfolding temperature, protein purification was performed using Ni-SEPHAROSE gravity columns (2 mL resin, Cytiva 17-5268-01) at a constant temperature of 4 °C. The composition of purification buffers is listed below. First, the CFE was resuspended in binding buffer (12 mL) and incubated for 20 minutes with the equilibrated resin under gentle shaking and subsequently washed with binding buffer (4 x 12 mL). The enzyme was eluted into a reaction tube by incubation of the resin in elution buffer (1 mL, 5 min). Desalting was done using PD-10 desalting columns (Cytiva 17085101), being equilibrated with the respective buffer (half-life time determination: 50 mM TRIS-HCl, pH 8; unfolding temperature: 50 mM KPi, pH 6). The protein was eluted in four separate 1 mL fractions, with subsequent BCA and SDS-page analysis. Further comparison of the most stable ancestors, N5, N131, and N164, to *Bb*AMDase was performed on purified enzyme using the ÄKTA pure system at a constant 4 °C with a 5 mL HisTrap FF Crude column (Cytiva, 17528601). The same buffer composition and desalting methodology were used as in the initial screening study. The eluted enzyme was received in a 1 mL fraction and used as described above.

Automated purification method using ÄKTA pure and Cytiva F9-T fraction collector: **Column volume (CV):** 5 mL, **Equilibration:** 2 mL/min, 5 CV, binding buffer; **Sample Application:** 1 mL/min, 50 mL Superloop filled with 20 mL CFE; **Column wash:** 2 mL/min, 15 CV, **Elution:** 0.5 mL/min up flow, 1 mL fraction collection, 4 CV, elution buffer, **Equilibration:** 2.5 mL/min, 10 CV, elution buffer → binding buffer 5 CV.

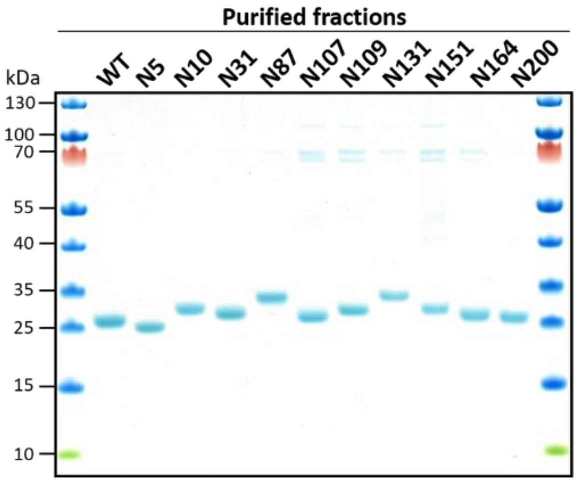

***Figure S27.*** *SDS-PAGE analysis of AMDase wildtype compared to the ancestors purified by Ni-SEPHAROSE gravity columns. Samples were normalized to 2 µg protein per pocket.*

#### 7.11 Determination of Unfolding Temperature by Differential Scanning Fluorimetry (DSF)

Unfolding temperatures were determined using Differential Scanning Fluorimetry (DSF) *via* the thermofluor method. The assay was done in triplicate with different protein concentrations (5, 10, and 15 µM, 45 µL, 50 mM KPi, pH 6) with the addition of SYPRP Orange (5 µL, 200-fold dilution in 50 mM KPi, pH 6, Sigma-Aldrich). Buffer was used as a control with the subtraction of the sample signals. Fluorescence was determined in intervals of 0.5 minutes during the temperature ramp from 25 to 95 °C with 0.5 °C/min increments. Detection of the signal was observed by RT-PCR(Rotor-Gene Q cycler, Qiagen) with FAM (green), ROX (orange), and TET (yellow) filters.

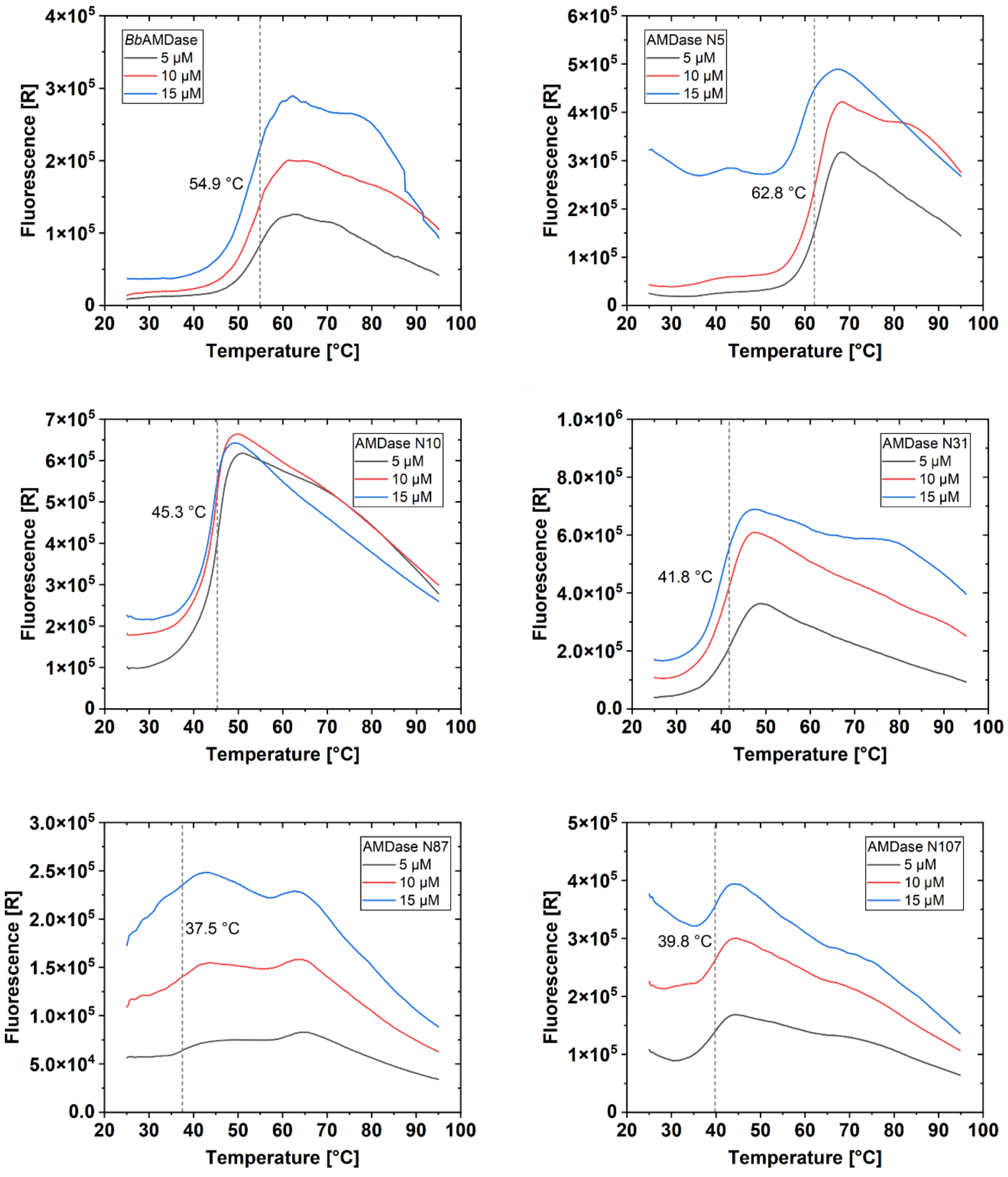

***Figure S28.*** *Unfolding curves of BbAMDase and ancestors N5 to N107 determined by Differential Scanning Fluorimetry (DSF).*

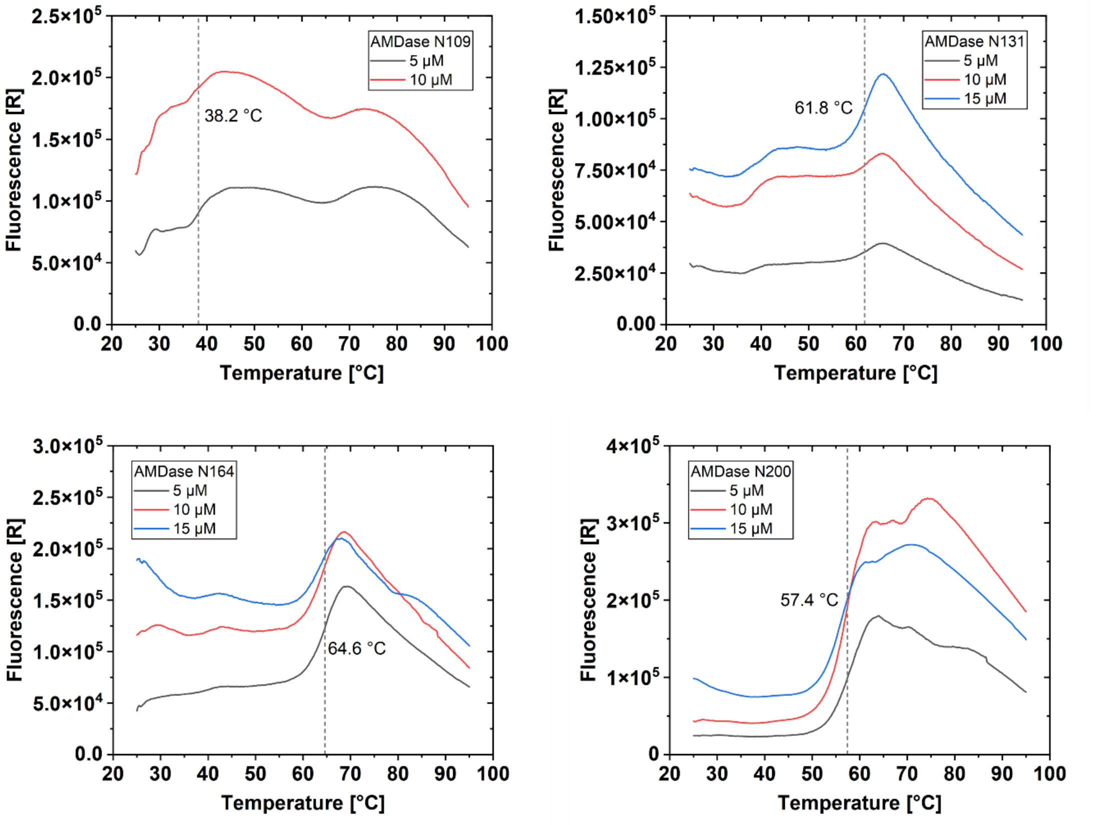

***Figure S29.*** *Unfolding curves of ancestors N109 to N200 determined by Differential Scanning Fluorimetry (DSF).*

#### 7.12 Determination of Unfolding Temperature by Circular Dichroism (CD) measurements

CD measurements to determine the unfolding temperatures of *Bb*AMDase, N131, and N164 were performed using a Jasco J-1500 with purified enzyme (0.2–0.4 mg/mL) in KPi (50 mM, pH 8). Briefly, 300 µL of the sample was loaded into a 1 mm quartz cuvette, and a small layer of paraffin oil was added on top. The wavelength for the temperature-interval scan was set as the average of local minima of five full wavelength scans at a constant temperature of 20 °C. The temperature-interval scan was then performed up to 95 °C.

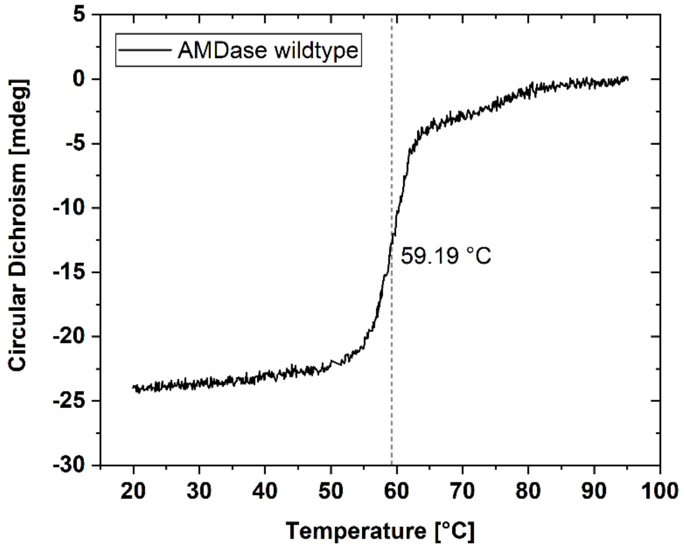

***Figure S30.*** *Circular dichroism (CD) spectrum of BbAMDase. The unfolding temperature of 59.19 °C was determined by a temperature-interval scan at 221 nm.*

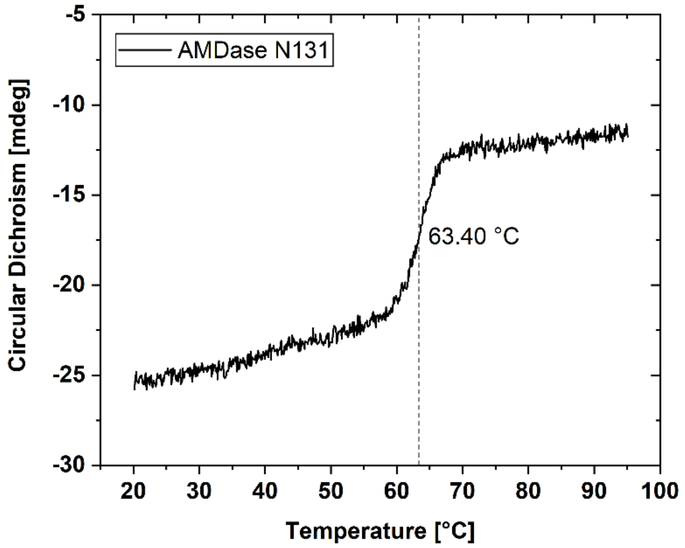

***Figure S31.*** *Circular dichroism (CD) spectrum of AMDase N131. The unfolding temperature of 63.40 °C was determined by a temperature-interval scan at 221 nm.*

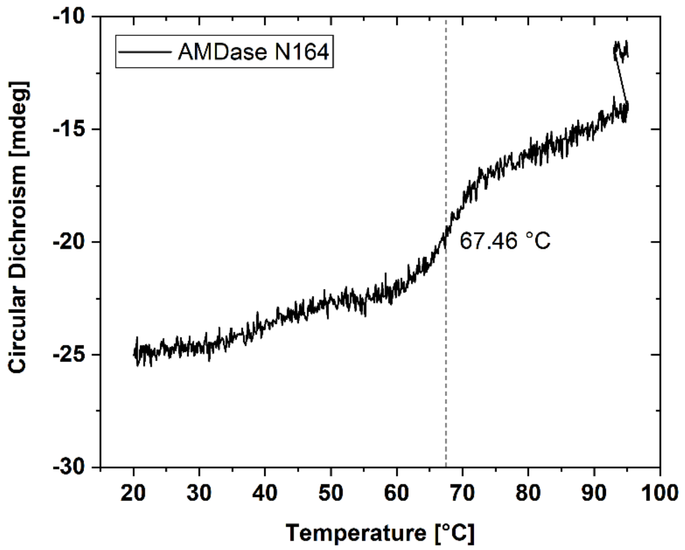

***Figure S32.*** *Circular dichroism (CD) spectrum of AMDase N164. The unfolding temperature of 67.46 °C was determined by a temperature-interval scan at 221 nm.*

#### 7.13 Determination of Half-life Time of AMDase

Determination of half-life time of *Bb*AMDase and the AMDase ancestors was performed using purified enzyme with determination of initial activity rates of the incubated aliquots. Therefore, the purified enzyme (2–5 mg/mL) was diluted ten-fold (Tris-HCl 50 mM, pH 8, 4 °C) and the protein concentration determined by BCA-assay. The solution was split into nine aliquots of 110 µL in PCR tubes and incubated using a PCR cycler (30 °C constant, 110 °C lid temperature). For each measurement point, one aliquot was taken out and 100 µL diluted again ten-fold (Tris-HCl 50 mM, pH 8, 4 °C). The initial activity rate was determined by splitting the received solution into triplicate with the assay described above in **section 7.6**.

The half-life time determination of the most stable ancestors N5 and N131 was further repeated at 45 °C in comparison to *Bb*AMDase. For this, the described assay conditions were not changed except for the incubation temperature.

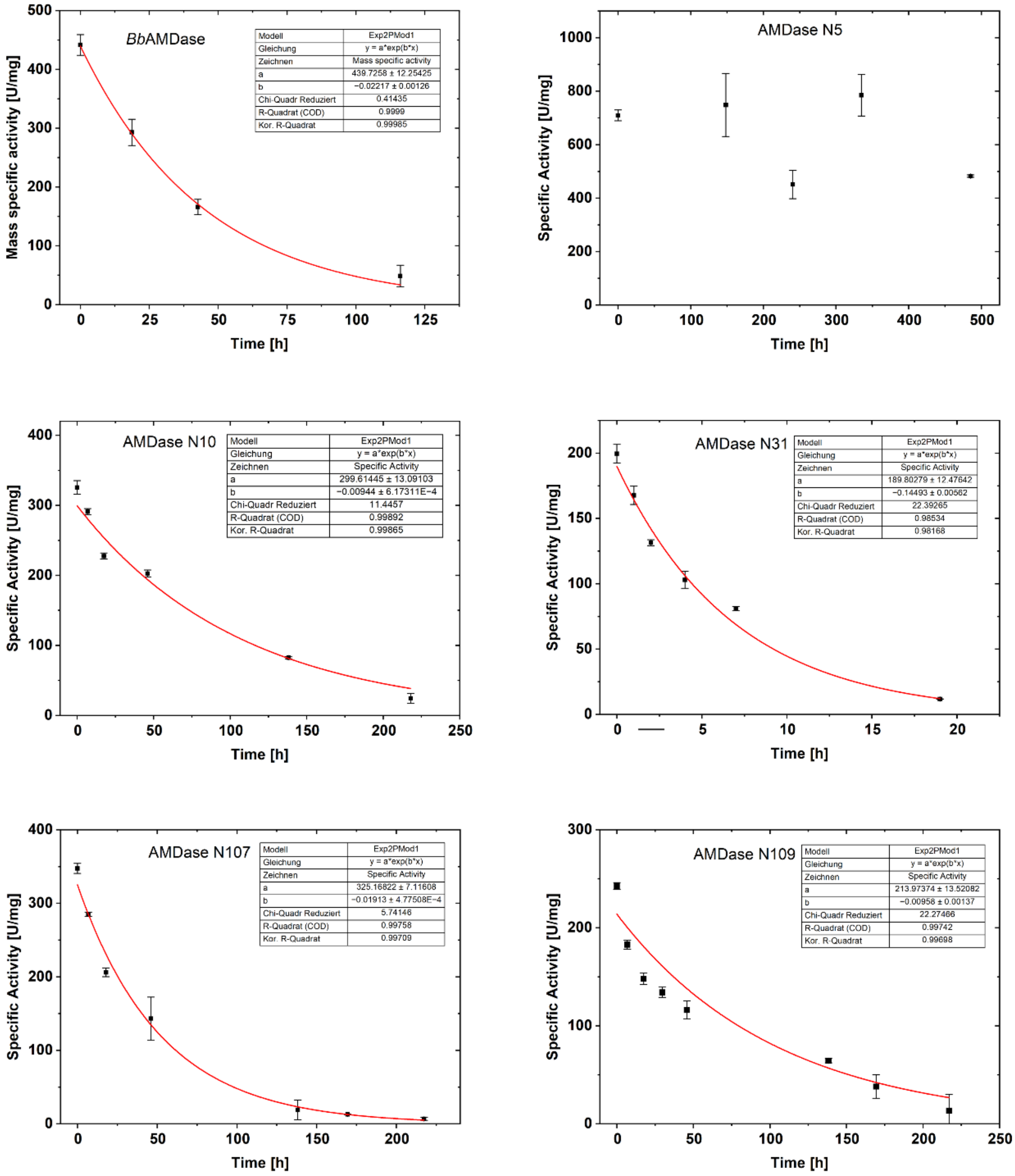

***Figure S33.*** *Time-course of initial mass-specific activity of purified BbAMDase and AMDase ancestors N5, N10, N31, N107, and N109.*

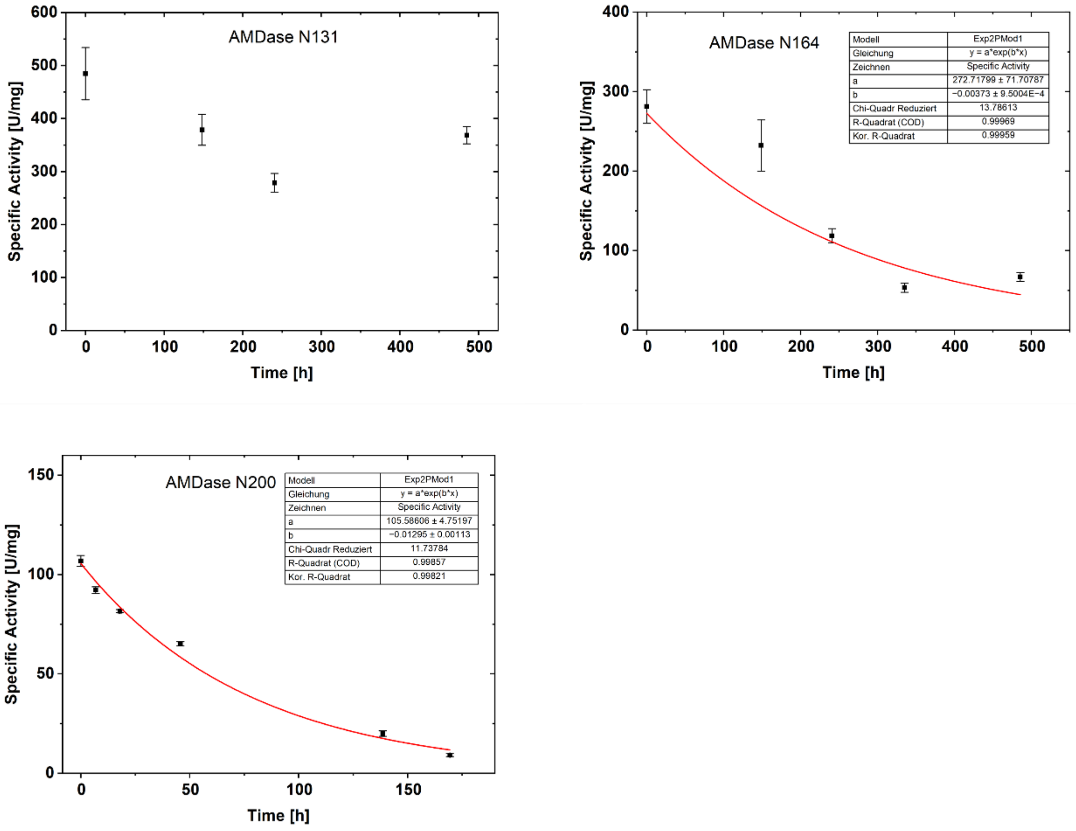

***Figure S34.*** *Time-course of initial mass-specific activity of purified AMDase ancestors N131, N164, and N200.*

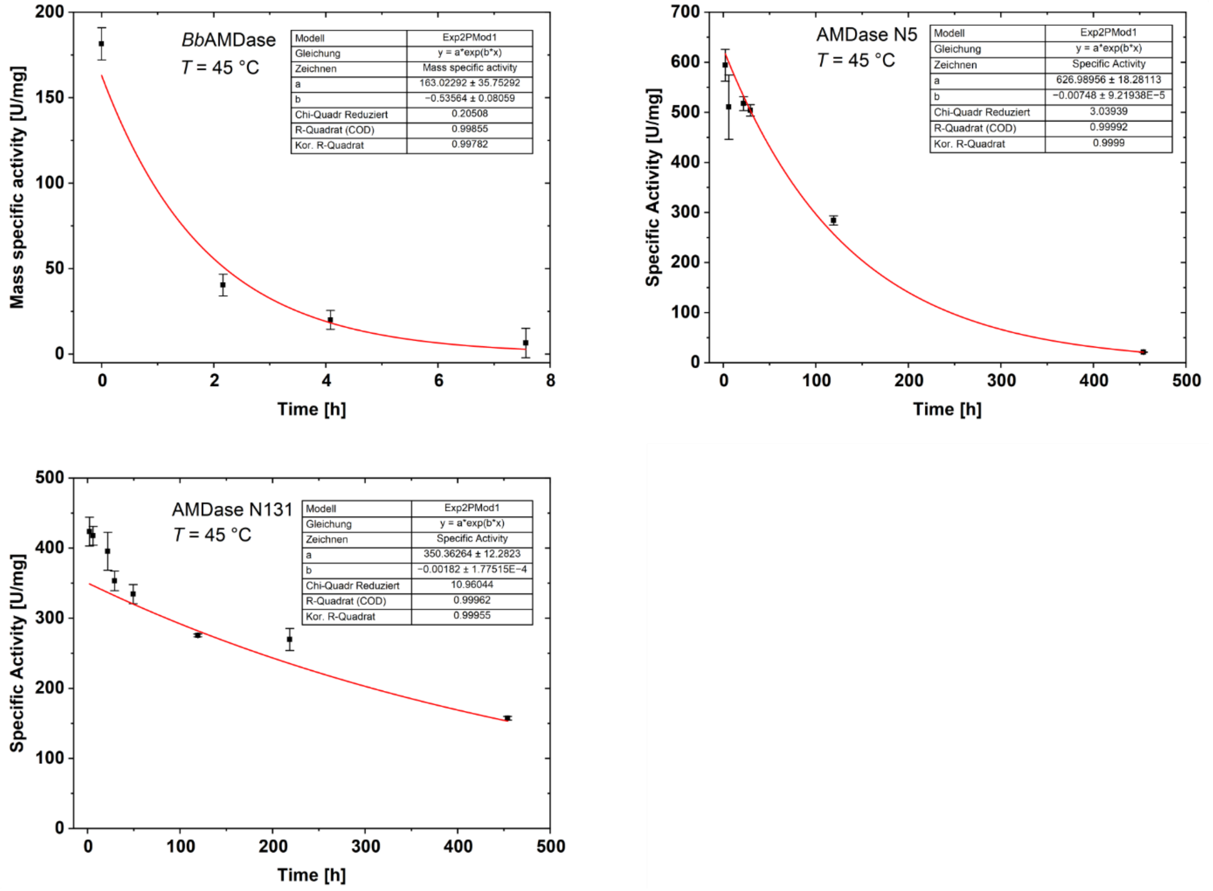

***Figure S35.*** *Determination of half-life time by fitting the time-course of initial mass-specific activity of purified BbAMDase and AMDase ancestors N5, N131 and N164 incubated at 45 °C.*

The observed total turnover numbers (*TTN*) were calculated in reference to Bommarius with equations S1–S3.[12] The TTN for the half-life determination at 30 °C are listed in ***Table S4***, and for the half-life determination at 45 °C are listed in ***Table S5***.

$$k_{cat,obs}\ (s^{-1}) = \frac{specific\ activity\ (\mathrm{U\ mg^{-1}}) \cdot enzyme\ molecular\ mass\ (\mathrm{g\ mol^{-1}})}{60\ 000} \qquad \text{(S1)}$$

$$k_{d,obs}\ (\mathrm{s^{-1}}) = \frac{\ln(2)}{half\ life} \qquad \text{(S2)}$$

$$TTN = \frac{k_{cat,obs}\ (\mathrm{s^{-1}})}{k_{d,obs}\ (\mathrm{s^{-1}})} \qquad \text{(S3)}$$

***Table S4.*** *Observed total turnover numbers for the half-life determination at 30 °C. The half-life of N5 and N131 was extended under the experimental conditions and could therefore not be determined. n.d.: not determined.*

| Enzyme | TTN |
|---|---|
| ***Bb*AMDase** | 18124462 |
| **N5** | n.d. |
| **N10** | 49511822 |
| **N31** | 2042972 |
| **N87** | n.d. |
| **N107** | 26516738 |
| **N109** | 34842714 |
| **N131** | n.d. |
| **N151** | n.d. |
| **N164** | 87551131 |
| **N200** | 12718517 |

***Table S5.*** *Observed total turnover numbers for the half-life determination at 45 °C.*

| Enzyme | TTN |
|---|---|
| ***Bb*AMDase** | 474780 |
| **N5** | 130760535 |
| **N131** | 300308571 |

#### 7.14 Preparative Scale Application

The preparative scale application of *Bb*AMDase and the best-performing ancestor N131 was conducted in a SpinChem® vessel V3 equipped with a 45-degree pitch blade turbine impeller (70 rpm). For the conversion of 2-methyl-2-phenylmalonate (1 g, 20 mM in 50 mM Tris-HCl, pH 8), the substrate was incubated to reach the reaction temperature (30 °C) and decarboxylated by the addition of CFE (12 mL; *Bb*AMDase: 4.1 mg/mL, N131: 2.8 mg/mL total protein determined by BCA). The conversion was determined using HPLC-DAD as described above. The product was isolated by acidifying the fully converted reaction mixture to pH 2 with 1 M HCl, followed by EtOAc extraction (3x 100 mL). The combined organic phases were dried with brine (50 mL) and $MgSO_4$, with subsequent removal of solvent under reduced pressure. The product was purified by flash-column chromatography (EtOAc:Cyclohexane, 1:10 → 1:5) to receive (*R*)-2-phenylpropionic acid as a colorless oil (yields: *Bb*AMDase = 96.6%, N131 = 95.8%). The *ee* was determined using chiral GC as described in **section 3.1** (*Bb*AMDase: 99.5% *ee*, N131: 99.4% *ee*).

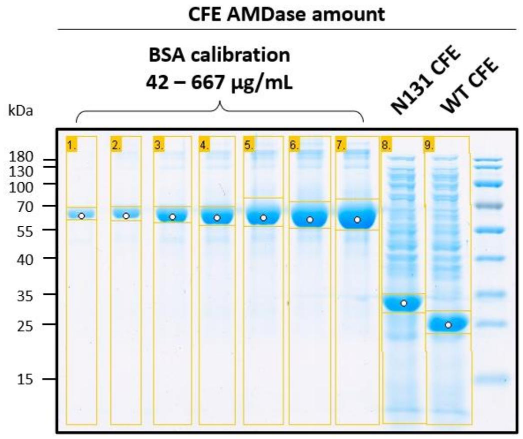

***Figure S36.*** *Determination of the amount of enzyme in the CFE of BbAMDase (WT) and ancestor N131 by densitometric SDS-PAGE analysis using GelAnalyzer 19.1 software. Enzyme overexpression yield: BbAMDase (WT) = 44.9%, N131 = 34.4%.*

**(*R*)-2-phenylpropionic acid**

**$^1$H-NMR** (400 MHz, $CDCl_3$): δ(ppm) = 7.40 – 7.33 (m, 5H), 7.35 – 7.29 (m, 1H), 7.31 – 7.25 (m, 1H), 3.77 (q, *J* = 7.2 Hz, 1H), 2.12 (s, 1H), 1.55 (d, *J* = 7.2 Hz, 3H)

**$^{13}$C-NMR** (101 MHz, $CDCl_3$): δ(ppm) = 180.65, 139.76, 128.70, 127.62, 127.42, 45.35, 18.12.

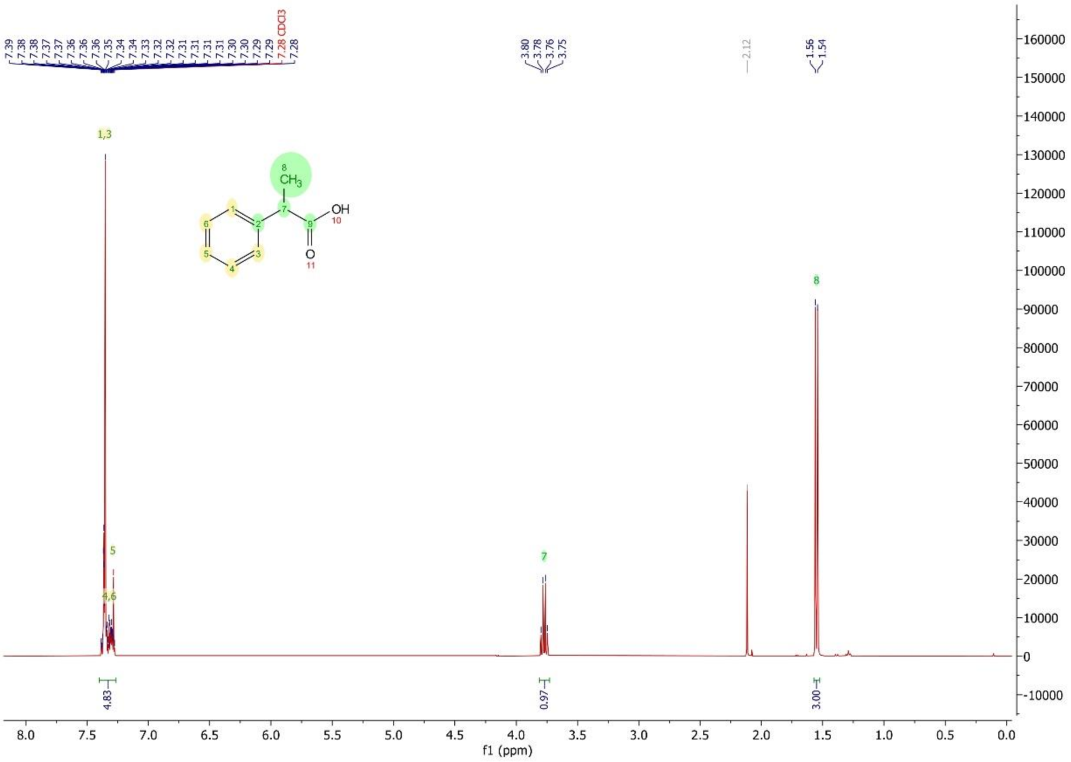

***Figure S37.*** *Measured $^1$H-NMR Spectrum of (R)-2-phenylpropionic acid in $CDCl_3$.*

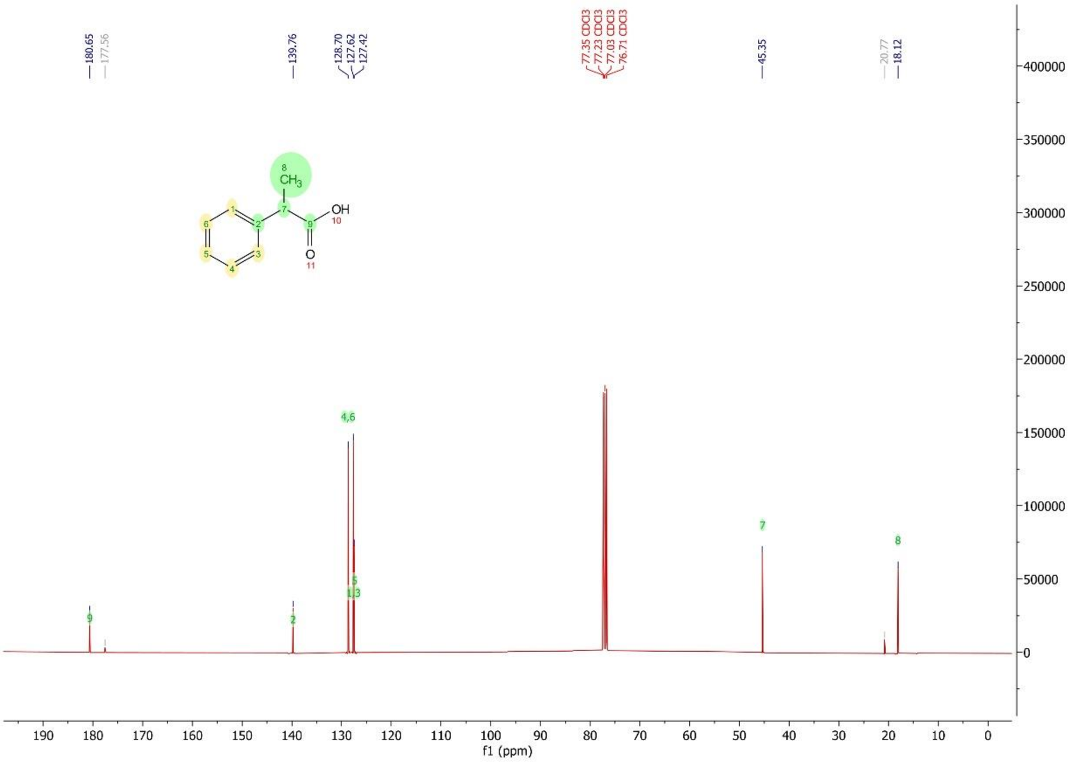

***Figure S38.*** *Measured* $^{13}$*C-NMR Spectrum of (R)-2-phenylpropionic acid in* $CDCl_3$.

#### 7.15 $^{13}$C isotope-labeled probe study

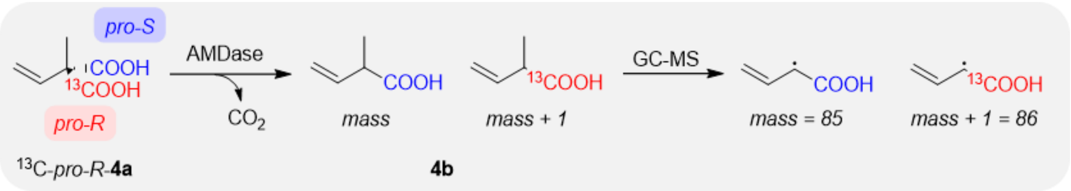

***Scheme S1.*** *Decarboxylation of* $^{13}$*C isotope-labeled* ***4a*** *(*$^{13}$*C-pro-R-**4a**) by AMDase. Cleavage of the isotope-labeled pro-R carboxylate group by AMDase variants yields* ***4b*** *(mass), whereas cleavage of the non-labeled pro-S carboxylate group yields* ***4b*** *(mass + 1). Due to the significantly higher intensities at m/z 85, we analyzed the signals at m/z 85/86.*[13]

The synthetic route towards $^{13}$C-*pro-R*-**4a**, experimental procedures, and analytical methods have been described in van der Pol *et al.*[13]

Biotransformation

AMDase N31 and N131 cell-free extract (500 μL) in Tris HCl, 50 mM, pH 8, (75 mg/mL wet cell mass) and (*R*)-2-methyl-2-vinylmalonic-1-$^{13}$C acid (500 μL, 10 mM) in Tris HCl (50 mM, pH 8) were combined in a 2 mL glass vial. The reactions were incubated at 30 °C, 600 rpm. After 2 h, AMDase CFE (250 μL) was added to the reaction mixtures, and after 4 h again (250 µL) to the reaction with N31 to ensure a completed reaction. Hence, the final substrate concentrations were 4 mM and 3.3 mM, respectively. After 4.5 h (N131) or 17 h (N31), the reactions went to completion according to TLC (EtOAc: cyclohexane:acetic acid, 3:3.5:0.5). After quenching the reaction by addition of HCl solution (4 M, 150 µL), the product was extracted with MTBE (1:3 (*v*/*v*), organic: aqueous) and centrifuged at 12 000 rpm, 10 min. The organic layer was dried over $MgSO_4$ and centrifuged at 12 000 rpm, 3 min. The samples were used for chiral GC-MS and chiral GC without any further modifications.

<u>Chiral GC-MS chromatograms</u>

**Decarboxylation of 4a by AMDase N31**

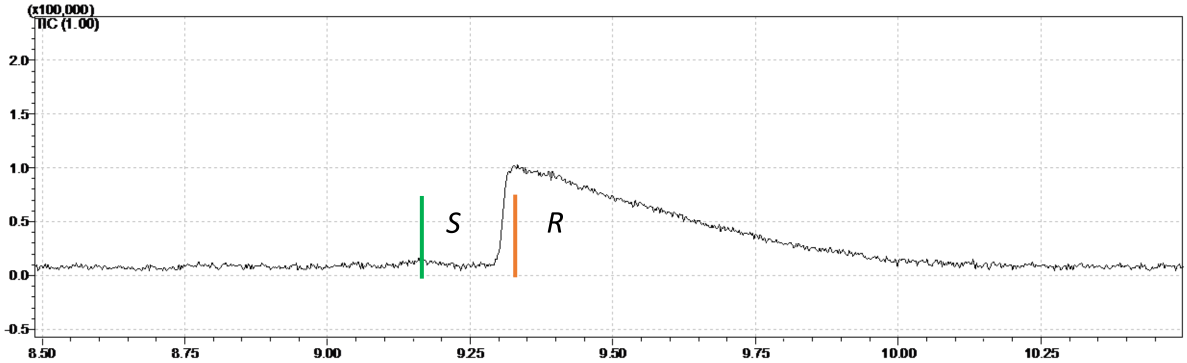

***Figure S39.*** *Chiral GC-MS chromatogram of 2-methyl-3-butenoic acid (**4b**), the product of (R)-2-methyl-2-vinylmalonic-1-$^{13}$C acid ($^{13}$C-proR-**4a**) converted by AMD N31. The green and orange bars indicate the retention times at which the mass spectra of (S)-**4b** and (R)-**4b** have been analyzed, respectively. Method: Chiral_GC-MS_M1.*

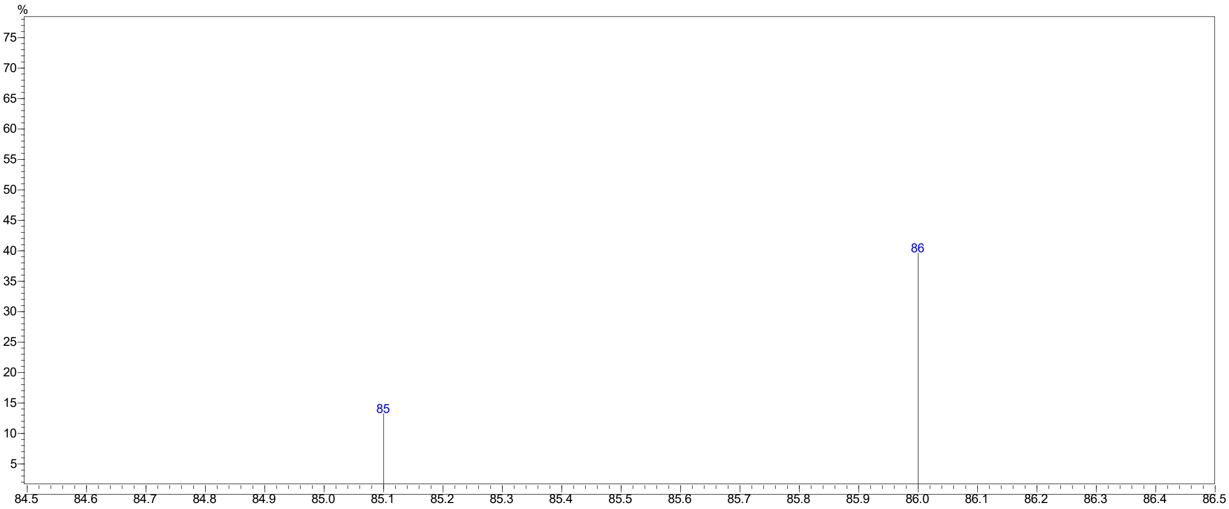

***Figure S40.*** *Mass spectrum of (S)-**4b** at 9.162 min (indicated in green), zoomed in, focusing on mass 85 and 86.*

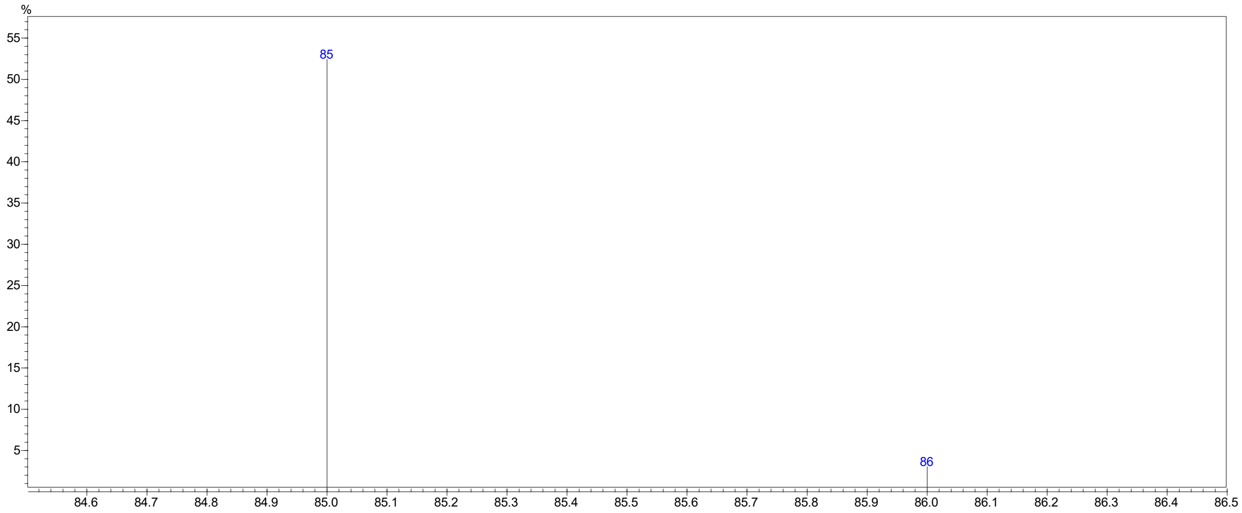

***Figure S41.*** *Mass spectrum of (R)-**4b** at 9.32 min (indicated in orange), zoomed in, focusing on mass 85 and 86.*

***Table S6.*** *Mass intensities of mass 85 and mass 86 of (S)-**4b** and (R)-**4b**.*

| | **Retention time (mass spec)** | **Intensity of mass 85** | **Intensity of mass 86** | **Percentage** |
|---|---|---|---|---|
| *(S)*-**4b*** | 9.162 min | - | - | 86>85 |
| *(R)*-**4b** | 9.32 min | 19156 | 1199 | 94.1% (of 85) |

**The peak of the (S)-enantiomer is very small. Hence, the intensities of the masses 85 and 86 of the (S)-enantiomer are not accurate. Nevertheless, mass 86 is dominant, indicating that the (S)-product results from cleavage of the pro-S carboxylate.*

*Note: due to the natural isotope abundance in carbon atoms ($^{13}C$), the peak of the (R)-enantiomer shows m/z of 86 (M+1).*

#### Decarboxylation of 4a by AMDase N131

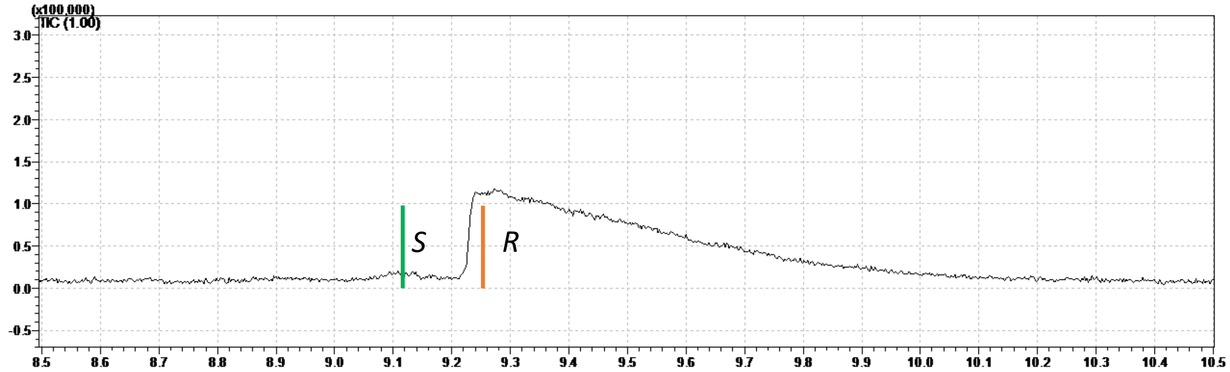

***Figure S42.*** *Chiral GC-MS chromatogram of 2-methyl-3-butenoic acid (**4b**), the product of (R)-2-methyl-2-vinylmalonic-1-$^{13}$C acid ($^{13}$C-proR-**4a**) converted by AMD N131. The green and orange bars indicate the retention times at which the mass spectra for (S)-**4b** and (R)-**4b** have been analyzed, respectively. Method: Chiral_GC-MS_M1.*

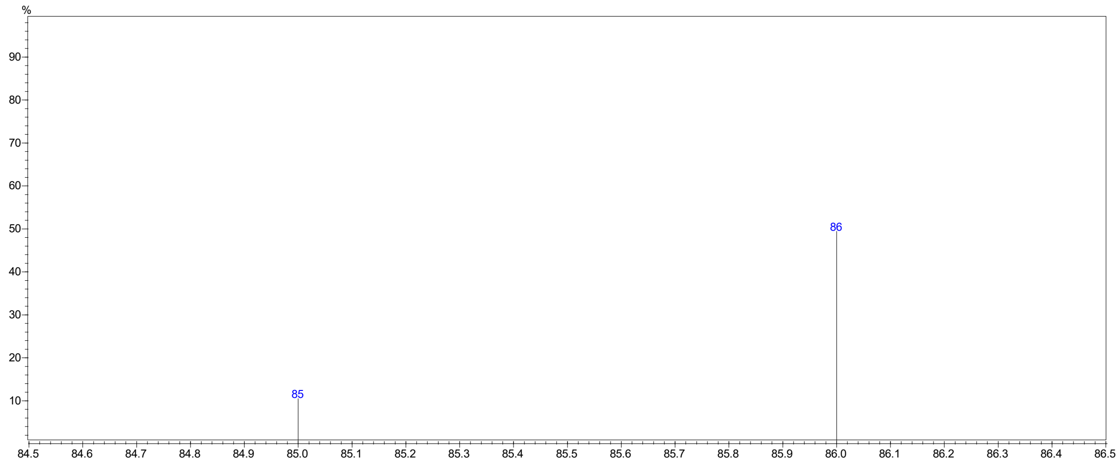

***Figure S43.*** *Mass spectrum of (S)-**4b** at 9.118 min (indicated in green), zoomed in, focusing on mass 85 and 86.*

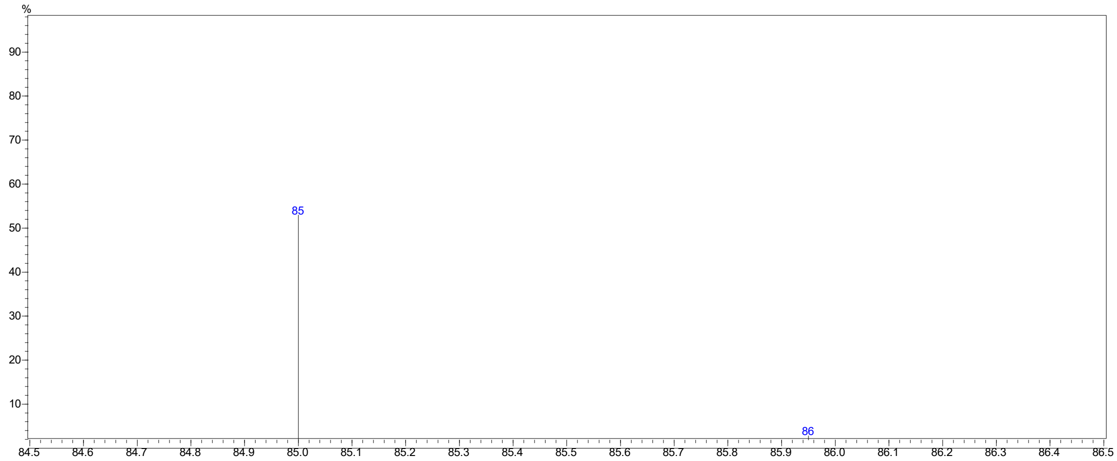

***Figure S44.*** *Mass spectrum of (R)-**4b** at 9.258 min (indicated in orange), zoomed in, focusing on mass 85 and 86.*

***Table S7.*** *Mass intensities of mass 85 and mass 86 of (S)-**4b** and (R)-**4b**.*

| | **Retention time (mass spec)** | **Intensity of mass 85** | **Intensity of mass 86** | **Percentage** |
|---|---|---|---|---|
| *(S)*-**4b*** | 9.118 min | - | - | 86>85 |
| *(R)*-**4b** | 9.258 min | 22958 | 1300 | 94.6% (of 85) |

**The peak of the (S)-enantiomer is very small. Hence, the intensities of the masses 85 and 86 of the (S)-enantiomer are not accurate. Nevertheless, mass 86 is dominant, indicating that the (S)-product results from cleavage of the pro-S carboxylate.*

*Note: due to the natural isotope abundance in carbon atoms ($^{13}C$), the peak of the (R)-enantiomer shows m/z of 86 (M+1).*

**Spontaneous decarboxylation of 4a**

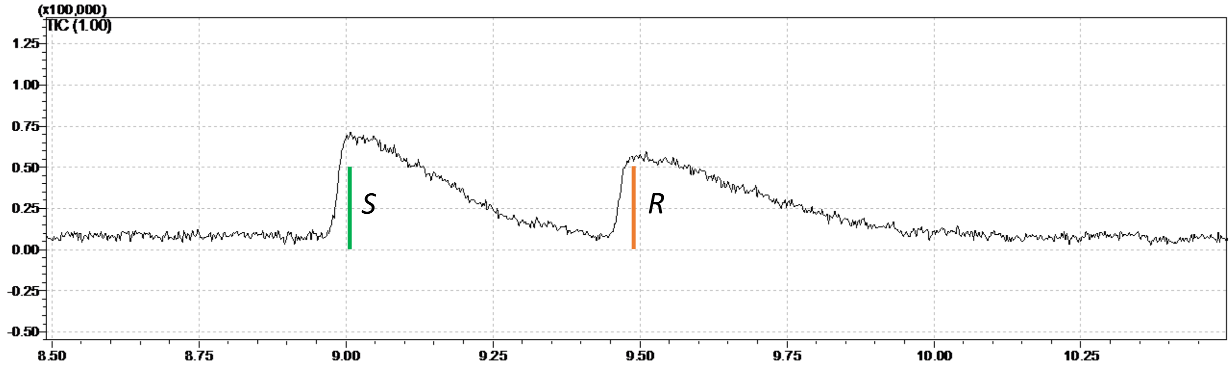

***Figure S45.*** *Chiral GC-MS chromatogram of 2-methyl-3-butenoic acid (**4b**), the product of (R)-2-methyl-2-vinylmalonic-1-$^{13}C$ acid ($^{13}C$-proR-**4a**) derived by heat-induced (60 ℃) spontaneous decarboxylation.*

*The green and orange bars indicate the retention times at which the mass spectra for (S)-**4b** and (R)-**4b** have been analyzed, respectively. Method: Chiral_GC-MS_M1.*

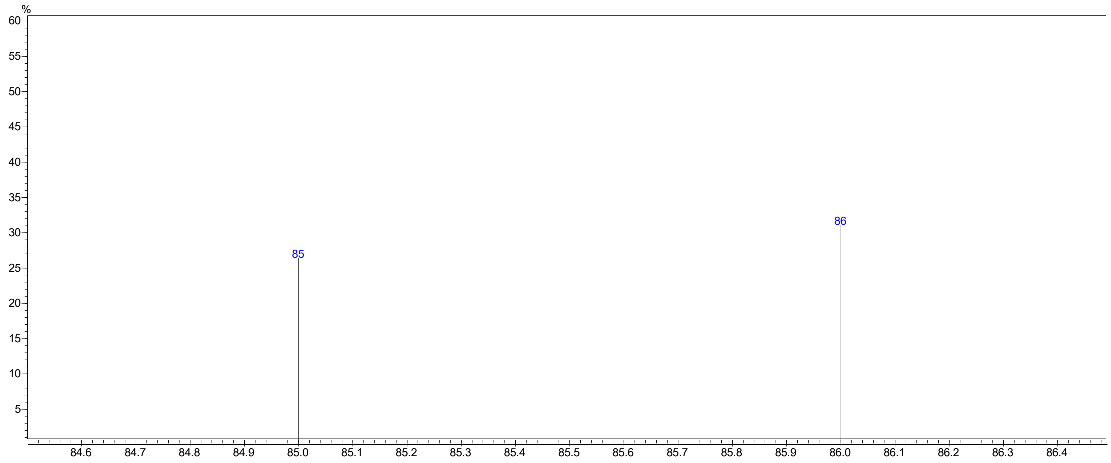

***Figure S46.*** *Mass spectrum of (S)-**4b** at 9.015 min (indicated in green), zoomed in, focusing on mass 85 and 86.*

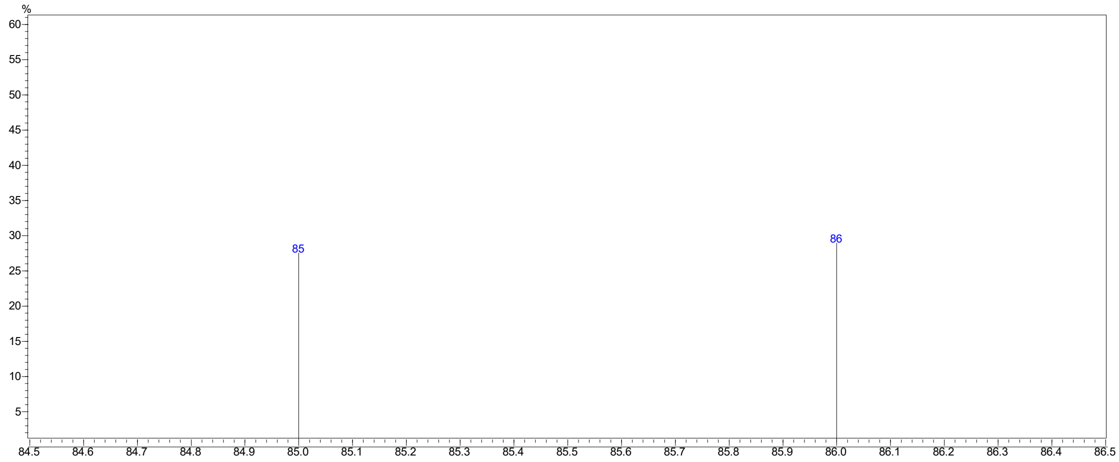

***Figure S47.*** *Mass spectrum of (R)-**4b** at 9.489 min (indicated in orange), zoomed in, focusing on mass 85 and 86.*

***Table S8.*** *Mass intensities of mass 85 and mass 86 of (S)-**4b** and (R)-**4b**.*

| | **Retention time (mass spec)** | **Intensity of mass 85** | **Intensity of mass 86** | **Percentage [%]** |
|---|---|---|---|---|
| *(S)*-**4b*** | 9.015 min | 6772 | 7719 | 53.3 (of 86) |
| *(R)*-**4b** | 9.489 min | 5735 | 6414 | 52.8 (of 86) |

#### 8. GIBBS TRANSITION STATE ENERGY CALCULATIONS

***Equation S4*** describes the relationship between the *E* value, the relative amount of (*R*)- and (*S*)-enantiomers formed during the reaction, and the ΔΔG‡. The ΔΔG‡ is the energy of the stereoselectivity, *R* is the molar gas constant (= 8.314 J·mol$^{-1}$·K$^{-1}$), and *T* is the temperature. Hence, by obtaining the *E* value (or *ee*) from a specific reaction, the ΔΔG‡ value can be calculated (***Equation S5***).[14] The energy difference between the two transition states is proportional to the temperature and the natural logarithm (ln) of the stereoselectivity. For 2-methyl-3-butenoic acid (**4b**), the Gibbs transition state energy ΔΔG‡ was calculated for *Bb*AMDase and for AMDase N31. The difference in Gibbs transition state energy ΔΔΔG‡ of *Bb*AMDase and AMDase N31 was determined by subtraction of the respective values, resulting in 1.4 kcal·mol$^{-1}$.

$$E = \frac{v_R}{v_S} = exp^{\frac{\Delta\Delta G^{\ddagger}}{RT}} \quad \text{(S4)}$$

$$\Delta\Delta G^{\ddagger} = RT \ln\left(\frac{v_R}{v_S}\right) \quad \text{(S5)}$$

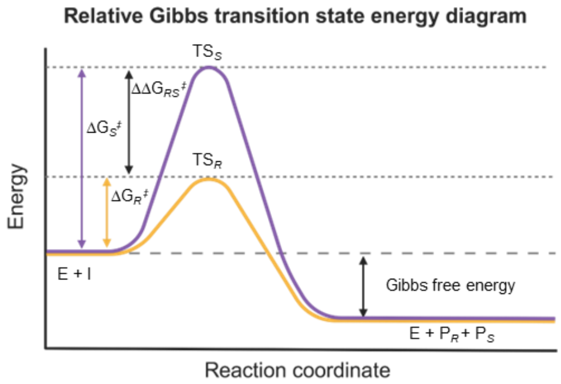

***Figure S48.*** *Relative Gibbs transition state energy diagram of a (R)-selective AMDase. The transition state energy forming the (S)-enantiomer (TS$_S$) is significantly higher than the transition state energy that produces the (R)-enantiomer (TS$_R$).*[14] *The energy differences are relative.*

#### 9. SUBSTRATE SYNTHESIS

Phenyl malonate (**1a**) and 2-methyl-2-phenyl malonic acid (**2a**) were purchased from Aaron Chemicals LLC. 2-Ethyl-2-phenyl malonic acid (**3a**) and 2-cyclohexene-1,1-dicarboxylic acid (**6a**) were synthesized according to van der Pol et al.[13]

##### 9.1 Synthesis of diethyl-2-alkyl-2-vinyl malonates

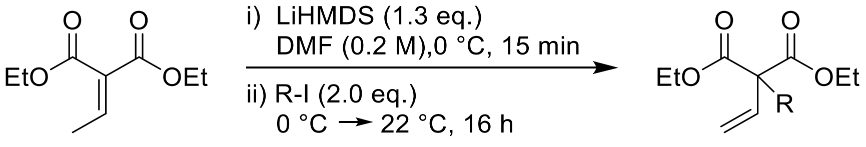

A 250 mL two-neck flask was flame-dried and put under an argon atmosphere. Diethyl 2-ethylidene malonate (5 g, 26.9 mmol, 1 eq.) was added to absolutely dry DMF (0.2 M, 134 mL) and was degassed by the freeze-pump-thaw method in two cycles[15] and cooled to 0 °C. LiHMDS (1 M in THF, 34.9 mL, 34.9 mmol, 1.3 eq.) was then added dropwise to the solution over 15 min while stirring (600 rpm). The reaction was stirred for 30 min, until the alkyl halide (methyl iodide or ethyl iodide, 0.67 mmol, 2.0 eq.) was added dropwise over 15 min and the reaction mixture was allowed to warm to 22 °C overnight. The reaction was terminated with double the volume of water and ice, and adjusted to pH 4 with HCl (1 M) and extracted with EtOAc (3 x 150 mL). The combined organic layers were washed with LiCl solution (1.0 M, 30 mL), brine (50 mL), and dried over $MgSO_4$. The crude solution was concentrated under reduced pressure and purified by column chromatography (Cyclohexane:EtOAc, 20:1) to obtain a colorless oil.
**TLC:** Ethylacetate:Cyclohexane - 1:19, $KMnO_4$-stain.

**Diethyl-2-methyl-2-vinyl malonate** (4.34 g, 21.7 mmol, 80.6%)
**$^1$H-NMR** (400 MHz, $CDCl_3$): δ(ppm) = 6.32 (dd, J=17.6, 10.7 Hz, 1H), 5.29 (dd, J=10.7, 0.5 Hz, 1H), 5.22 (dd, J=17.6, 0.5 Hz, 1H), 4.22 (q, J = 7.1 Hz, 2H), 1.58 (s, 3H), 1.28 (t, J = 7.1 Hz, 6H).

**$^{13}$C-NMR** (101 MHz, $CDCl_3$): δ(ppm) = 171.01, 136.09, 115.99, 77.34, 77.03, 76.71, 61.57, 56.19, 19.60, 14.00.

**Diethyl-2-ethyl-2-vinyl malonate** (4.67 g, 21.8 mmol, 81.2%)
**$^1$H-NMR** (400 MHz, $CDCl_3$): δ(ppm) = 6.34 (dd, J = 17.8, 10.9Hz, 1H), 5.32 (dd, J = 10.9, 0.6Hz, 1H), 5.19 (dd, J = 17.8, 0.6 Hz, 1H), 4.23 (q, J = 7.1 Hz, 2H), 2.11 (q, J = 7.5 Hz, 2H), 1.27 (t, J=7.1 Hz, 7H), 0.87 (t, J=7.5 Hz, 3H).

**$^{13}$C-NMR** (101MHz, $CDCl_3$): δ(ppm) = 170.51, 134.64, 116.60, 77.34, 77.02, 76.71, 61.35, 60.56, 27.98, 14.03, 8.62.

#### 9.2 Synthesis of 2-alkyl-2-vinyl malonic acids

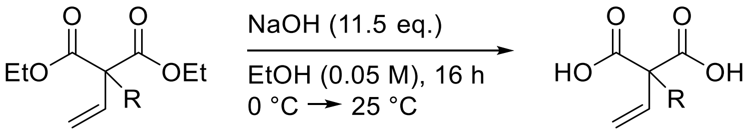

The synthesis of 2-alkyl-2-vinylmalonates was carried out under an inert atmosphere. For this, ethanol (250 mL) and aqueous NaOH (14.3 M, 10.8 mL, 11.5 eq.) were degassed by the freeze-pump-thaw method in two cycles.[14] The solution was cooled to 0 °C, and diethyl 2-alkyl 2-vinylmalonate (13.5 mmol, 1 eq., degassed) was added dropwise within 15 min. The reaction mixture was slowly warmed to room temperature overnight with stirring (600 rpm). The reaction solution was diluted with an equal volume of water and ice. Adjustment to pH 4 was done *carefully* with HCl (1 M) and extracted with MTBE (3 x 150 mL). The combined organic phases were washed with brine (200 mL) and dried over $MgSO_4$. The solvent was removed under reduced pressure at 30 °C to obtain a white, powdery solid.
**TLC**: Cyclohexane: Ethylacetate, 1:9, 1% AcOH; $KMnO_4$-stain.

**2-methyl-2-vinylmalonate** (1.74 g, 12.1mmol, 89.7%)
**$^1$H-NMR** (400 MHz, DMSO-$d_6$): δ(ppm) = 12.86 (s, 2H), 6.23 (dd, J=17.6, 10.7 Hz, 1H), 5.19 (dd, J=10.7, 0.8 Hz, 1H), 5.14 (dd, J=17.6, 0.8 Hz, 1H), 1.39 (s, 3H).

**$^{13}$C-NMR** (101MHz, DMSO-$d_6$): δ(ppm) = 172.59, 137.84, 115.38, 56.01, 19.64.

**2-ethyl-2-vinylmalonate** (1.07 g, 6.8 mmol, 62.4%)
**$^1$H-NMR** (400 MHz, DMSO-$d_6$): δ(ppm) = 12.80 (s, 1H), 6.23 (dd, J=17.9, 10.9 Hz, 1H), 5.25 (dd, J=10.9, 0.9 Hz, 1H), 5.13 (dd, J=17.9, 0.9 Hz, 1H), 1.93 (q, J=7.4 Hz, 2H), 0.78 (t, J=7.5 Hz, 3H).

**$^{13}$C-NMR** (101MHz, DMSO-$d_6$): δ(ppm) = 172.03, 136.13, 116.12, 60.36, 27.37, 9.14.

#### 10. COMPUTATIONAL ANALYSIS

***Table S9.*** *Sequence-based analysis on wild-type BbAMDase and the N131 ancestor.*[a]

| Property | *Bb*AMDase | N131 |
|---|---|---|
| $T_m$ (°C, experimental) | 55 | 62 |
| Half-life relative to wild-type | -- | 300 x |
| SCMTPP Propensity Score | 404.69 | 421.84 |
| Protein-Sol Predicted Scaled Solubility | 0.512 | 0.661 |
| Ala | 15 | 14.2 |
| Arg | 7.1 | 6.7 |
| Asn | 1.2 | 1.2 |
| Asp | 5.4 | 4.7 |
| Cys | 1.7 | 0.4 |
| Gln | 1.7 | 0.8 |
| Glu | 4.6 | 10.2 |
| Gly | 10.8 | 11 |
| His | 0.4 | 2.8 |
| Ile | 3.3 | 3.9 |
| Leu | 11.7 | 10.6 |
| Lys | 0.8 | 0.4 |
| Met | 2.5 | 2.8 |
| Phe | 2.9 | 2.8 |
| Pro | 5.8 | 5.5 |
| Ser | 6.7 | 6.3 |
| Thr | 6.2 | 5.5 |
| Trp | 0.4 | 0.4 |
| Tyr | 2.1 | 1.6 |
| Val | 9.6 | 8.3 |
| Total negatively charged residues (Asp+Glu) | 24 | 38 |
| Tot positively charged residues (Arg + Lys) | 19 | 18 |
| Instability index (II), <40 = stable, > 40 = unstable | 44.3 | 42.2 |
| Aliphatic index | 101.3 | 95.0 |
| GRAVY score | 0.355 | 0.08 |

[a] Sequence composition analysis was performed using the ProtParam tool on the ExPasy webserver.[16] Protein thermostability was predicted from sequence based on propensity scores calculated using the scoring-card method based on the SCMTPP webserver.[17] Scaled solubility was predicted from sequence using Protein-Sol.[18] Properties that show larger deviations between the two variants are highlighted in blue.

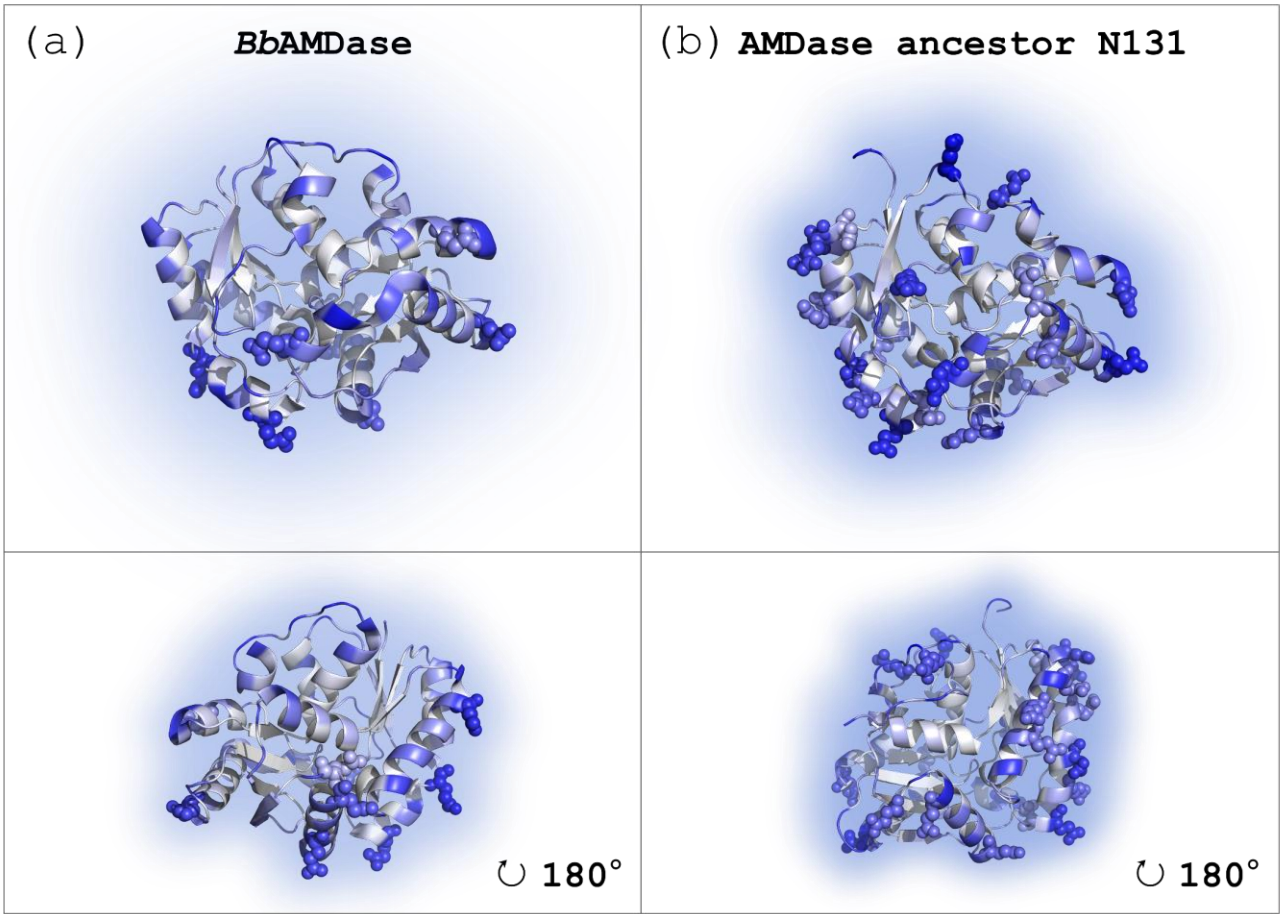

***Figure S49.*** *Depiction of BbAMDase (PDB ID: 3DG9) and the predicted ancestral AMDase N131 structures surrounded by a water shell. Glu side chains with a relative solvent-accessible surface area (SASA) above 0.25, calculated using PyMOL, are highlighted. Increasing color intensity indicates increasing solvent accessibility.*

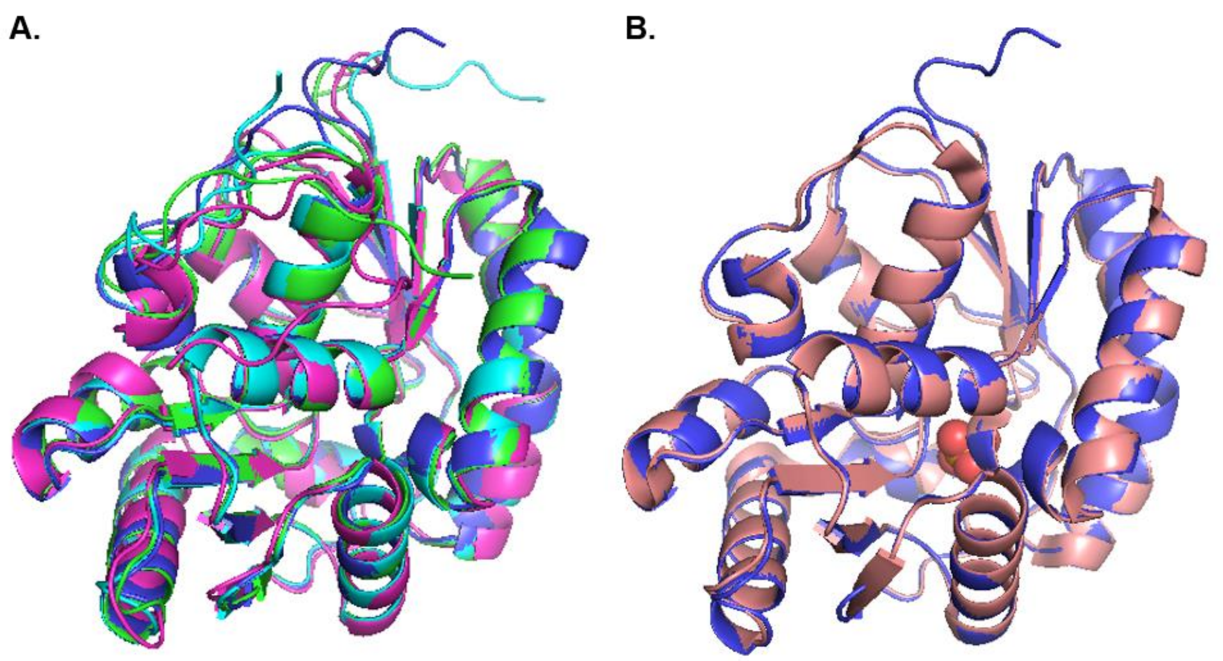

***Figure S50.*** *(**A**) Overlay of predicted structures by Chai of BbAMDase (purple), N5 (blue), N31 (green), and N131 (pink). The Overlay was visualized by PyMOL.(**B**) Overlay of the Chai-predicted structure of BbAMDase (purple) and the crystal structure of BbAMDase (light pink, PDB ID 3DG9).*

***Table S10.*** *Calculated Solvent Accessible Surface Area (SASA) for wild-type (WT) BbAMDase and the N131 ancestor, from simulations run at 300 and 360 K.[a]*

| **SASA** | **WT (300 K)** | **WT (360 K)** | **N131 (300 K)** | **N131 (360 K)** |
|---|---|---|---|---|
| Total (Å$^2$) | 11575.4 ± 364.8 | 12084.5 ± 409.3 | 12858.6 ± 522.4 | 13195.5 ± 533.7 |
| Hydrophobic (Å$^2$) | 5410.8 ± 217.1 | 5685.9 ± 254.9 | 5654.6 ± 315.5 | 5804.3 ± 340.3 |
| Hydrophilic (Å$^2$) | 6164.6 ± 200.8 | 6398.6 ± 205.9 | 7204.0 ± 252.4 | 7391.2 ± 262.3 |

[a] Shown here are average values and standard deviations calculated over 10 x 0.5 μs of simulation time per system.

***Table S11.*** *Calculated number of interactions by interaction type for wild-type (WT) BbAMDase and the N131 ancestor, from simulations run at 300 and 360 K.[a]*

| **Interaction type** | **WT (300 K)** | **WT (360 K)** | **N131 (300 K)** | **N131 (360 K)** |
|---|---|---|---|---|
| Salt bridges | 18.1 ± 3.8 | 16.7 ± 4.2 | 16.0 ± 4.5 | 18.9 ± 4.7 |
| Hydrophobic contacts | 885.5 ± 18.2 | 861.4 ± 20.0 | 896.6 ± 19.2 | 873.3 ± 20.4 |
| H-bond (intramolecular) | 100.0 ± 7.7 | 89.2 ± 7.6 | 101.3 ± 7.4 | 94.5 ± 7.9 |

[a] Shown here are average values and standard deviations calculated over 10 x 0.5 μs of simulation time per system.

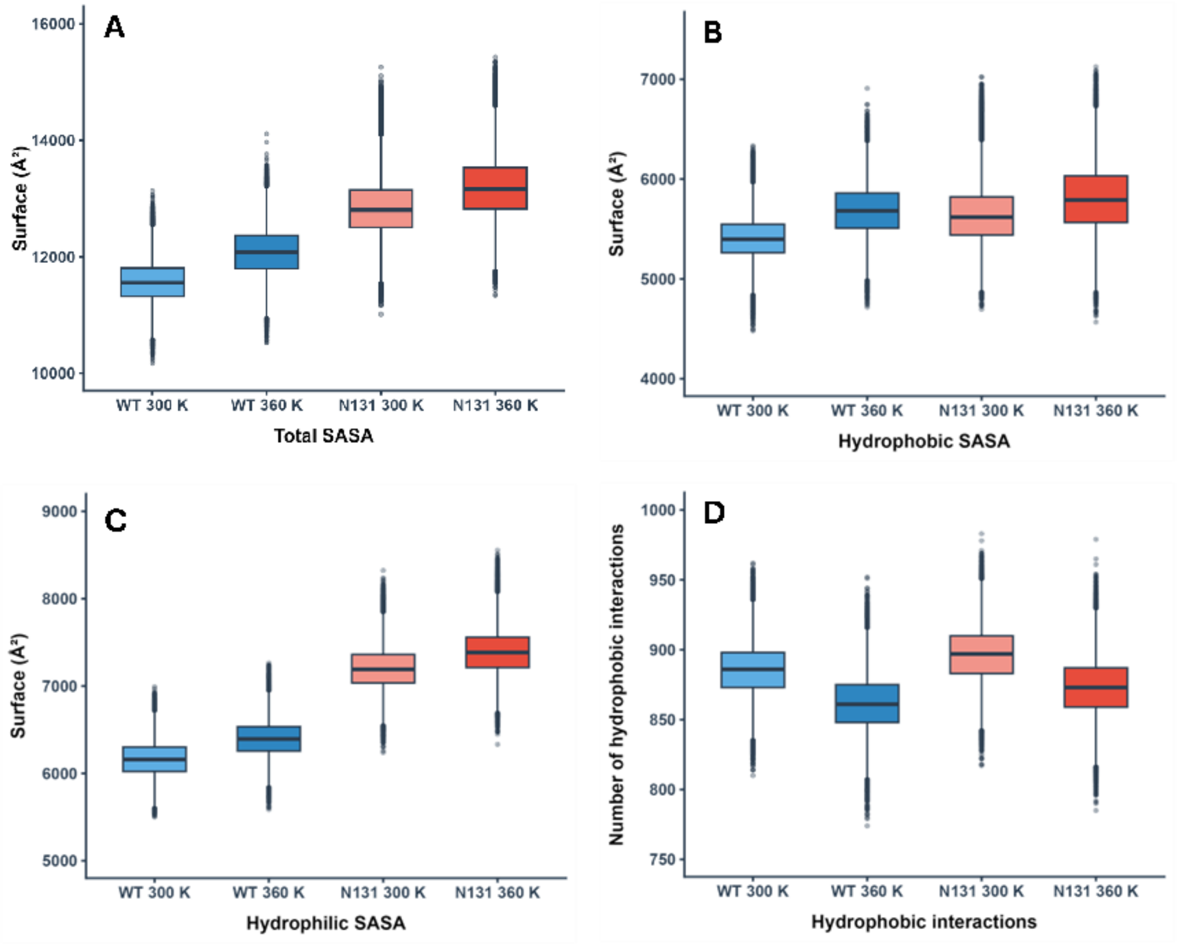

***Figure S51.*** *Breakdown of the (**A**) total, (**B**) hydrophobic, (**C**) hydrophilic and (**D**) hydrophobic ratio of the solvent accessible surface areas (SASA, Å2) of wild-type BbAMDase and the N131 ancestor, calculated from 10 x 0.5 µs simulations of each system at 300 and at 360 K, respectively. SASA values were calculated as described in the **Materials and Methods**. Raw data for this figure is shown in*

*Table S10*.

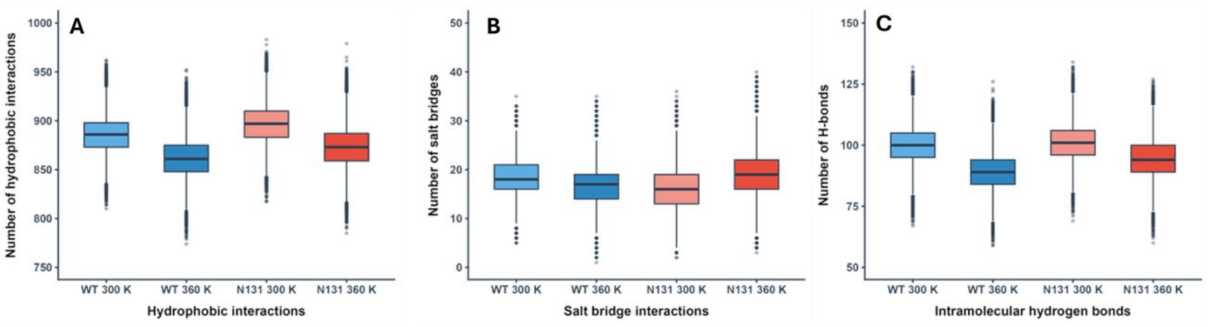

***Figure S52.*** *Average numbers of (**A**) hydrophobic interactions, (**B**) salt bridges and (**C**) intramolecular hydrogen bonds, calculated across 10 x 0.5 µs simulations of wild-type BbAMDase and the N131 ancestor, at each of 300 and 360 K, respectively. Analysis was performed as described in the **Materials and Methods**. Raw data for this figure is shown in **Table S11**.*

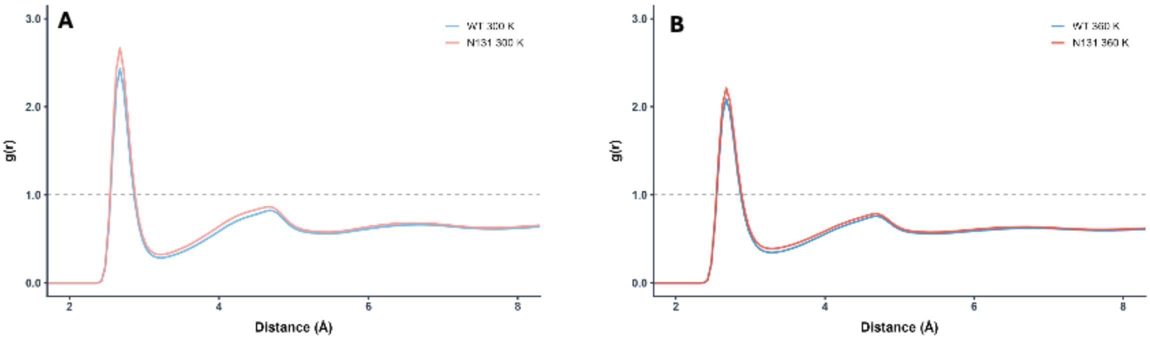

***Figure S53.*** *Radial distribution functions of water molecules surrounding the terminal oxygen atoms of acidic residues (aspartic and glutamic acid residues), based on 10 x 0.5 µs simulations of (blue) wild-type BbAMDase and (red) the N131 ancestor, at (**A**) 300 and **(B**) 360 K. The functions were derived with CPPTRAJ,[19] with a resolution of 0.05 Å, and are depicted as a function of distance from the terminal oxygen atoms.*

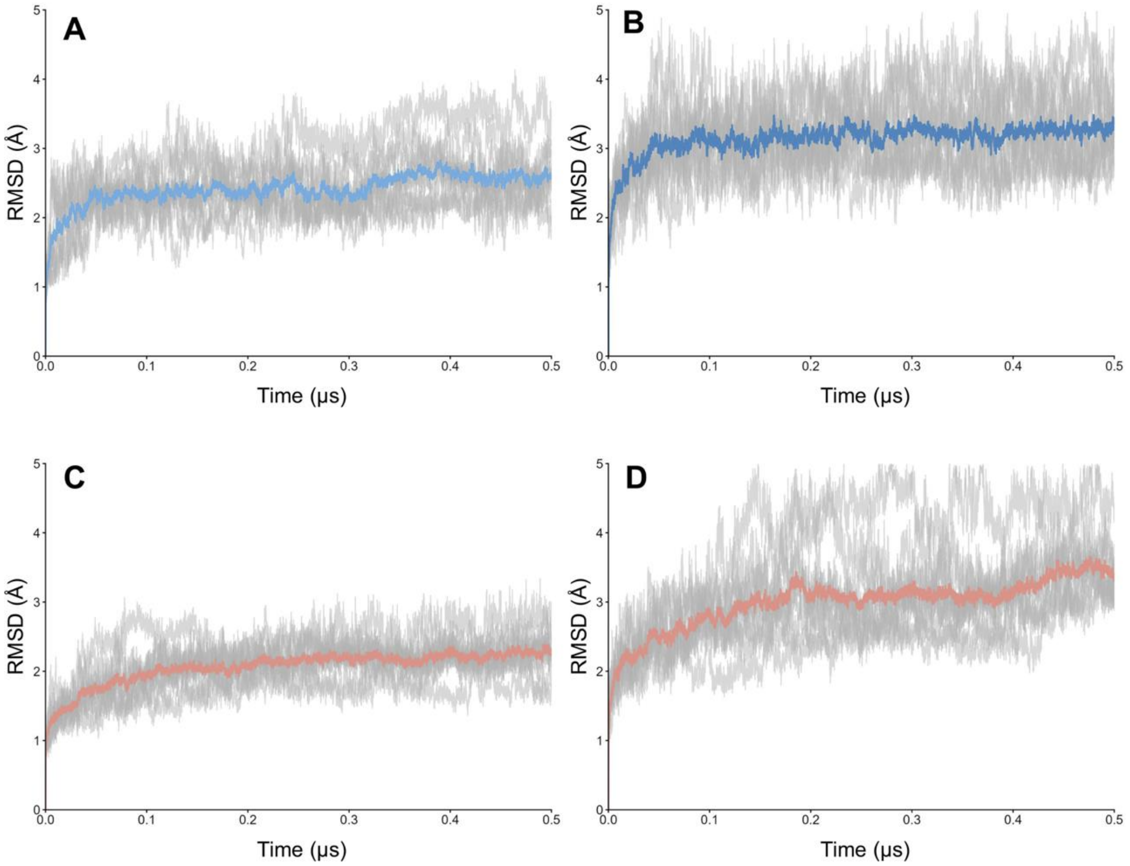

***Figure S54.*** *Root mean square deviation (RMSD, Å) of all Cα-atoms of wild-type BbAMDase and the N131 ancestor, as a function of simulation time, shown over 10 x 0.5 µs simulations of each system. Shown here is analysis from simulations of (**A**, **B**) wild-type and (**C**, **D**) N131 AMDase, with simulations run at (**A**, **C**) 300 and (**B**, **D**) 360 K, respectively. The shaded area shows data from individual replicates, and the solid line shows the average RMSD across all trajectories.*

#### 11. NMR SPECTRA

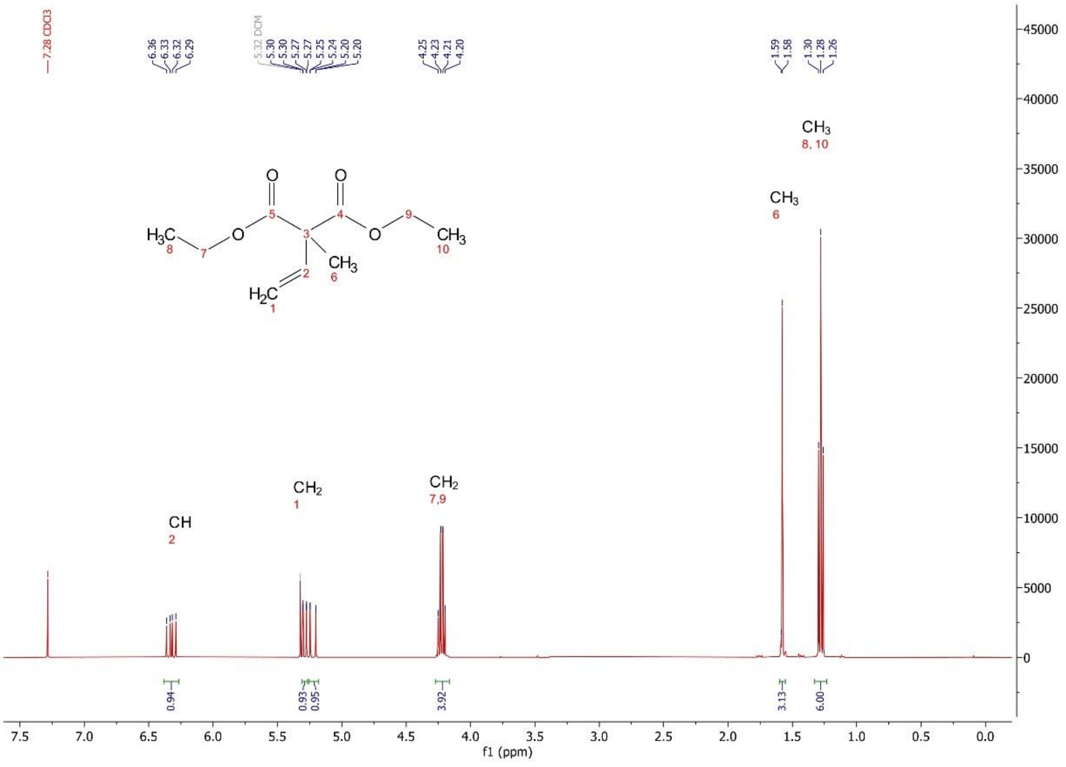

***Figure S55.*** *Measured $^{1}$H-NMR Spectrum of diethyl-2-methyl-2-vinyl malonate in $CDCl_3$.*

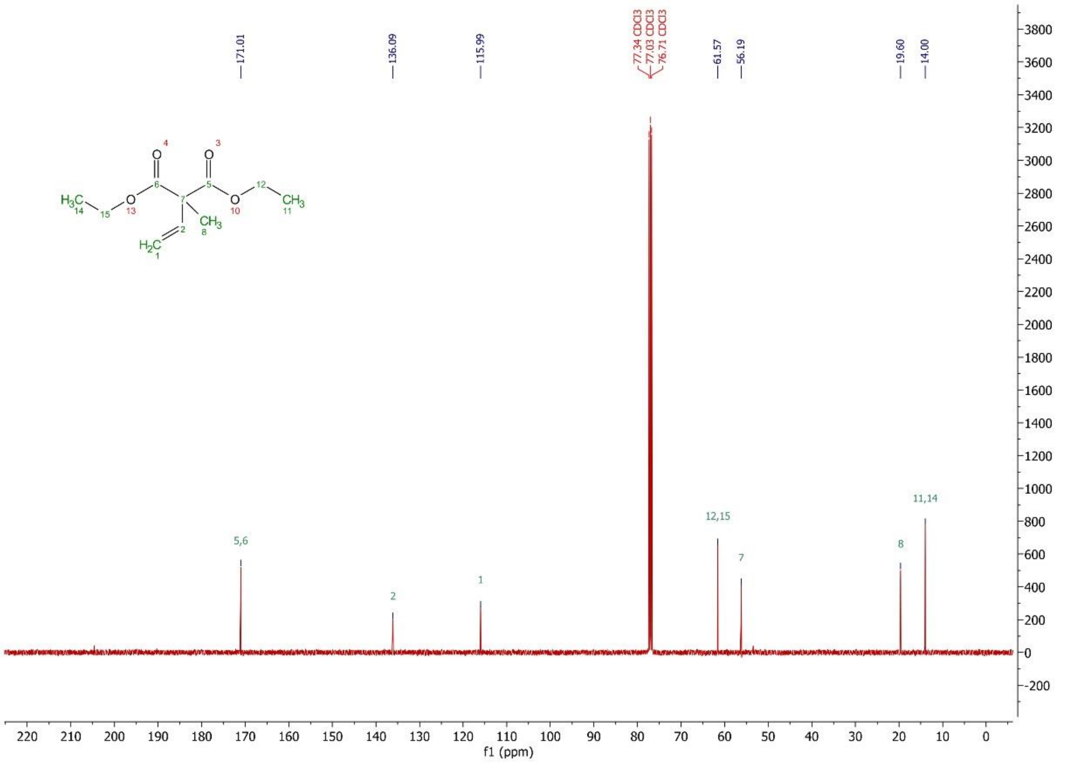

***Figure S56.*** *Measured $^{13}$C-NMR Spectrum of diethyl-2-methyl-2-vinyl malonate in $CDCl_3$.*

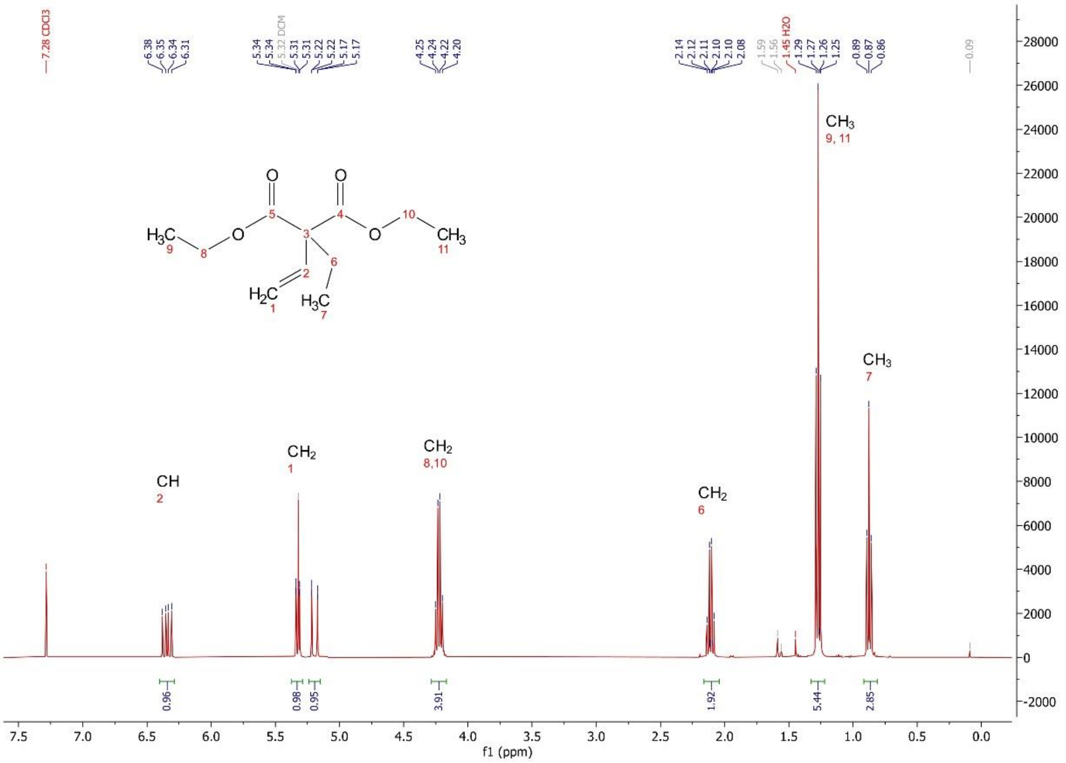

***Figure S57.*** *Measured [1]H-NMR Spectrum of diethyl-2-methyl-2-vinyl malonate in $CDCl_3$.*

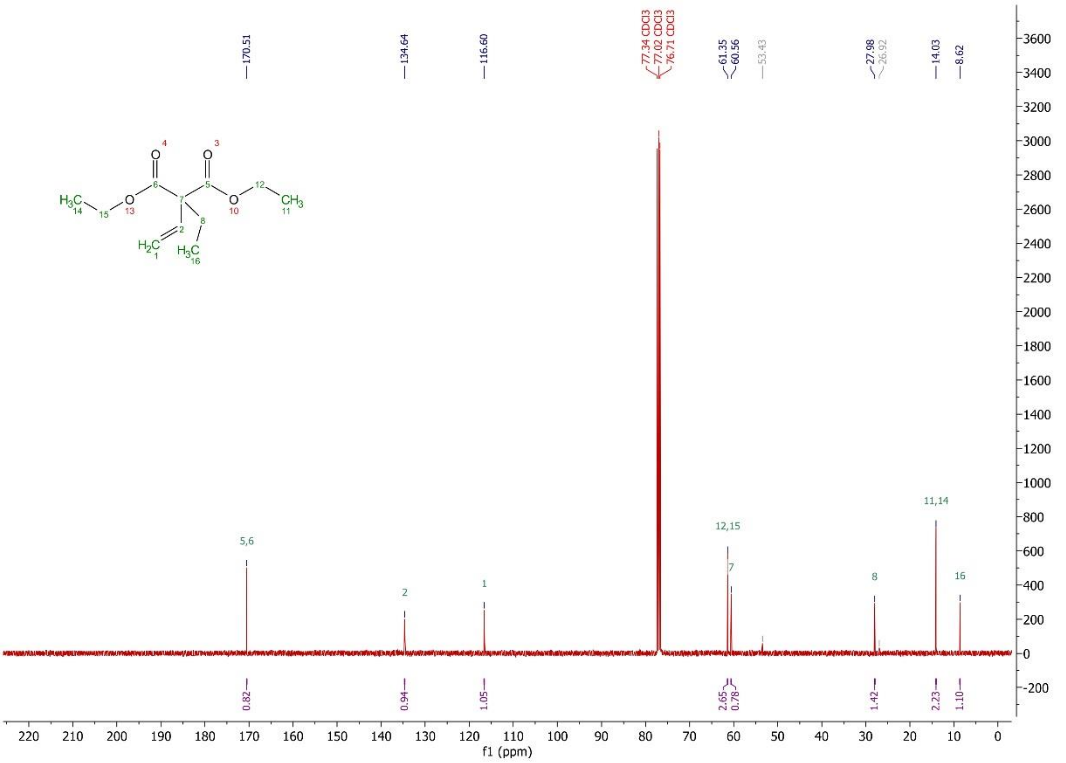

***Figure S58.*** *Measured [13]C-NMR Spectrum of diethyl-2-methyl-2-vinyl malonate in $CDCl_3$.*

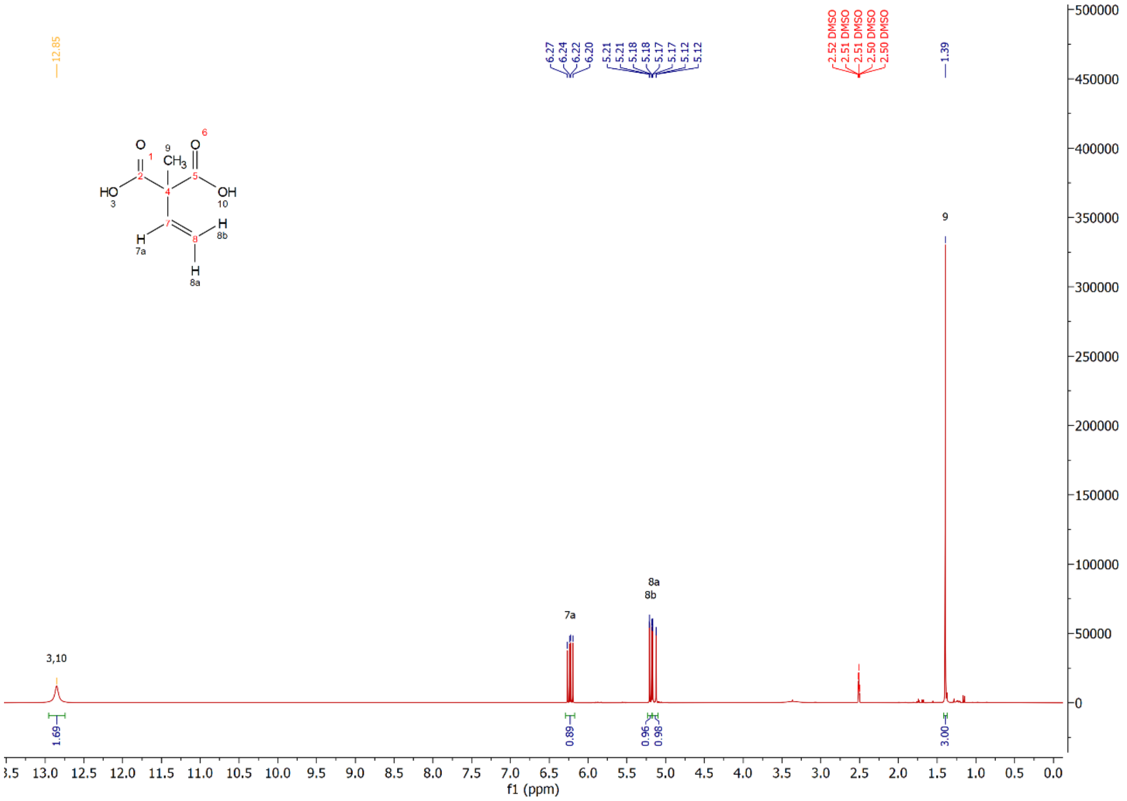

***Figure S59.*** *Measured $^{1}$H-NMR Spectrum of 2-methyl-2-vinylmalonate in DMSO-$d_6$.*

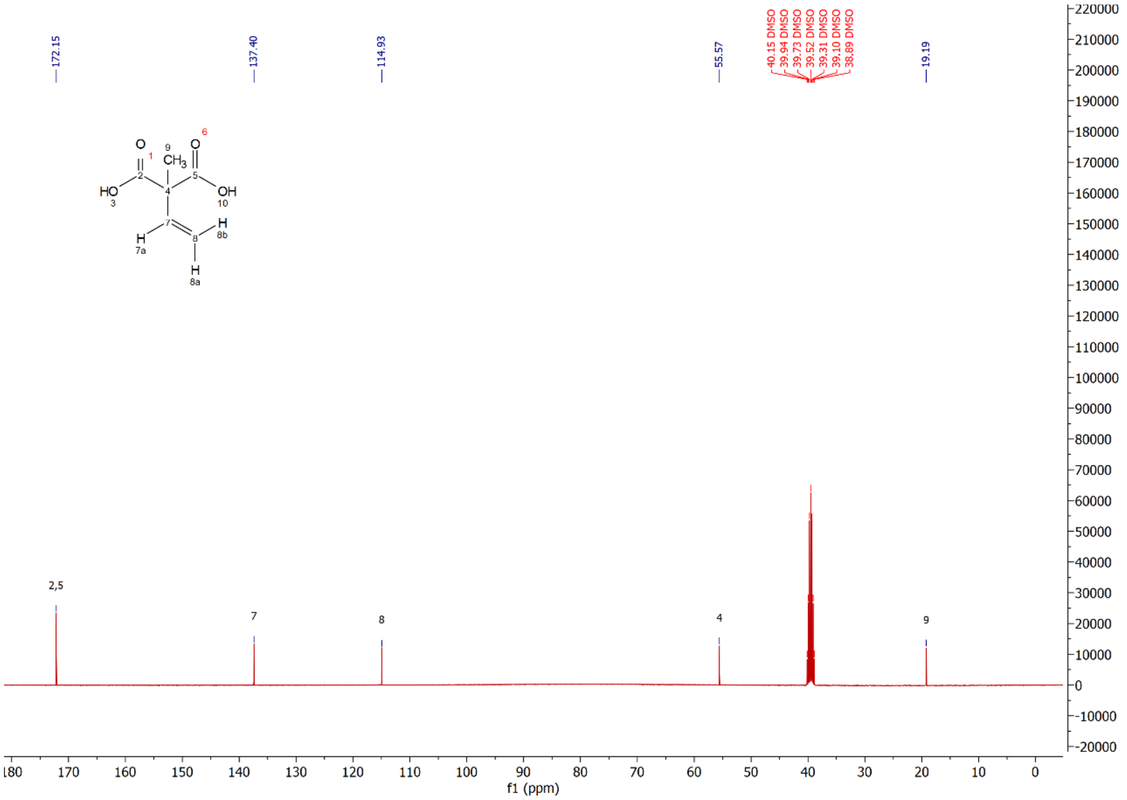

***Figure S60.*** *Measured $^{13}$C-NMR Spectrum of 2-methyl-2-vinylmalonate in DMSO-$d_6$.*

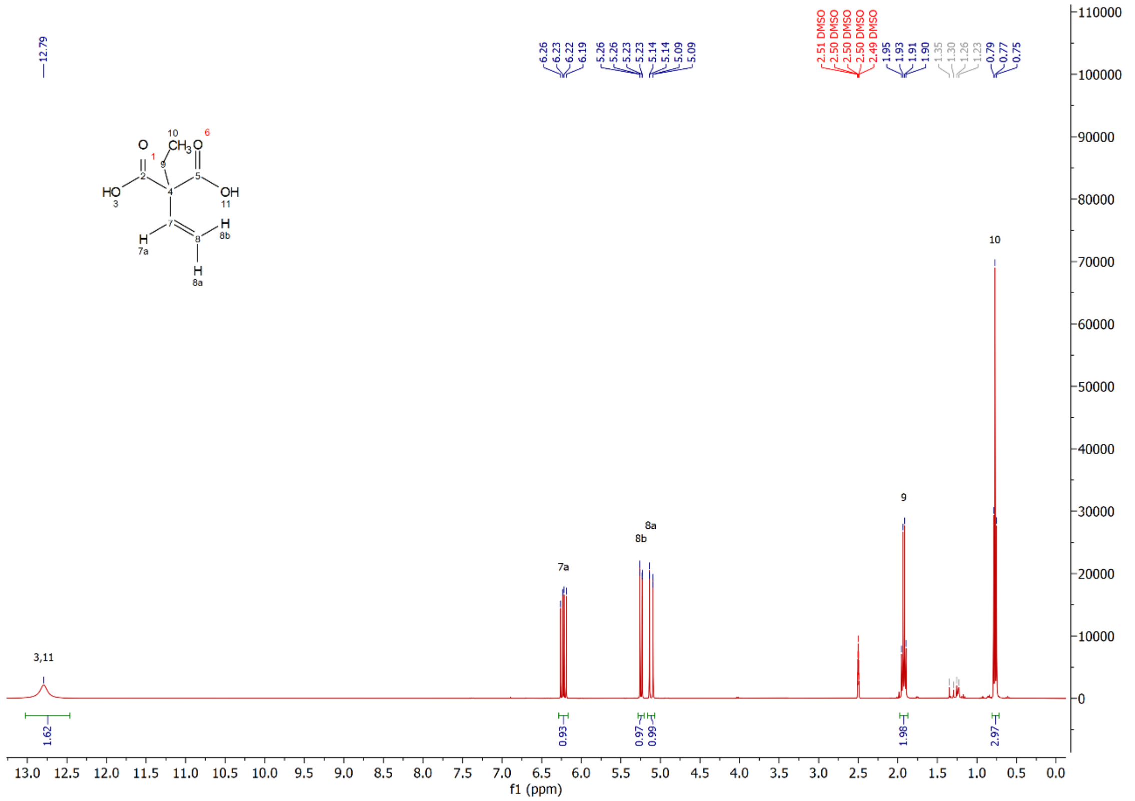

***Figure S61.*** *Measured ¹H-NMR Spectrum of 2-ethyl-2-vinylmalonate in DMSO-$d_6$.*

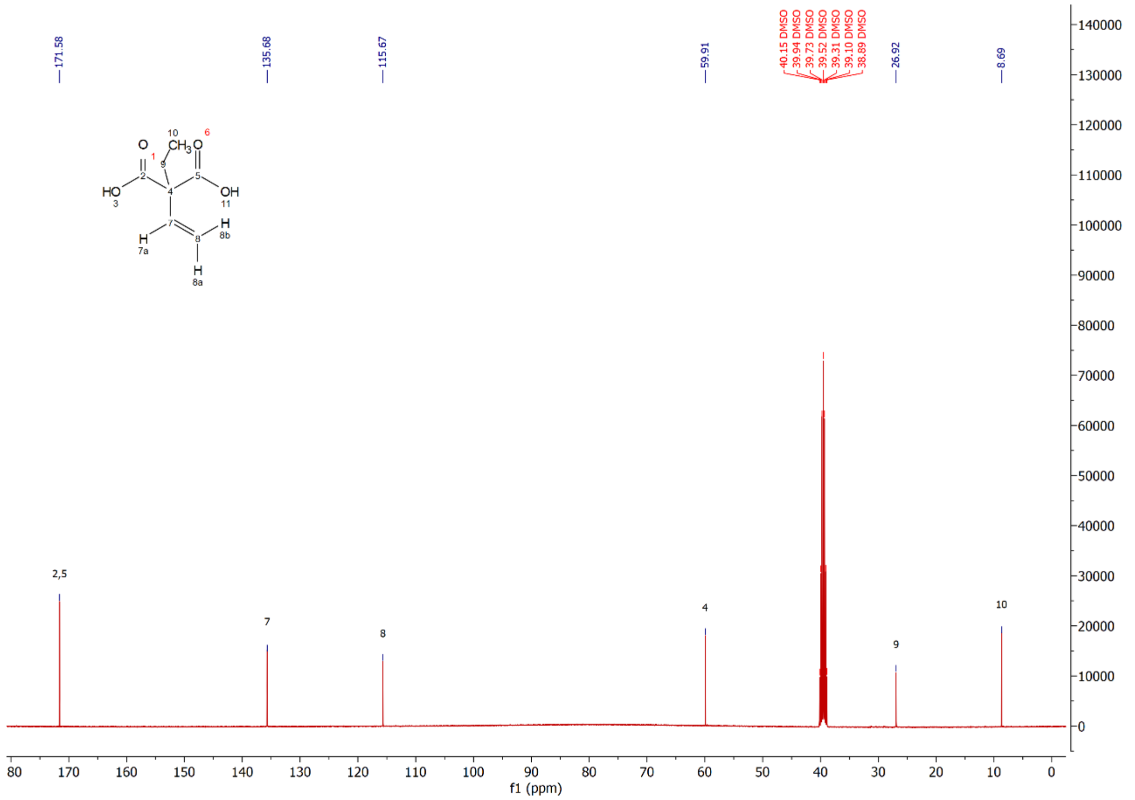

***Figure S62***. *Measured ¹³C-NMR Spectrum of 2-ethyl-2-vinyl malonate in DMSO-$d_6$.*
